# Soil microbiome structure and function reflect environmental variation rather than reindeer presence in a northern peatland

**DOI:** 10.64898/2026.05.13.724277

**Authors:** Tommi Välikangas, Hannu Fritze, Juha-Matti Pitkänen, Krista Peltoniemi, Eeva Järvi-Laturi, Torben R. Christensen, Maria Väisänen, Juho Lämsä, Riku Paavola, Jenni Hultman

## Abstract

Northern peatlands store large carbon stocks but are sensitive to disturbance. Hydrology, vegetation, herbivory and snow conditions may affect the soil microorganisms driving methane (CH₄) and nitrous oxide (N₂O) cycling. We investigated how reindeer exclusion and snow depth (increased and reduced relative to ambient) manipulations (ongoing for three seasons) influenced archaeal and bacterial communities in a boreal rich fen. Metagenomic (MG) and metatranscriptomic (MT) sequencing were combined with pore-water chemistry and CH₄ flux measurements to link the microbiome to ecosystem processes.

Microbial communities differed between outside and inside the exclosure. However, these patterns primarily reflected underlying hydrological variation. Slightly wetter inside plots showed higher expression of denitrification genes (*norB*, *nosZ*) and lower (*nirS*+*nirK*)/*nosZ* ratios, indicating greater potential for complete denitrification to N₂ instead of N₂O. Methane dynamics were mainly associated with vegetation: plots associated with *Carex rostrata* exhibited lower *pmoA*/*mcrA* ratios and elevated CH₄ fluxes. Snow manipulations had subtle effects: reduced snow depth decreased the expression of taxa dependent on microbial interactions, while the effect to the investigated metabolic marker genes was small.

Overall hydrology, leading to variations in redox conditions and nutrient availability, together with vegetation appeared as the primary drivers on microbial greenhouse gas processes in this peatland.

## Introduction

The rate of global warming has been observed to be pronounced in the northern hemisphere compared to the global average with recent estimates predicting the temperature in the Arctic rising several times faster than the rest of the globe (Rantanen *et al*. 2022, Xie *et al*. 2022). Northern peatlands are major carbon storages, estimated to hold ∼80% of the world’s total peatland carbon and nitrogen, and have an essential role in global carbon cycling (Hugelius *et al*. 2020, Harris *et al*. 2022). Historically, these peatlands have acted as carbon dioxide (CO_2_) sinks (Limpens *et al*. 2008) as well as methane (CH_4_) sources (Blodau 2002). However, their contribution to greenhouse gas (GHG) emissions, such as CO_2_, CH_4_, and nitrous oxide (N_2_O), are expected to increase in the future due to global warming (Schuur *et al*. 2015, Voigt *et al*. 2017, Hugelius *et al*. 2020, Turetsky *et al*. 2020). The biogeochemical processes in these northern peat soils, also those influencing atmospheric GHG concentrations, are largely controlled by microbial activity (Falkowski, Fenchel, and Delong 2008). Microbial reactions steer the biological flow of the major elements, namely hydrogen, carbon, nitrogen, oxygen and sulfur (Falkowski, Fenchel, and Delong 2008) and microbial processes in the soil, such as denitrification and methanogenesis are major sources of atmospheric N₂O (Baggs 2011) and CH_4_ (Falkowski, Fenchel, and Delong 2008).

The source, flow and storage of water (hydrology), is the main determinator of peatland structure and function, driving geochemical and biotic processes alike. However, water table levels and pore water chemistry may vary both temporally and spatially within both bogs and fens (Tahvanainen *et al*. 2002, Griffiths, Sebestyen, and Oleheiser 2019, Krishnankutty *et al*. 2025). The soil microbiome in peatlands also reflects hydrological and chemical regimes. Both the peat soil microbial community composition and its metabolic activity are connected to hydrological and geochemical gradients, especially in minerotrophic fens where mineral-rich water inputs raise the pH, alkalinity and base cation concentrations, supporting higher microbial diversity and enzymatic activity (Lin *et al*. 2012, Andersen, Chapman, and Artz 2013, Urbanová and Bárta 2014, Seward *et al*. 2020). Microbial communities can, in turn, affect the peat-accumulation and carbon dynamics of peatlands through for example carbon decomposition, CH_4_ production and carbon export (Schimel and Schaeffer 2012, Richy *et al*. 2024).

Northern peatlands are affected by large mammalian herbivores such as reindeer or caribou (*Rangifer tarandus*) that use peatlands as their grazing grounds mainly during spring and summer (Kolari *et al*. 2019). Reindeer influence the ecosystems they are found in through selective grazing of the vegetation, trampling and fertilization via their droppings and urine (Suominen and Olofsson 2000, Sørensen *et al*. 2009, Barthelemy *et al*. 2018, Santalahti *et al*. 2018, Laiho *et al*. 2024). In addition to aboveground effects, grazing may also impact belowground properties, such as soil respiration, carbon to nitrogen ratio and soil microbial communities (Andriuzzi and Wall 2017, Santalahti *et al*. 2018, Ylänne *et al*. 2021, Stark *et al*. 2023, Laiho *et al*. 2024). In Finland, reindeer are semi-domesticated, roaming freely mainly in Finnish Lapland within a region equivalent to one-third of Finland, with numbers estimated approximately 187.000–200.000 in 2000–2001 and the maximum allowed number of reindeers for 2020–2030 set to 203700 (Suominen and Olofsson 2000, Susiluoto *et al*. 2008, Maa-ja metsätalousministeriö 2018, Santalahti *et al*. 2018). Laboratory incubations have associated reindeer droppings with increased CH_4_ production in peat soil which may be caused by methanogenic reindeer rumen microbes (Laiho, Penttilä, and Fritze 2017, Fritze *et al*. 2021). However, in field studies, reindeer presence and dropping addition treatments, while influencing the activity of soil microbial communities, did not directly affect the measured CH_4_ fluxes (Laiho *et al*. 2024). In addition, the droppings of Svalbard reindeer (*Rangifer tarandus platyrhynchus*) have been associated with a small and short initial increase in N_2_O emissions in mineral soil (Hayashi *et al*. 2014).

Historically, the soils in northern latitudes have been covered in snow throughout large parts of the year. However, global warming is changing the snow conditions by decreasing snow cover duration and snow depths, especially during spring time (Luomaranta, Aalto, and Jylhä 2019, Richardson *et al*. 2024). During the snow covered period, snow acts as an insulator, affecting soil temperatures and the soil freezing and thawing dynamics (Richardson *et al*. 2024). On the other hand, during periods of decreased snow cover, the soil is exposed to more frequent and intense freeze-thaw cycles, resulting in the potential physical breakdown of soil aggregates and altered microbial activity (Ruan and Robertson 2017). Increased N_2_O emissions from agricultural soil have been connected to decreased snow cover (Ruan and Robertson 2017). In addition, the depth of snow cover significantly affects the moisture in peat soils with snow melt during spring-time altering the water table level and the hydrological conditions of the peatlands (Metcalfe and Buttle 2001, Ketcheson, Whittington, and Price 2012), which can in turn significantly alter the microbial community structure (Jaatinen *et al*. 2007, Urbanová, Picek, and Bárta 2011). Consequently, snow cover can modify the peatland soil microbial community structure as well as the activity of the microbial metabolic pathways through regulating soil temperatures, freeze-thaw cycle dynamics and soil moisture.

To investigate how large herbivores and environmental conditions influence below-ground processes in northern peatlands, we conducted a reindeer-exclusion experiment combined with experimental manipulation of snow depth in a boreal rich fen in northeastern Finland. We compared soil microbiomes outside and inside the exclosure and across snow treatments (increased and reduced relative to ambient) three growing seasons after the start of the experiments. Using parallel metagenomic (MG) and metatranscriptomic (MT) sequencing, we assessed the impacts of the treatments to microbial community composition, functional potential and activity.

We hypothesized that (1) reindeer exclusion alters microbial community structure and metabolic functionality, particularly key microbial pathways involved in methane (CH₄) and nitrous oxide (N₂O) cycling and (2) altered snow depth modifies microbial functional community composition and these pathways by changing winter soil conditions (e.g. insulation). Because the experimental plots are located along a mild hydrological gradient in the fen, we further hypothesized (3) that variation in soil moisture and associated redox conditions interacts with the treatments to influence microbial responses. Specifically, we expected that slightly wetter and resulting more reduced conditions would favor complete denitrification processes.

Although vegetation composition was broadly similar across the study area, finer-scale differences in plant community structure were accounted for in the analyses and evaluated in relation to key methane- and nitrogen-related processes. By combining multi-omic data with pore-water chemistry and methane flux measurements, this study aimed to link microbial community structure and metabolic activity to ecosystem-level processes in a northern peatland.

## Methods

### Study site and experimental data

The study took place in Puukkosuo (**Figure 1A**). Puukkosuo is located in northeastern Finland (66.377299° N, 29.308062° E), at the national park of Oulanka in Kuusamo within the northern boreal zone. Puukkosuo is an open, minerotrophic fen with a mild (1.66°) slope in NW to SE direction. The vegetation in Puukkosuo consists mainly of vascular plants (e.g. *Carex* spp., *Trichophorum* spp., *Molinia caerulea*, *Potentilla erecta*, *Menyanthes trifoliata*) and brown and peat mosses (e.g. *Scorpidium cossonii*, *Campylium stellatum*, *Cinclidium stygium*, *Sphagnum* spp.) (Järvi-Laturi *et al*. 2025).

**Figure 1.**
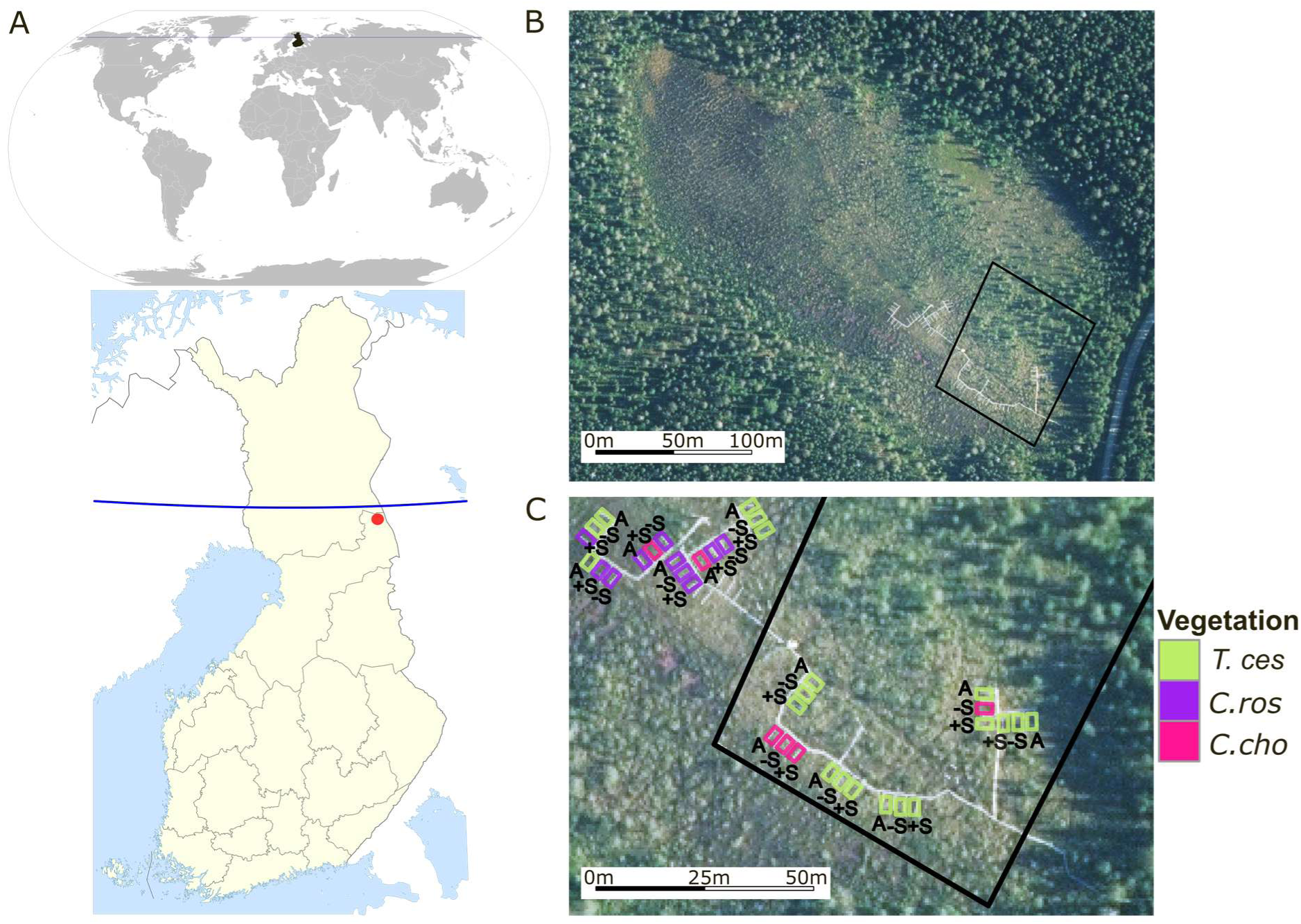
Location and experimental design at the Puukkosuo rich fen in Oulanka, northeastern Finland. **A)** The location of Finland within the world and Puukkosuo (red dot) in Finland. The blue line represents the Arctic Circle. **B)** An aerial view of the exclusion fence (black outline) and boardwalk (grey). North is up. **C)** A close-up of the experimental plots showing positions relative to the exclusion fence and boardwalk. Plot colors indicate dominant vegetation clusters: *T.ces* = *Trichophorum cespitosum*, *C.ros* = *Carex rostrata*, *C.cho* = *Carex chordorrhiza*. Abbreviations for the snow manipulaton treatments, A = ambient snow cover (control), -S = snow removal, +S = snow addition. The world base map in A and the Finland map are adapted from [Wikimedia Commons]. Aerial imagery for B and C are provided by the National Land Survey of Finland. For a detailed license description, see Supplementary Methods.

The study area encompassed approximately one hectare and was divided into 12 spatial blocks, each containing three study plots (36 plots in total). Plots were established in summer 2018 and measured 2 m × 3.5 m, including a 0.5 m-wide buffer zone. Wooden boardwalks were constructed to access the plots and minimize disturbance to the peat surface. In spring 2019, a fence was erected around half of the plots (n = 18) to exclude reindeer presence (**Figure 1B-C**). At the time of this study (September 2021), the reindeer exclusion had been in place for nearly 2 and a half years covering three growing seasons. Outside the exclusion fence, reindeer were present at natural levels to the area. The fence and the plots were adjacently aligned in NW-SE direction (**Figure 1B-C**, **Supplementary Figure 1**) with the exclusion fence situated slightly downslope and downstream from the open upslope area accessible to reindeer. The exclusion treatment partly coincided with differences in vegetation composition (see text below and (Järvi-Laturi *et al*. 2025)). For clarity, we treat the exclusion treatment contrast as combined hydrological effect and reindeer exclusion (outside vs. inside), while noting that vegetation composition may also influence the observed responses.

The plots were also subjected to snow manipulation treatments that were initiated in January 2019. Within each block, one plot served as an untreated control (AMB) with ambient snow depth, one as a snow removal treatment (-S) where snow depth was maintained at 0.2-0.25 m throughout the snow season and one as a snow addition (+S) treatment receiving snow transferred from the removal plot. Snow treatments were started once ambient snow was 0.3 m deep. At each plot, snow depth was measured regularly at three spots to calculate average depth per plot. March is the time of maximum snow depth and the average for snow depth in March 2021 in each plot by the exclusion and snow treatments are shown in **Supplementary Figure 2A-B**.

### Plant community data and vegetation clusters

We defined vascular plant community clusters using hierarchical cluster analysis on plant biomass estimates. Clustering was based on Sorensen (Bray–Curtis) distance measure and Flexible Beta group linkage method (β = -0.25). Clusters with less than six plots were excluded. To evaluate which species exhibited the highest statistical connectivity to the different clusters, we carried out an indicator species analysis. This showed that clusters were characterized mostly by different sedge species: *Carex rostrata* (*C.ros*), *C. chordorrhiza*, (*C.cho*) and *Trichophorum cespitosum* (*T.ces*) (**Figure 1C**). For a more detailed description of the plant community analyses, see (Järvi-Laturi *et al*. 2025).

### Peat chemistry and methane flux measurements

Peat pore-water was collected five times from May to September 2021 (Rhizon samplers, Rhizosphere Research Products, Netherlands) at 10 cm depth into evacuated opaque syringes, after which samples were filtered (0.45 μm sterile nylon, Sarstedt, Germany) and frozen at −18 °C. Thawed samples were analyzed as in (Järvi-Laturi *et al*. 2025) for pH (913 pH/DO Meter, Metrohm, Switzerland), dissolved total organic (TOC) and inorganic carbon (IC) (TOC-L CPN analyzer, Shimadzu, Japan), dissolved total nitrogen (Total N), ammonium (NH₄⁺) and nitrite + nitrate (NO₂⁻ + NO₃⁻) (AA500 AutoAnalyzer, SEAL Analytical, Germany). TOC and IC concentrations are expressed in mg L⁻¹ and Total N, NH₄⁺ and NO₂⁻ + NO₃⁻ in μg L⁻¹. Methane fluxes were measured on 12 and 20 September 2021 between 08:00 and 18:00. Fluxes were determined using the manual closed-chamber method (portable LI-7810 CH₄/CO₂/H₂O Trace Gas Analyzer, LI-COR Biosciences, United States) and a transparent polycarbonate chamber (height 38 cm, diameter 29 cm) equipped with a fan to ensure air circulation. The chamber was placed on the PVC collar (inner diameter 29.5 cm) to achieve an airtight seal and each measurement lasted 5 minutes. Methane flux rates are expressed as mg CH₄ m⁻² h⁻¹ (for more details, see (Järvi-Laturi *et al*. 2025)).

### Soil temperature

Soil temperature was measured in all study plots at 5 cm depth (T107 temperature probes, Campbell Scientific, United States). Temperature data were recorded at 10-minute intervals (CR1000X measurement and control datalogger, Campbell Scientific, United States). These data were used to calculate monthly mean soil temperatures for each plot from December 2020 to September 2021.

### Hydrological description and properties of the study site

Mineral-rich groundwater forms the main water source in the area, while surface runoffs also contribute, especially during spring snowmelt (Korhonen *et al*. 2025). Although the mild slope influences the hydrology and nutrient flow within the fen, the water table level in Puukkosuo has been found to be quite stable (Järvi-Laturi *et al*. 2025). Similarly, in the summer of 2021, water table dynamics were broadly similar between outside and inside the exclosure (**Supplementary Figure 3A**). More specifically, water table measurements from individual plots showed substantial spatial variability but strong overlap between the outside and inside areas (**Supplementary Figure 3B**). The level of the water table in relation to the ground level (moss surface) was measured weekly during 15.7.-27.9.2021. Across the late summer months, water levels in both areas remained close to the peat surface (generally within approximately ±5 cm), indicating a consistently saturated fen environment. However, topographic wetness index mapping and *in situ* soil moisture data indicated slightly higher water accumulation potential within the exclosure part of the fen, especially during July, August and early September (**Supplementary Figure 1, Supplementary Figure 4**), suggesting slightly wetter conditions in the inside area compared to outside area. Soil volumetric water content (VWC) in each plot was monitored at a depth of 5 cm (SM150T soil moisture sensors, Delta-T Devices, United Kingdom). All sensor data were recorded and stored (CR1000X measurement and control datalogger, Campbell Scientific, United States) at 10-minute intervals during 1 May to 30 September 2021. Thus, rather than representing a strong hydrological gradient, the two areas are best characterized as sharing a broadly similar saturated fen hydrological regime with small-scale spatial variability associated with microtopography and slope position. Such subtle hydrological variation may nevertheless contribute to differences in redox conditions, for example by promoting slightly more aerated microsites outside the exclosure area and somewhat more reduced microsites inside the exclosure (Blodau 2002, Estop-Aragonés, Knorr, and Blodau 2012).

Pore-water chemistry broadly supported this interpretation (**Supplementary Figure 5**). Variables reflecting the overall fen environment, such as pH, TOC and TC, showed similar seasonal trajectories between outside and inside the exclosure, consistent with a shared minerotrophic groundwater-influenced system. In contrast, some variables showed tendencies toward differences between the areas. Inorganic carbon (IC) tended to be higher inside the exclosure, which may reflect greater groundwater accumulation and associated bicarbonate inputs (Kemmers and Jansen 1988, Lamers *et al*. 2015). Nitrogen species also showed some divergence, including generally higher NO₃⁻+NO₂⁻ outside the enclosure and pronounced early-summer peaks in NH₄⁺, Total N and NO₃⁻+NO₂⁻ in the outside plots, with several months showing significant differences between the areas, although variability among the plots was high (**Supplementary Figure 5**). These peaks likely reflect the influence of spring snowmelt, which can mobilize nitrogen accumulated in peat during winter and generate short-lived pulses of dissolved nitrogen in pore water (Brooks, Williams, and Schmidt 1998, Rixen *et al*. 2022). The stronger expression of these peaks in the outside plots may result from more rapid flushing or mobilization of solutes following snowmelt, whereas the slightly wetter inside conditions may promote dilution, plant uptake, or more rapid microbial denitrification (Bremner and Shaw 1958, Luo, Tillman, and Ball 1999, Niboyet *et al*. 2025). Together, these patterns are consistent with subtle differences in nitrogen transformation processes along the slope, potentially associated with small differences in aeration and redox conditions. Seasonal snowmelt therefore likely plays an important role in establishing the hydrological and chemical conditions of the fen each year, with spring recharge and solute flushing followed by the development of spatial variation in pore-water chemistry during the snow free period. Consequently, although water table differences between the outside and inside areas were negligible, even minor variations in soil moisture can substantially influence oxygen availability and associated redox processes in peat soils.

### Peat soil sample collection and nucleid acid extraction

Peat samples for metagenomic (MG) and metatranscriptomic (MT) analyses were collected on 15.9.2021. Peat was sampled adjacent to the greenhouse gas (GHG) collar in each study plot (n = 36). A 4 × 4 cm section of peat was cut with a mineral wool knife. From the middle of each peat core, sample material was collected from approximately 15 cm depth from the top using forceps and transferred into plastic bags. To preserve nucleic acid integrity, the samples were immediately snap-frozen in liquid nitrogen using a dry-shipper in the field. All sampling tools were treated with 70% ethanol between each sample.

### Nucleic acid extraction and sequencing

Nucleic acid extraction was performed as described previously (Viitamäki *et al*. 2022). Briefly, 0.5 g of soil was processed using a hexadecyltrimethylammonium bromide (CTAB), phenol–chloroform and bead-beating protocol. DNA and RNA was purified with the AllPrep DNA/RNA Mini Kit (QIAGEN, Germany) and quantified using the Qubit dsDNA BR and RNA HS Assay Kit (ThermoFisher Scientific, United States). RNA integrity was assessed using the Agilent TapeStation 4150 system (Agilent Technologies, United States) with the High Sensitivity RNA ScreenTape assay. Illumina metagenome libraries were prepared with the Nextera XT DNA Library Preparation Kit (Illumina, United States). The NEBNext Ultra II Directional RNA Library Prep Kit for Illumina (New England Biolabs, United States) was used to prepare cDNA libraries. Concentrations were quantified using a Qubit fluorometer with the dsDNA BR/HS kit (Invitrogen, United States) and library quality was verified on a Fragment Analyzer (Advanced Analytical, United States). Sequencing was performed in paired-end mode (2 × 150 bp) on an Illumina NovaSeq 6000 (Illumina, United States) at the Institute of Biotechnology, University of Helsinki, Finland.

### Sequencing data processing and analysis

#### Quality control

The raw sequencing data was assessed with the high throughput quality control tools FastQC (Andrews 2010) version 0.11.9 and MultiQC (Ewels *et al*. 2016) version 1.19. CutAdapt (Martin 2011) version 4.6 (-m 50 and --nextseq-trim 20) was used to trim the adapter sequences and low quality reads. Paired-end reads were intentionally non-overlapping. During metagenomic library preparation with the Nextera XT DNA Library Preparation Kit, the DNA-to-transposome ratio was optimized to produce insert sizes larger than the combined read lengths, precluding the merging of R1 and R2 reads. Similarly the enzymatic shearing step in metatranscriptomic library preparation was targeted to produce approximately 300 bp fragments.

#### Taxonomy through Small-Subunit ribosomal RNA profiling

Microbial taxonomic composition in both MG and MT datasets was estimated using phyloFlash (Gruber-Vodicka Harald R., Seah Brandon K. B., and Pruesse Elmar 2020) v3.4.2 (-skip_spades -zip -log -poscov) and the preformatted SILVA 138.1 database (Pruesse *et al*. 2007, Chuvochina *et al*. 2026). To ensure appropriate taxonomic ranks for each nearest taxonomical unit (NTU) in the Phyloflash output, the SILVA 138.1 database taxonomy files were used to look up ranks for all hierarchy levels for each NTU. The PhyloFlash output files containing abundance data across all provided main taxonomic ranks (domain, kingdom, phylum, class, order, family, genus) were then retained for further analysis.

#### Ribosomal RNA removal from the metatranscriptomics data

To enable downstream analyses of microbial metabolic functional activity, ribosomal RNA sequences, which comprised the majority of reads in the total RNA, were removed from the MT data using SortMeRNA v4.3.6 (Kopylova, Noé, and Touzet 2012) (--fastx --paired_in --out2) with the sensitive reference database v4.3.

For additional details about the processing of sequencing data and downstream data analysis, see Supplementary Methods.

### Assembly, binning and processing of the metagenomic data

MG reads were co-assembled separately for samples from outside and inside the exclusion fence. For each co-assembly reads were pooled and assembled with MEGAHIT (Li D *et al*. 2015) v1.2.9 (--min-contig-len 1000 --k-min 57 --k-max 157 --k-step 10).

The assembled contigs were processed in anvi’o (Eren *et al*. 2021) v8-dev. Contigs were filtered to retain only contigs ≥ 2 kb, open reading frames were predicted with Prodigal (Hyatt *et al*. 2010) v2.6.3 and the single-copy core genes identified with anvi-run-hmms. KEGG KOs were annotated with anvi-run-kegg-kofams. To identify metabolic marker genes, contig gene sequences were exported and aligned with DIAMOND (Buchfink, Xie, and Huson 2015) v2.1.6 (--query-cover 80 --max-target-seqs 1 --outfmt 6 qseqid stitle pident length qstart qend sstart send evalue bitscore qcovhsp scovhsp slen) and the annotations were imported back into anvi’o.

MG and MT reads were mapped to contigs with Bowtie2 (Langmead and Salzberg 2012) v2.4.4 (--no-unal) and indexed with SAMtools (Li H *et al*. 2009) v1.18. Sample profiles were generated (anvi-profile) and merged (anvi-merge). Genome bins were generated with anvi-cluster-contigs (driver MetaBAT2 (Kang *et al*. 2019) v2.17) and manually refined with anvi-refine following minimum information about metagenome-assembled genome (MIMAG) (Bowers *et al*. 2017) standards (≥ 50 % completeness, < 10 % redundancy). Final taxonomy and phylogeny were assigned with GTDB-Tk (Chaumeil *et al*. 2020) v2.4.0 (database release 220). Bins from the two co-assemblies were dereplicated with anvi-dereplicate-genomes (FastANI (Jain *et al*. 2018) v1.34, similarity threshold 0.99). Relative MAG abundance was estimated with CoverM (Aroney *et al*. 2025) v0.7.0 (--methods relative_abundance --min-read-aligned-percent 80) using Minimap2 (Li H 2018) v2.28.

To obtain unbiased gene- and transcript-level abundance and expression estimates, contigs from both co-assemblies were also pooled into a single contig-based workflow and processed as above. All MG and MT reads were mapped once to this unified contig set to avoid double counting and minimize batch effects. Gene and transcript counts were generated with featureCounts (Liao, Smyth, and Shi 2014) (Subread v2.0.6). MAG genes were then linked to their corresponding contig-based workflow genes and counts for these genes were normalized as in the read-based analysis: reads per kilobase (RPK) adjusted for gene length and converted to transcripts or copies per million using total prokaryotic RPK per sample derived from the read-based KEGG alignments.

## Results

### The snow treatments alter soil temperatures during the winter season

During the time of the maximum snow depth in March 2021, the average snow depth did not differ significantly (Wilcoxon signed rank test significance value 0.58) between outside and inside the exclosure (**Supplementary Figure 2A**). As expected, a clear difference was observed due to the different snow treatments with snow removal plots having significantly less snow and snow addition plots having significantly more snow compared to the control ambient plots, indicating a successful snow manipulation treatment (**Supplementary Figure 2B**). Accordingly, the soil temperatures during the winter season were affected significantly by the snow treatments with temperatures being lower in the snow removal and higher in the snow addition plots from January to May, but not anymore during the summer and autumn period from June to September (**Supplementary Figure 2C**). Soil temperatures between plots outside and inside the exclosure did not differ significantly (Wilcoxon test p-values 0.21-0.78) during the measurement time from December 2020 to September 2021.

### Ordination shows clear separation of the soil microbiomes by the exclusion treatment

Ordination analyses revealed separation in the soil microbiome between plots outside and inside the exclosure (**Figure 2**). To comprehensively explore overall patterns between samples in the microbiome data, the Multiomics factor analysis (MOFA) (Argelaguet *et al*. 2018) framework was applied. From the MOFA ordination, which jointly integrated metagenomic (MG) and metatranscriptomic (MT) taxonomic as well as KEGG KO-level functional data, factors 3 and 6 were associated with the exclusion treatment (**Figure 2A**). Random forest classification of the MOFA factors further confirmed this pattern, with outside plot class status predicted with the highest accuracy (balanced accuracy = 0.75) (**Figure 2B**), compared to snow treatment or vegetation. When datasets were ordinated separately, both MG and MT taxonomic data (**Figure 2C**) and KO-level functional data (**Figure 2D)** showed moderate separation by the exclusion treatment.

**Figure 2.**
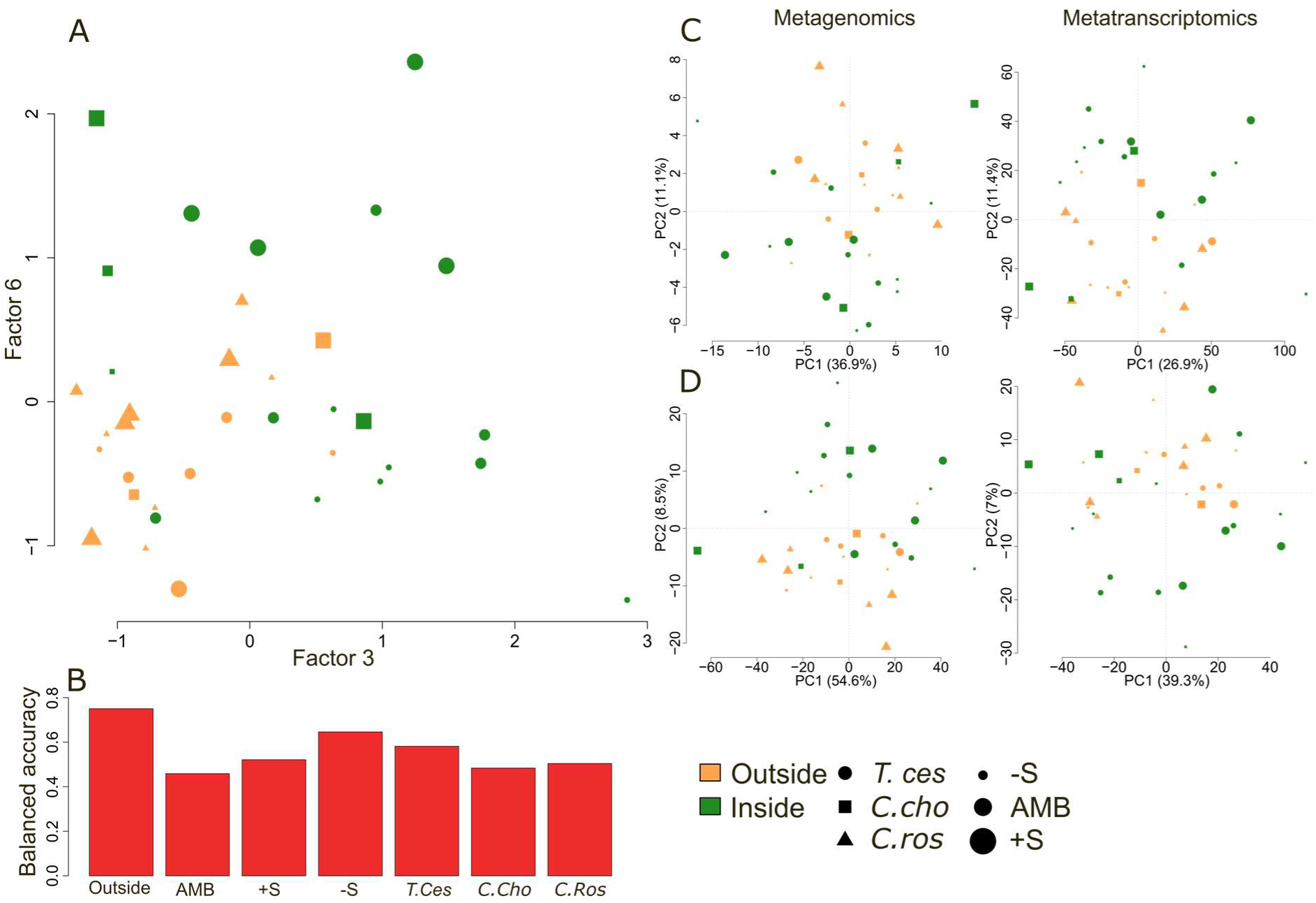
Ordination of the archaeal and bacterial soil microbiomes from the Puukkosuo fen. **A)** Multi-Omics Factor Analysis (MOFA) integrating metagenomic (MG) and metatranscriptomic (MT) taxonomic and Kyoto Encyclopedia of Genes and Genomes (KEGG) functional data, showing separation by the exclusion treatment. **B)** Balanced accuracies using leave-one-out cross-validation from random-forest models trained on MOFA factors to predict levels for the samples for the exclusion treatment, snow treatment and vegetation cluster. **C)** Principal component analysis (PCA) of MG and MT taxonomic profiles. **D)** PCA of MG and MT KEGG functional (KEGG Orthologous group (KO)-level) profiles. Abbreviations for the vegetation clusters, *T.ces* = *Trichophorum cespitosum*, *C.ros* = *Carex rostrata*, *C.cho* = *Carex chordorrhiza*. Abbreviations for the snow treatments, AMB = ambient snow cover (control), -S = snow removal, +S = snow addition. Taxonomic profiles were generated with phyloFlash and the SILVA 138.1 database, retaining only non-eukaryotic reads. Functional metabolic profiles were based on alignments to the KEGG prokaryote database summarized to KO – level, expressed as copies per million (as for transcripts-per-million) and log₂-transformed. All taxonomic data were CLR-transformed.

### Pseudomonadota dominate while the exclusion treatment alters order-level expression

The phyla *Pseudomonadota* and *Chloroflexota* dominated both MG and MT datasets (**Figure 3A**). On the other hand, several phyla showed variability between abundance (MG) and expression (MT): *Patescibacteria* were among the most abundant groups, but showed less expression whereas *Planctomycetota*, *Myxococcota*, and *Tectomicrobia* were less abundant but showed more expression. At the order level, the most abundant and expressed groups included *Burkholderiales*, *Rhizobiales*, *Anaerolineales*, *Pedosphaerales*, and *Geobacterales*. In contrast, *Methylococcales*, *Gemmatales*, *Polyangiales*, *Haliangiales*, and *Entotheonellales* were proportionally more expressed than abundant (**Figure 3B**).

**Figure 3.**
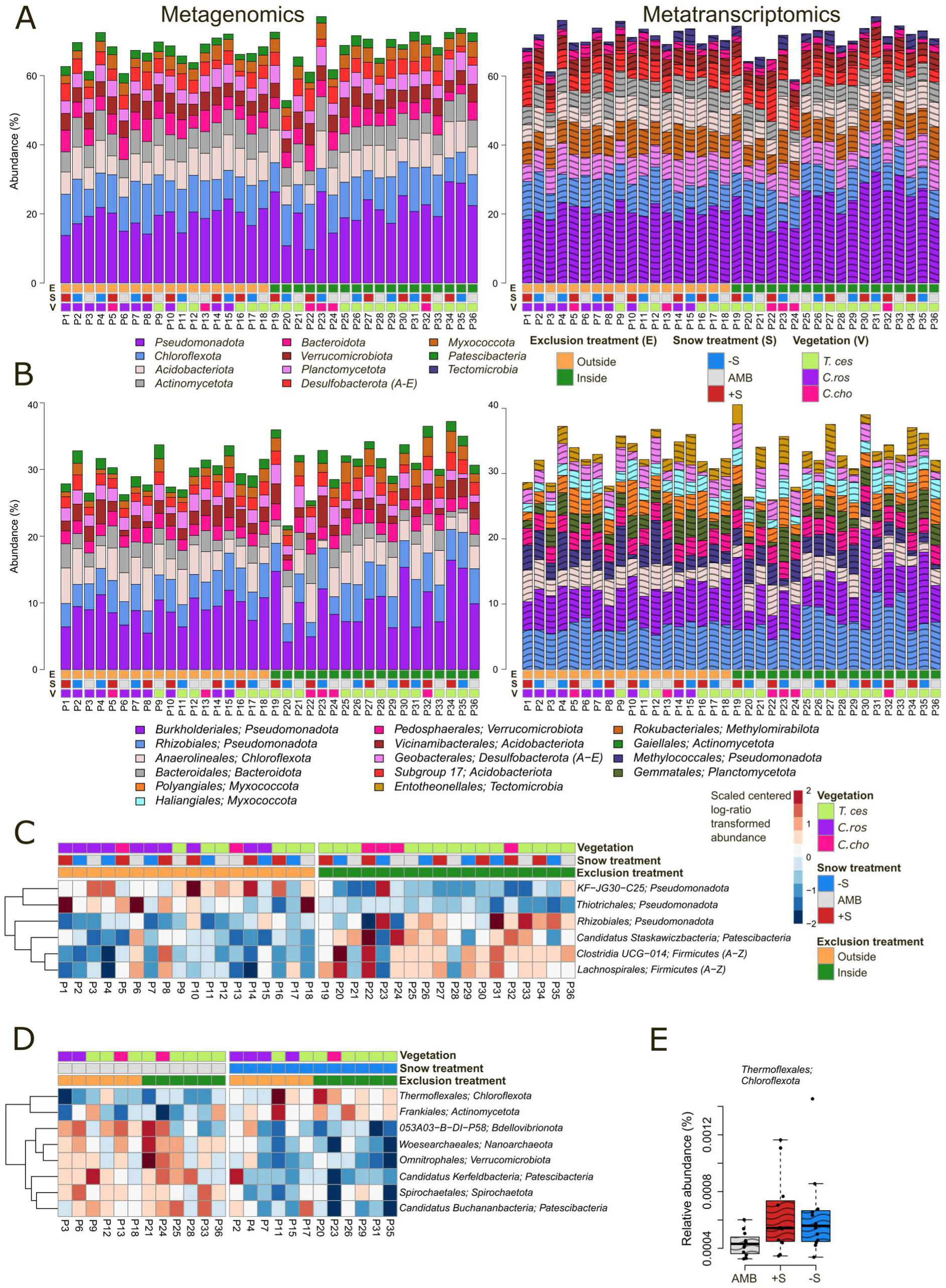
Taxonomic composition and differentially expressed archaeal and bacterial taxa. **A)** Relative abundance and expression of the ten most common phyla in metagenomic (MG) and metatranscriptomic (MT) datasets. **B)** Relative abundance and expression of the ten most common orders in MG and MT datasets. **C)** Relative expression (MT) of orders showing significantly differential transcript expression between plots outside and inside the exclusion fence. **D)** Relative expression (MT) of orders with differential expression between the snow removal and control plots. **E)** Orders with differential expression (MT) between the snow addition and control plots. Abbreviations for the vegetation clusters, *T.ces* = *Trichophorum cespitosum*, *C.ros* = *Carex rostrata*, *C.cho* = *Carex chordorrhiza*. Abbreviations for the snow treatments, AMB = ambient snow cover (control), -S = snow removal, +S = snow addition. Taxonomic profiles were generated with phyloFlash using the SILVA 138.1 Small Subunit (SSU) ribosomal RNA database and filtered to include only non-eukaryotic reads. Linear mixed-effects models (LMMs) (exclusion treatment and snow treatment as fixed effects, vegetation cluster as a random effect) were used to identify differentially expressed taxa (false discovery rate ≤ 0.1). For the LMMs the data were aggregated to order level and centered-log-ratio (CLR) transformed. The heatmaps (C–E) show scaled CLR-transformed expression (MT) clustered using hierarchical clustering and Euclidean distance.

Differential expression (DE) analysis revealed six orders with altered transcriptional expression associated with the exclusion treatment, one with snow addition, eight associated with snow removal and three with interactions between the exclusion and snow treatments (**Figures 3C–E, Supplementary Figure 6A, Supplementary Tables 1–4**). For example, within *Pseudomonadota*, *KF-JG30-C25* and *Thiotrichales* were more expressed while *Rhizobiales* and *Firmicutes* groups (*Clostridia UCG-014, Lachnospirales*) were less expressed outside the exclosure than inside. Snow removal plots showed reduced expression in orders of *Bdellovibrionota (053A03–B–DI–P58)*, *Verrucomicrobiota (Omnitrophales)*, *Nanoarchaeota (Woesearchaeales)*, *Patescibacteria (Candidatus Kerfeldbacteria, Candidatus Buchananbacteria)* and *Spirochaetota (Spirochaetales)* and increased expression of *Actinomycetota (Frankiales)* and *Chloroflexota (Thermoflexales)* in comparison to ambient snow (**Figure 3D**). Order *Thermoflexales* were more expressed under both snow removal and snow addition in comparison to ambient (**Figure 3D-E**). At the phylum level, *Iainarchaeota*, *Firmicutes* and the *SAR324* clades were less expressed outside than inside the exclosure, while *Actinomycetota* and *Aquificota* were more expressed under snow removal and *Abditibacteriota* were observed to have interactions between the exclusion and snow treatments (**Supplementary Figure 6B-D Supplementary Tables 5–7**).

In a previous laboratory study, the genus *Methanobrevibacter* was discovered to compose the majority (>90%) of the methanogenic archaeal community in reindeer droppings and the majority of archaeal communities in peat samples treated with reindeer droppings (Fritze *et al*. 2021). Consequently, we investigated whether this major group of reindeer rumen methanogenic archaea could be detected in this field study, potentially suggesting the presence and influence of reindeer in the study area. We explored the abundance and expression of *Methanobrevibacter* in our MG and MT data. Although *Methanobrevibacter* was not detected in MG data, low levels of transcripts were present in MT data (**Supplementary Figure 7**). Expression of *Methanobrevibacter* and the family *Methanobacteriaceae* were slightly higher in plots outside the exclosure than inside, although not statistically significant.

### Metabolic functionality links exclusion treatment to altered nitrogen cycling

**Figure 4.**
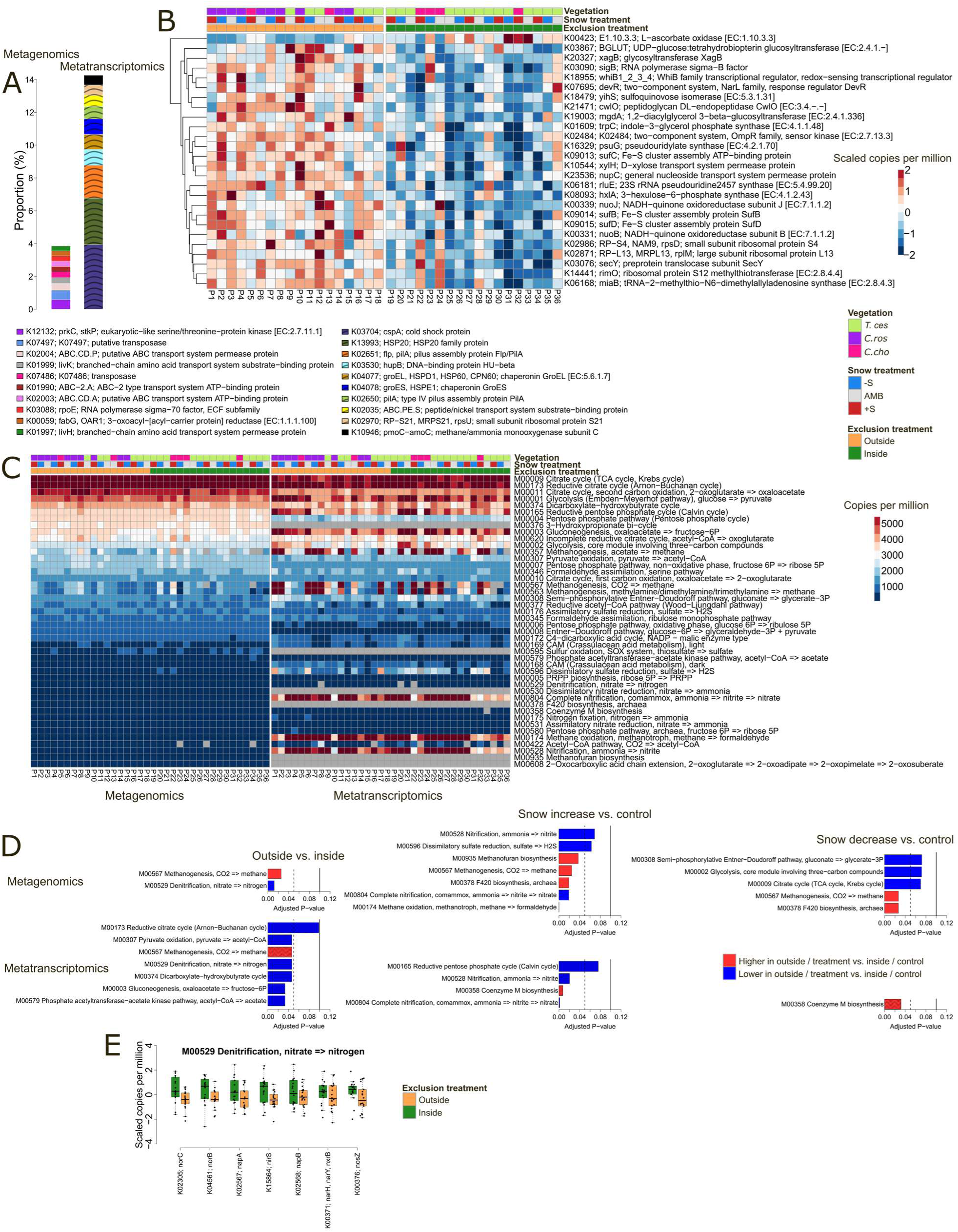
Metabolic functional potential and expression of the Puukkosuo archaeal and bacterial soil microbiome. **A)** The most abundant and expressed Kyoto Encyclopedia of Genes and Genomes (KEGG) orthologs (KOs) in metagenomic (MG) and metatranscriptomic (MT) datasets. **B)** KO groups differentially abundant (MG) between plots outside and inside the exclusion fence. **C)** Abundance (MG) and expression (MT) of the selected core KEGG metabolic modules detected in MG or MT data. Grey color indicates module not detected in the sample. **D)** Gene set enrichment analysis (GSEA) showing KEGG modules enriched (false discovery rate (FDR) ≤ 0.1) in MG and MT data for the exclusion treatment and the snow treatments. The dashed vertical line indicates FDR 0.05 and solid vertical line FDR 0.1. **E)** Scaled abundance of “leading-edge” KO groups driving GSEA enrichment of the denitrification module (M00529) in the exclusion treatment in the MG data. Abbreviations for the vegetation clusters, *T.ces* = *Trichophorum cespitosum*, *C.ros* = *Carex rostrata*, *C.cho* = *Carex chordorrhiza*. Abbreviations for the snow treatments, AMB = ambient snow cover (control), -S = snow removal, +S = snow addition. MG and MT reads were mapped to the KEGG prokaryote database and summarized to KO-level. KO counts were converted to copies-per-million (as for transcripts-per-million). Linear mixed-effects models (LMMs) (exclusion treatment and snow treatment as fixed effects, vegetation cluster as a random effect) were used to identify differentially abundant/expressed KOs (FDR ≤ 0.1). For the LMMs the KO data were log₂-transformed. KEGG modules were considered present when ≥75 % of their constituent KOs were detected. The heatmap in B show scaled log₂-transformed copies-per-million clustered using hierarchical clustering and Euclidean distance.

### Overall KO-level patterns

In MG, abundant KOs were linked to nutrient acquisition, lipid metabolism, stress response and genomic plasticity (**Figure 4A**). In MT, the most expressed KOs reflected stress adaptation, protein homeostasis, pilus-mediated interactions, information processing and nutrient uptake.

#### Differential KO abundance and expression

LMMs identified 26 KOs more abundant (FDR ≤ 0.1) in plots outside the exclosure than inside in MG, including gene clusters related to signal transduction, stress response (e.g. sigma factors), RNA/protein processing, nutrient acquisition, and redox-related energy and sulfur metabolism (**Figure 4B, Supplementary Table 8**). In MT, metH (methionine biosynthesis) was more highly expressed outside the exclosure than inside (**Supplementary Figure 8A, Supplementary Table 9**). Snow also influenced specific KOs: in MG, NAMPT (NAD⁺ cycling) was less abundant under snow removal while in MT, CIRBP (cold-shock RNA-binding protein) was more lowly expressed under snow removal (**Supplementary Figure 8B-C, Supplementary Tables 10-11**).

#### Core metabolic modules

Of the 52 targeted core metabolic KEGG modules (see Methods), 44 were present in MG and 39 in MT (**Figure 4C**). Highly represented modules in both MG and MT data were associated with the citric acid cycle, central carbon metabolism, energy metabolism, carbon cycling, and redox balancing. Interestingly, several methanogenesis, methane oxidation and nitrification related modules were markedly expressed in MT despite lower prevalence in MG.

#### Enrichment analysis

Gene set enrichment analysis (GSEA) (see Supplementary Methods) revealed significant enrichment (FDR ≤ 0.1) of nitrogen and methane cycling modules (**Figure 4D, Supplementary Tables 12–17**). Methanogenesis (CO₂ → CH₄) was enriched in both MG and MT data outside the exclusion fence compared to inside and in MG in both snow treatments compared to the ambient control. Denitrification (NO₃⁻ → N₂) was reduced in abundance (MG) and expression (MT) outside the exclusion fence compared to inside with several KO groups in the module consistently showing lower abundance in the outside plots compared to areas inside the exclosure (**Figure 4E**). Modules for complete nitrification (comammox) and methane oxidation were also less abundant and expressed under snow addition.

#### Urea associated KO groups

To investigate whether a signal associated with increased urea processing, suggesting increased urea contributions by the reindeer, could be detected outside the exclosure, we explored the abundance and expression of the key components of the urease enzyme in the MG and MT data. No differences between outside and inside the exclosure for the different urease enzyme subunits could be detected in neither abundance nor expression (**Supplementary Figure 9**).

### Metabolic marker genes

Several enriched KEGG modules were observed to be associated with nitrogen and methane cycling. To further expand our investigations to the N_2_O and CH_4_ associated metabolic pathways of the bacterial and archaeal soil microbiomes we utilized the Greening lab metabolic marker gene databases (Leung and Greening 2021) (see Supplementary Methods).

#### Nitrogen cycling

The marker gene analysis supported findings observed in the overall metabolic KEGG data (**Figure 5**). *norB* and *nosZ*, encoding enzymes for NO reduction and N₂O reduction, were significantly less expressed outside the exclosure than inside in MT data. *nirS* showed a similar but non-significant trend. In addition to the genes themselves, gene ratios *nrfA*-(*nirK*+*nirS*), assessing the potential of the microbial community for N retention (Saghaï *et al*. 2023) and (*nirS*+*nirK*)/*nosZ*, comparing the potential for N_2_O production to the potential for N_2_O reduction (Saarenheimo *et al*. 2015), were investigated. The *nrfA-(nirK+nirS)* was generally negative in both MG and MT (**Supplementary Table 18**). The *(nirS+nirK)/nosZ* ratio was consistently >1 (**Supplementary Table 18**). Moreover, significantly higher values for the (*nirS*+*nirK*)/*nosZ* ratio were observed for the plots outside the exclosure than inside in MT (**Figure 5A**).

**Figure 5.**
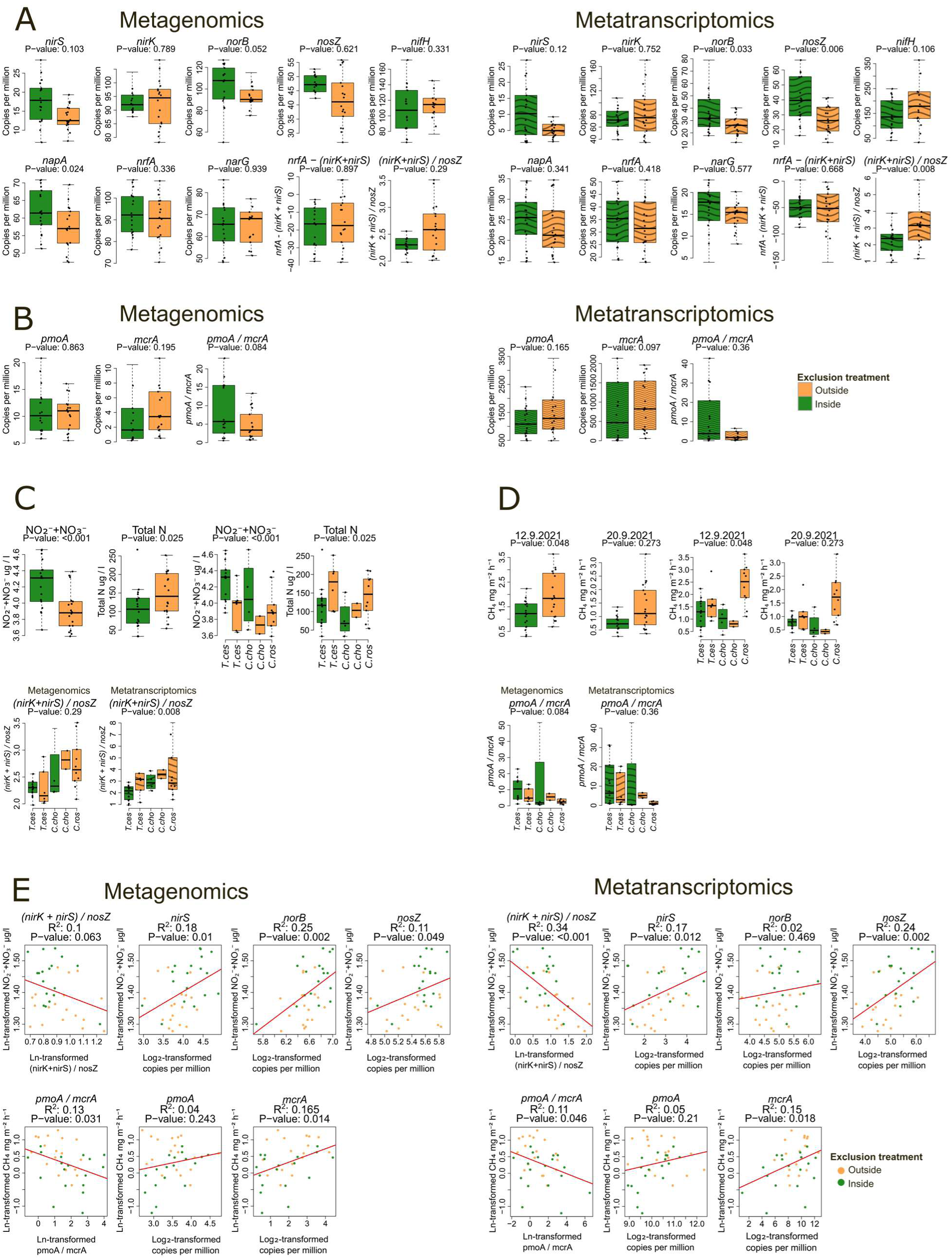
Soil archaeal and bacterial nitrogen- and methane-cycling marker genes, pore-water nitrogen and methane fluxes under the exclusion treatment in Puukkosuo. **A)** Relative abundance and expression of the key nitrogen-cycling marker genes and their functional ratios in metagenomic (MG) and metatranscriptomic (MT) data comparing plots outside and inside the exclusion fence. **B)** Relative abundance and expression of the methane-cycling marker genes and the methane oxidation/methanogenesis ratio in MG and MT data. **C)** September pore-water nitrate + nitrite (NO₃⁻ + NO₂⁻) and total nitrogen (Total N) by the exclusion treatment and vegetation cluster with the associated (*nirS + nirK)/nosZ* ratios in MG and MT data. **D)** Methane fluxes measured near the soil sampling dates by the exclusion treatment and vegetation cluster and the associated *pmoA/mcrA* ratios in MG and MT data. **E)** Linear regressions showing associations between pore-water NO₃⁻ + NO₂⁻ and the denitrification genes/ratios, methane fluxes and methane oxidation/methanogenesis genes/ratios. Abbreviations for the vegetation clusters, *T.ces* = *Trichophorum cespitosum*, *C.ros* = *Carex rostrata*, *C.cho* = *Carex chordorrhiza*. MG and MT reads were aligned to the eukaryote-filtered Greening Lab metabolic marker database and summarized to marker-gene level. Marker gene counts were converted to copies-per-million (as for transcripts-per-million). Linear mixed-effects models (LMMs) (exclusion treatment and snow treatment as fixed effects, vegetation cluster as a random effect) were used to determine significance of the MG abundance and MT expression association to the treatments. For the LMMs the marker gene data were log₂-transformed. Pore-water NO₃⁻ + NO₂⁻, Total N, methane fluxes and functional ratios were natural-log transformed for the regression analyses. Boxplots are plotted without potential outlier observations for better visualization.

#### Methane cycling

The methanogenesis marker gene *mcrA* was moderately more abundant and expressed outside the exclosure than inside (**Figure 5B**). The *pmoA*/*mcrA* ratio was generally > 1 (**Supplementary Table 18**). Interestingly, the ratios were lower, although not quite statistically significantly, in areas outside the exclosure compared to inside in MG and showed similar trends in MT (**Figure 5B**).

#### Snow effects and contig-based confirmation

In general, the snow treatments had little effect on the abundance and expression of the marker genes, except for lower *norB* expression in both snow addition and removal compared to the ambient control (**Supplementary Figure 10**). Analyses of the nitrogen and methane cycling marker genes and ratios in the assembled contigs yielded patterns consistent with the read-based analyses, though based on fewer reads (**Supplementary Figures 11–12**).

### Pore water chemistry and methane fluxes

In addition to the soil microbiome (MG and MT data) analyses, several pore water variables (**Supplementary Figure 5**, **Supplementary Figure 13**) were measured throughout the snow free period 2021 from May to September. To compare the relevant nitrogen and methane environmental variables with the soil microbiome data, we used September pore water data (**Supplementary Figure 13**), including nitrate+nitrite and total nitrogen (**Figure 5C**) as well as methane fluxes (**Figure 5D**), measured near (12.9.2021 and 20.9.2021) the soil microbiome data sampling date (15.9.2021). While the September concentrations of pore water nitrite+nitrate were significantly lower outside the exclusion than inside, the opposite was true for total nitrogen, with concentrations being significantly higher outside the exclosure (**Figure 5C).** However, it should be noted that during most of the snow-free period, nitrate+nitrite were higher outside the exclosure than inside (**Supplementary Figure 5**) with September values showing the concentrations converging towards more similar values. Methane fluxes were also significantly elevated in the outside plots compared to plots inside the exclusion (**Figure 5D**).

Pore water nitrate+nitrite and methane fluxes correlated with microbial functional traits **(Figure 5E**). Nitrate+nitrite levels were negatively correlated with the (*nirS*+*nirK*)/*nosZ* ratios and positively with several individual denitrification genes in both MG and MT data (**Figure 5E**). Methane fluxes were negatively associated with the *pmoA*/*mcrA* ratios and positively with *mcrA* abundance and expression (**Figure 5E, Supplementary Figure 14**).

### Taxa-function associations

As the ratios of especially nitrogen and also methane related functional genes were associated with exclusion treatment (**Figure 5**), we further explored how the abundance and expression of the order level taxa were correlated with these ratios (**Supplementary Figure 15**). Correlations linked methanogenic archaea (*Methanosarcinales*, *Methanomicrobiales*) with low *pmoA*/mcrA ratios, while several bacterial orders (e.g. *Gemmatales*, *Vicinamibacterales*) correlated with nitrogen gene ratios.

### Nitrogen-related processes align with the exclusion treatment, whereas methane patterns reflect vegetation differences

Because the exclusion treatment coincided with differences in vegetation composition, the ability of the linear mixed-effects models (LMMs) to fully disentangle exclusion effects from vegetation influences is limited. Although the LMMs indicated a small but significant effect of the exclusion treatment on methane fluxes (**Supplementary Figure 16),** the higher CH₄ emissions in the outside plots appear largely driven by *Carex rostrata* associated plant communities, which occur only outside the exclosure (**Figure 1C**, **Figure 5D**).

In contrast, nitrogen-related environmental variables and microbial functional indicators showed patterns more consistently associated with the exclusion treatment than with vegetation composition (**Figure 5C**). To further control for vegetation effects, we restricted additional analyses within the largest vegetation cluster present both outside and inside the exclosure, with *T. cespitosum* as the indicator species (**Supplementary Table 19**). Within this subset, the main observed trends observed in **Figure 5C** persisted: pore-water nitrate+nitrite concentrations (Wilcoxon test, p = 0.03), total nitrogen (p = 0.09), and the (*nirS*+*nirK*)/*nosZ* ratio in MT data (p = 0.11) differed, or showed near-significant differences, between the outside and inside sample plots.

### Metagenome assembled genomes

To complement the read-based analyses, MG reads were co-assembled separately for the plots outside and inside the exclosure. Contigs were processed with anvi’o (Eren *et al*. 2021) and binning with Metabat2 followed by manual refinement yielded 113 medium- and high-quality MAGs (average completion 70%, minimum 51%, average redundancy 4.6%, maximum 9.9%) (**Supplementary Tables 20-21**).

The taxonomic composition of the MAGs broadly matched the read-based results, with *Pseudomonadota*, *Chloroflexota*, *Actinomycetota* and *Desulfobacterota* as dominant phyla (**Figure 6A**). Additionally, MAGs included members of *Nitrospirota* and *Methylomirabilota*, phyla less frequent in the read-based community profiles. MAG abundances generally tended to be higher in the plots inside the exclusion, particularly for *Pseudomonadota*, *Methylomirabilota* and *Desulfobacterota* (**Figure 6B**). Many MAGs contained denitrification genes: *nirS* copies were restricted to *Pseudomonadota*, *nirK* was widespread across *Chloroflexota*, *Pseudomonadota*, *Actinomycetota* and *Nitrospirota*, *norB* occurred mainly in *Nitrospirota*, *Pseudomonadota*, *Acidobacteriota* and *Bacteroidata* while *nosZ* was less frequent, occurring mostly in *Acidobacteriota* and *Myxococcota*. For methane-related functions, only *pmoA* was detected (in two *Pseudomonadota* MAGs), whereas *mcrA* was absent.

**Figure 6.**
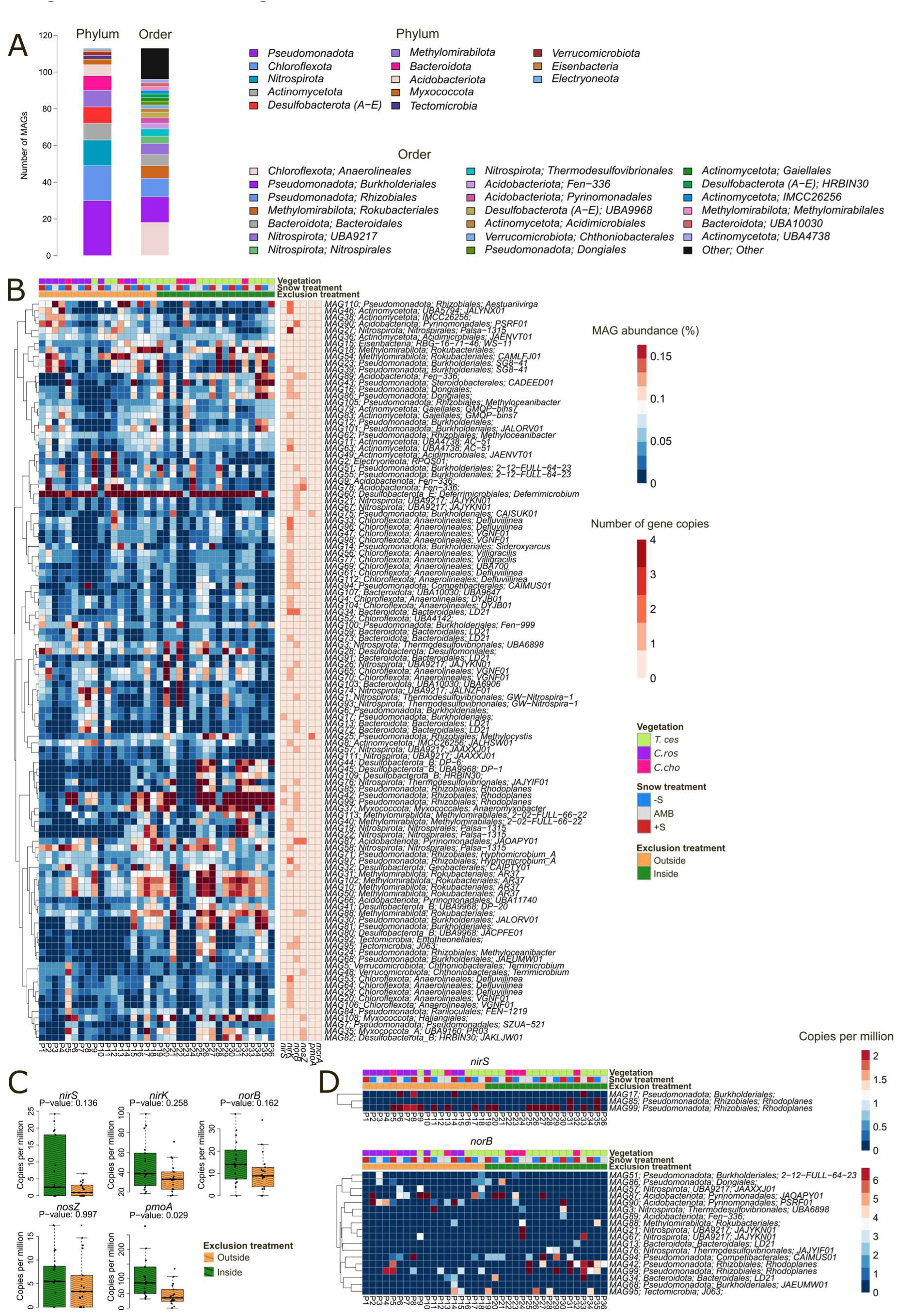
Metagenome-assembled genomes (MAGs) from the Puukkosuo soils. **A)** Taxonomic composition of the 113 medium- and high-quality MAGs, showing the numbers of phylum- and order-level annotations. **B)** Relative abundance of each MAG across the samples, with the accompanying counts of the key denitrification- and methane-cycling marker genes (right column). **C)** Expression of the key denitrification- and methane-related marker genes (summarized across all MAGs) in metatranscriptomic data comparing plots outside and inside the exclusion treatment. **D)** Examples of MAG-level expression patterns for two denitrification-related genes (*nirS* and *norB*). Abbreviations for the vegetation clusters, *T.ces* = *Trichophorum cespitosum*, *C.ros* = *Carex rostrata*, *C.cho* = *Carex chordorrhiza*. Abbreviations for the snow treatments, AMB = ambient snow cover (control), -S = snow removal, +S = snow addition. Metagenomic reads were co-assembled separately for the plots outside and inside the exclosure with MEGAHIT, processed in anvi’o and binned with MetaBAT2, followed by manual refinement to MIMAG standards (≥ 50 % completeness, < 10 % redundancy). MAGs were dereplicated using FastANI and taxonomy was assigned with GTDB-Tk. For panels B and D, MAGs are clustered by hierarchical clustering with correlation distance. MAG MT genes were aligned to the eukaryote-filtered Greening Lab metabolic marker database and summarized to marker-gene level. Marker gene counts were converted to copies-per-million (as for transcripts-per-million). Linear mixed-effects models (LMMs) (exclusion treatment and snow treatment as fixed effects, vegetation cluster as a random effect) were used to determine significance of the associations of the MT gene expression to treatments. For the LMMs the marker gene data were log₂-transformed.

Expression (MT) patterns of denitrification genes in the MAGs resembled those from the read-based analyses, though not statistically significant (**Figure 6C**). However, as the MAGs recruited only a subset of reads compared to the read-based approach, the expression patterns of the investigated MAG marker genes are less complete than those observed for the same genes in the read-based analysis. In particular, the methane-related genes were poorly represented, with only limited *pmoA* expression and no detectable *mcrA*. Furthermore, most observed transcriptional patterns were driven by only a few or some MAGs (**Figure 6D, Supplementary Figure 17**). Consequently, as the expression of *nirS*, *nirK* and *nosZ* was zero in many samples even over all MAGs, ratios such as (*nirS*+*nirK*)/*nosZ* could not be reliably calculated from the MAG data. Together, these genome-resolved data support the read-based findings but highlight that the latter provide a more comprehensive view of functional gene prevalence and expression across the peatland microbiome.

## Discussion

This study examined how reindeer exclusion, altered snow depth, and small-scale hydrological variation jointly shape the soil microbiome in a northern peatland using parallel metagenomic (MG) and metatranscriptomic (MT) approaches. While MG delivers information on the microbial community and its metabolic potential as a whole, it does not discriminate between the active and non-active, living and dead microbes (Ojala *et al*. 2023). MT on the other hand complements the MG measurements and offers a view into the living and active members and metabolic functions of the microbial community.

Three hypotheses were tested: whether (1) reindeer exclusion and (2) altered snow depth affect microbial community structure and metabolic functionality, particularly pathways involved in N₂O and CH₄ cycling, and whether (3) variation in soil moisture and associated redox conditions influences these microbial responses. Although clear differences in microbial community composition and greenhouse gas–related processes were observed in the exclusion treatment, these patterns were not directly attributable to reindeer presence. Instead, they are best explained by underlying hydrological variation and, for methane dynamics, differences in plant community composition, supporting hypothesis (3). Snow manipulation had more limited effects, primarily influencing the expression of specific taxa and the module-level microbial responses for nitrogen and methane cycling, providing partial support for hypothesis (2). In the following chapters of discussion, we address the findings of this study in more detail.

### Hydrology, rather than reindeer presence, structures microbial responses in the exclusion treatment

Hydrology is a primary determinant of peatland structure and function, regulating both geochemical conditions and microbial processes (Rydin and Jeglum 2013, Nunes *et al*. 2015). In peat soils, both microbial community composition and metabolic activity are associated with hydrological and geochemical gradients. Differences in water flow and nutrient availability, particularly within and across minerotrophic fens, strongly influence these microbial communities (Lin *et al*. 2012, Andersen, Chapman, and Artz 2013, Urbanová and Bárta 2014, Seward *et al*. 2020).

Although the exclusion treatment appeared as the main structuring factor across both taxonomic and functional profiles of the soil microbial community, this pattern plausibly reflects underlying subtle hydrological differences between the outside and inside areas rather than a direct effect of reindeer presence/grazing. The experimental layout for the exclusion treatment follows a natural, mild hydrological gradient in the area. Accordingly, plots inside the exclosure showed a higher water accumulation potential and indications of stronger groundwater influence, including elevated inorganic carbon concentrations consistent with bicarbonate inputs (Kemmers and Jansen 1988, Lamers *et al*. 2015). In contrast, plots outside the exclosure exhibited higher concentrations of NO₃⁻+NO₂⁻ and pronounced early-season peaks in NH₄⁺ and total nitrogen, suggesting differences in nutrient availability linked to water flow dynamics and associated microbial responses.

Reindeer presence has previously been connected to nitrogen cycling by increasing gross or net nitrogen mineralization in northern Fennoscandia and Finnish Lapland (Stark *et al*. 2000, Stark, Strömmer, and Tuomi 2002, Johan Olofsson 2009). Such effects are suggested to be primarily regulated by reindeer-mediated fertilization (e.g. urine / feces) with stimulation to primary production and microbial respiration in nutrient-rich systems (Stark *et al*. 2000, Stark, Strömmer, and Tuomi 2002, Johan Olofsson 2009). However, multiple lines of evidence indicate that reindeer-derived inputs played a minor role in structuring the observed microbial patterns in this study. Despite the occasional observed presence of reindeer in the area, no differences were detected in the abundance or expression of urease genes, which would be expected if substantial urea inputs from urine were influencing nitrogen cycling. Likewise, only low levels of *Methanobrevibacter* transcripts, the main methanogenic archaea in reindeer droppings (Fritze *et al*. 2021), were observed, with no significant treatment-related differences. The minor influence of reindeer to the Puukkosuo soil microbiome is further supported by the preliminary findings from a yet unpublished data (Maria Väisänen et al.) on long-term (2018-2024) vegetation responses due to the exclusion treatment in Puukkosuo. There, only limited support was found for reindeer grazing effects on vegetation, suggesting that other environmental factors (e.g. hydrological differences) could be the primary drivers for the observed effects. Together, these observations suggest that hydrological variation and associated differences in redox conditions and nutrient availability, rather than reindeer presence, is the primary driver of the microbial differences associated with the exclusion treatment. However, reindeer presence may still potentially fine-tune these responses through for example soil disturbance (e.g., trampling) or nutrient hotspots (urine, feces), but the effects are likely small and localized. Future work that combines high-resolution hydrological measurements with data on reindeer movement and density, would help to disentangle these now intertwined effects of hydrology and reindeer.

### Nitrogen cycling along the exclusion treatment is consistent with hydrologically mediated differences in redox conditions

Patterns in nitrogen cycling were consistent with hydrologically mediated differences in redox conditions. Although redox conditions were not directly measured, the combination of soil moisture patterns, pore-water chemistry, and functional gene expression suggests variation in oxygen availability between the plots. The overall negative *nrfA*-(*nirK*+*nirS*) ratios indicate a general dominance of denitrification over dissimilatory nitrate reduction to ammonium, while overall (*nirS*+*nirK*)/*nosZ* ratios greater than one suggests a system with stronger potential for N₂O production relative to reduction.

Despite this, expression patterns of denitrification genes indicate a shift toward more complete denitrification inside the exclosure as compared to outside. More specifically, significantly higher expression of *norB* and *nosZ*, combined with lower (*nirS*+*nirK*)/*nosZ* ratios, suggests a greater potential for complete denitrification to N_2_ instead of N₂O under the slightly wetter and resulting more reduced conditions inside the exclosure. These findings align with previous with previous work from fens, where increased soil moisture has been connected to promoting complete microbial denitrification to N_2_ (Rückauf *et al*. 2004, Lohila *et al*. 2010). Additionally, while (Palmer and Horn 2015) observed stimulated N_2_O production briefly due to nitrate-addition to the soil in Puukkosuo, the authors concluded that the absence of *in situ* N_2_O emissions from Puukkosuo is likely caused by the occurrence of complete denitrification with N_2_ as the end-product, a pattern observed also in previous studies from pristine fens at ambient nitrate levels (Lohila *et al*. 2010, Roobroeck *et al*. 2010).

Importantly, these nitrogen-related patterns were consistent across vegetation clusters. Linear mixed-effects models accounting for vegetation as a random factor, as well as analyses restricted to the largest vegetation cluster, confirmed that the observed differences were not driven by plant community composition. This suggests that nitrogen cycling in this system is primarily controlled by environmental gradients associated with hydrology, rather than vegetation or reindeer presence.

Genome-resolved analyses complemented these findings by linking denitrification potential to diverse microbial taxa. Expression patterns were broadly similar to the read-based findings but were weaker and less statistically robust because MAGs recruited only a subset of all the reads. This highlights the value of genome-resolved data for taxonomic context, while confirming that read-based approaches can provide a more comprehensive picture of functional capacity and expression in complex peat microbiomes.

### Methane dynamics are primarily linked to vegetation composition

In contrast to nitrogen cycling, methane dynamics appeared more strongly associated with vegetation than with hydrological differences or the exclusion treatment. Although methane oxidation potential exceeded methanogenesis overall (*pmoA*/*mcrA* > 1), both marker gene data and flux measurements indicated relatively increased methanogenesis in plots outside the exclosure compared to inside. Plots outside the exclosure showed trends for both higher *mcrA* abundance and expression, lower *pmoA*/*mcrA* ratios and higher CH₄ fluxes. Closer inspection revealed that these patterns were linked to the presence of *Carex rostrata*, which occurred exclusively outside the exclosure. Sedges, such as *C. rostrata*, have previously been associated with CH_4_ emissions in peatlands and northern fens (Noyce *et al*. 2014, Ge *et al*. 2023) and also very recently in the same study sites in Oulanka during the subsequent year to our sampling (Järvi-Laturi *et al*. 2025), further supporting vegetation as the likely primary driver of the observed methane-related responses.

Consistent with this interpretation, dominant methanogenic archaea, *Methanosarcinales* and *Methanomicrobiales*, were both abundant and transcriptionally expressed and displayed strong negative correlations to the *pmoA*/*mcrA* gene ratios but did not differ significantly between treatments when controlling for vegetation. Both archaea orders are commonly abundant in peatlands, especially in minerotrophic fens (Browne *et al*. 2017, Finn *et al*. 2020, Chen *et al*. 2023). *Methanomicrobiales* are exclusively hydrogenotrophic, they reduce CO₂ using H₂ to produce CH₄ (Browne *et al*. 2017). Members of *Methanosarcinales* on the other hand are metabolically divergent and possess the ability for methanogenesis via acetoclastic, hydrogenotrophic, and methylotrophic methanogenic pathways (Liu and Whitman 2008, Vanwonterghem *et al*. 2016).

While reindeer droppings have been connected to increased methane production potential in laboratory studies of peat soils (Laiho, Penttilä, and Fritze 2017, Fritze *et al*. 2021), this connection was not detected in a subsequent field study (Laiho *et al*. 2024). Rather, increased methane fluxes were concluded to be connected to vegetation, in particular increased sedge leaf area (Laiho *et al*. 2024). Taken together, the findings from the current study align well with previous work showing that vegetation, rather than reindeer presence, is the primary driver of methane emissions in similar systems. While hydrology can influence methanogenesis through its effects on redox conditions and substrate supply, vegetation appears to exert a stronger control on methane dynamics in this fen.

### Snow manipulation has modest effects on microbial responses

Snow treatments, especially snow removal, influenced the microbial communities, although these effects were modest and primarily reflected in differential expression of specific taxa. Snow cover acts as a thermal insulator, maintaining relatively stable winter soil temperatures (Ruan and Robertson 2017), and its removal likely exposes the microbial communities to greater environmental variability.

Reduced expression under snow removal was observed for several taxa with limited metabolic autonomy or strong dependence on microbial interactions, including members of *Bdellovibrionota*, *Omnitrophales*, *Woesearchaeales*, and candidate phyla *Candidatus Kerfeldbacteria*, *Candidatus Buchananbacteria*. These groups encompass bacterial predators (Kadouri and O’Toole 2005), symbionts (St. John and Reysenbach 2019, Huang *et al*. 2021, Srinivas *et al*. 2024) and ultra-small (Srinivas *et al*. 2024) microorganisms, and their reduced expression may indicate a weakening of tightly coupled microbial interaction networks under more variable conditions in snow removal. In contrast, the increased expression of *Frankiales* and *Thermoflexales*, a member of the metabolically versatile phylum *Chloroflexota* (Freches and Fradinho 2024), suggests a shift toward more independent heterotrophic strategies, potentially reducing reliance on tightly coupled microbial interactions (Barka Essaid Ait *et al*. 2015, Freches and Fradinho 2024, Dobrzyński *et al*. 2026).

Functional enrichment analyses indicated altered representation of methane oxidation, methanogenesis and nitrification modules under snow addition and methanogenesis under snow removal. Despite these overall functional patterns, only a few individual KOs or marker genes related to CH₄ or N₂O cycling (with the exception of *norB*) were significantly affected, suggesting a limited impact for the snow treatments on microbial functionalities of these processes. The results are consistent with previous studies showing variable microbial responses to snow manipulation, ranging from strong effects in long-term experiments to minimal changes where seasonal variation dominate (Ricketts *et al*. 2016, Männistö *et al*. 2018). While the snow treatments had the expected effect on both snow depth and soil temperatures during winter and spring (both lower in snow removal and higher in snow addition compared to ambient), this difference in soil temperatures had leveled out by June with no significant differences between the plots in the different snow treatments. Consequently, stronger effects in the soil microbiomes might emerge in early summer closer to snowmelt, when the soil temperature and moisture changes are more pronounced.

### Microbial community composition reflects typical peatland assemblages

Across both MG and MT datasets, the microbial community was dominated by *Pseudomonadota* and *Chloroflexota*, with substantial contributions from *Acidobacteriota* and *Actinomycetota*. These groups are commonly observed in peatland ecosystems and encompass a wide range of metabolic functions, including carbon degradation, methane oxidation, and nitrogen transformations (Zhou *et al*. 2017, Kujala *et al*. 2018, Kolton Max *et al*. 2022, Rakitin *et al*. 2022, Bovio-Winkler *et al*. 2023, Freches and Fradinho 2024). In a previous study from Puukkosuo utilizing 16S rRNA gene amplicon sequencing (Kujala, Schmidt, and Horn 2024), *Proteobacteria* (current *Pseudomonadota*), *Actinobacteriota* (current *Actinomycetota*), *Acidobacteria* (current *Acidobacteriota*) and *Firmicutes* were among the most abundant groups, consistent with the results in this study. Similarly, genome-resolved analyses showed similar taxonomical patterns to the read-based analysis, supporting the consistency of these results with previous studies of fen microbiomes.

### Synthesis and implications for greenhouse gas cycling

This study aimed to test three hypotheses concerning the effects of reindeer exclusion, snow depth, and subtle hydrologic variation on peatland microbial communities, particularly microbial pathways involved in N₂O and CH₄ cycling. While the exclusion treatment was associated with clear differences in microbial community composition and processes related to N₂O and CH₄, these differences are best explained by underlying hydrological and environmental variation rather than direct effects of reindeer presence. Thus, hypothesis (1) was not supported.

In contrast, support was found for hypothesis (3): variation in soil moisture and associated redox conditions influenced nitrogen cycling, particularly by promoting more complete denitrification under slightly wetter conditions. Methane dynamics, however, were primarily linked to vegetation composition, more specifically to the presence of C. rostrata, highlighting the importance of plant–microbe interactions in regulating CH₄ fluxes.

Snow manipulation provided partial support for hypothesis (2), with especially snow removal resulting in detectable effects on specific taxa and both snow treatments altering microbial modules participating in N₂O and CH₄ cycling, although not mainly on individual marker gene level. These findings suggest that while winter conditions can influence microbial communities, their impact may be transient or secondary to other environmental drivers in this system.

Taken together, our results suggest hydrological variation as the major driver of the microbial processes relevant to greenhouse gas dynamics in northern peatlands. By modulating redox conditions and nutrient availability, water flow dynamics can regulate denitrification pathways, while vegetation may exert stronger control over methane dynamics. While no strong support was detected for reindeer effects on the peat microbiome, reindeer may still locally fine-tune these processes through trampling and nutrient inputs but the effects are likely small and secondary to other environmental factors. These findings emphasize that hydrology, vegetation, herbivory and microbial processes are tightly coupled in peatlands, particularly in nutrient-rich fens. Future studies combining factorial vegetation and grazer manipulations with specific consideration for the hydrological environment, will be necessary to clarify these interactions.

### Conclusions

By integrating metagenomic and metatranscriptomic data with pore-water chemistry and methane flux measurements, this study provides a multi-layered view of the environmental controls shaping peatland microbiomes. Hydrology emerges as the primary driver structuring both microbial composition and function, primarily through its effects on redox conditions and nutrient availability. Vegetation plays a key role in regulating methane dynamics, while potential reindeer effects are likely only minor and localized. Understanding these interacting controls is essential for predicting carbon and nitrogen cycling feedbacks in northern ecosystems under climate and land-use change.

## Acknowledgements

The authors want to acknowledge and thank Eero Koskinen for conducting the methane flux measurements and methane flux calculations.

The authors want to acknowledge and thank Petra Korhonen for the SAGA wetness information and image for Supplementary Figure 1.

The authors want to acknowledge and thank Anne Tyvijärvi for the reindeer visualization used in the graphical abstract.

The authors would like to acknowledge CSC - IT Centre for Science, Finland for making this study possible through supplying the needed computational infrastructure and resources. The authors used OpenAI’s ChatGPT (GPT-5, accessed September 2025 – May 2026) to assist with language editing and improving readability. The text was originally written by the authors, who reviewed and approved the final manuscript in its entirety.

## Author contributions

T.V. methodology, software, formal analysis, investigation, writing - original draft, writing - review & editing, visualization, data curation. H.F. conceptualization, investigation, writing - review & editing, supervision, project administration, funding acquisition. J-M.P. investigation, writing - review & editing. K.P. conceptualization, investigation, writing - review & editing. E.J-L. investigation, formal analysis, writing - review & editing. T.R.C. conceptualization, supervision, writing - review & editing. M.V. conceptualization, investigation, resources, funding acquisition, writing - review & editing. J.L. investigation, writing - review & editing. R.P. conceptualization, writing - review & editing, resources, funding acquisition. J.H. conceptualization, methodology, investigation, resources, writing - review & editing, supervision, project administration, funding acquisition.

## Data availability

The raw sequencing data for this study is available at European Nucleotide Archive (ENA) with the project accession number PRJEB104445.

The used code for analysis in this study is available at https://github.com/ArcticMicrobialEcology/puukkosuo-soil-microbes2021

