## Supplementary Methods for "Soil microbiome structure and function reflect environmental variation rather than reindeer presence in a northern peatland"

<u>Processing sequencing data</u>

*Overall metabolic functional potential and activity*

To assess the overall metabolic functional potential of the metagenomes (MG) and the realized metabolic functional activity of the metatranscriptomes (MT), trimmed sequencing reads from each sample were aligned to the Kyoto Encyclopedia of Genes and Genomes (KEGG) prokaryote database (release 86) (Kanehisa and Goto 2000) using DIAMOND blastx v2.1.6 (Buchfink, Xie, and Huson 2015) (--query-cover 80 --max-target-seqs 1 --outfmt 6 qseqid stitle pident length qstart qend sstart send evalue bitscore qcovhsp scovhsp slen). KEGG functional orthologous group (KO) annotations for the KEGG genes in the used KEGG prokaryote database were downloaded using the *KEGGREST* R-package (Tenenbaum and Maintainer 2024) version 1.4.2. Paired-end reads (R1 and R2) were aligned independently against the KEGG database.

To leverage information from both mates while avoiding ambiguous assignments, a hierarchical filtering procedure was applied to combine the R1 and R2 alignments. For read pairs where only one mate (R1 or R2) produced a KEGG gene hit, the corresponding alignment was retained.

For read pairs where both mates produced KEGG gene alignments, the following rules were applied:

1. If both mates aligned to the same KEGG gene, the R1 alignment was retained.
2. If mates aligned to different KEGG genes and the absolute bitscore difference was ≥15, the alignment with the higher bitscore was retained.
3. For remaining cases without a clear bitscore difference, if only one of the alignments had an associated KEGG Ortholog (KO) annotation, that alignment was retained.
4. If both alignments had KO annotations, only shared KO annotations between R1 and R2 were retained. If no overlap in KO annotations was observed, the read pair was discarded as ambiguous.
5. For cases where neither alignment had a KO annotation, similarity between KEGG gene descriptions was evaluated using the stringsim function (stringdist R package v0.9.12, optimal string alignment / restricted Damerau–Levenshtein distance). Read pairs with description similarity < 0.7 were discarded as ambiguous.

This procedure leverages paired-end information while minimizing ambiguous functional assignments. For downstream analyses, only KO-annotated KEGG gene hits were used to construct KO-level abundance/expression matrices. However, all retained KEGG gene hits (including those without KO annotations) were used to estimate total prokaryotic reads per kilobase (RPK) per sample for TPM adjustment.

A schematic overview and representative assignment scenarios for KO annotated KEGG genes are shown in **Supplementary Figure 18**.

*Metabolic marker genes*

To further examine microbial metabolic functional potential and activity in the MG and MT, we used the Compiled Greening Lab metabolic marker gene database (Leung and Greening 2021). As the focus of this study was on the non-eukaryotic microbiome, the *taxonomizr* R package v0.10.6 (Sherrill-Mix 2023) was used to download the annotated National Center for Biotechnology Information (NCBI) protein database and remove all annotated eukaryotic proteins from the marker gene database. The NCBI taxonomy database was further supplemented with manually identified eukaryotic proteins (**Supplementary Table 22**).

Trimmed sequencing reads were aligned to the filtered marker gene database using DIAMOND blastx (--query-cover 80 --max-target-seqs 1 --outfmt 6 qseqid stitle pident length qstart qend sstart send evalue bitscore qcovhsp scovhsp slen). Paired-end reads (R1 and R2) were aligned independently against the marker gene database. The alignment results were further filtered with minimum percentage identity thresholds set at 80% (*psaA*), 75% (*hbsT*), 70% (*psbA, isoA, atpA, ygfK, aro*), 60% (*coxL, mmoA, amoA, nxrA, rbcL, nuoF, fefe* hydrogenases, *nife* Group 4 hydrogenases), or 50% (all other genes), following (Lappan *et al.* 2023). To leverage information from both mates, while controlling for ambiguous assignments, a similar filtering procedure to the KEGG gene alignments was applied. In short:

1. For read pairs where only one mate (R1 or R2) produced a marker gene hit, the corresponding alignment was retained.
2. If both mates aligned to the same marker gene, the R1 alignment was retained.
3. If mates aligned to different marker genes and the absolute bitscore difference was ≥15, the alignment with the higher bitscore was retained.
4. For remaining cases without a clear bitscore difference and the mates aligned to different marker genes, the read pair was discarded as ambiguous.

The number of total reads and the proportion of reads used for various data types (KEGG, metabolic marker gene data, taxonomic data) is given in **Supplementary Table 23**.

<u>Downstream data analysis</u>

*Processing of microbial taxonomic community data*

Nearest taxonomic unit (NTU) count data with ensured appropriate taxonomic rank annotations obtained from phyloFlash were compiled into a matrix and all eukaryotic counts were aggregated and removed from further analysis. For ordination analyses (Multiomics Factor Analysis, Redundancy Analysis), NTU data were handled within the *phyloseq* R package v1.46.0 (McMurdie and Holmes 2013). Unaggregated NTU data were transformed to compositional form using the transform function in the *microbiome* R package v1.24.0 (Lahti and Shetty 2017). To remove spurious taxa, only NTUs with a relative abundance/expression equivalent to at least five counts in the sample with the smallest total NTU count (smallest rRNA library size) in at least six samples (equal to the number of samples in the smallest vegetation cluster group, C. cho) were retained. The filtered NTU data were then centered log-ratio (CLR) transformed using the transform function in the *microbiome* package, a common approach for microbiome abundance data (Gloor *et al.* 2017).

For differential abundance (DA) (MG) and differential expression (DE) (MT) analyses, unfiltered NTU data were summarized to order and phylum levels. Rare taxa were aggregated using the aggregate_rare function in the *microbiome* package, applying the same detection threshold (relative abundance/expression equivalent to five counts in the smallest library) and prevalence threshold (≥ six samples) as in the ordination filtering. Both order- and phylum-level datasets were then CLR-transformed prior to analysis. All steps were applied identically to both MG and MT NTU datasets.

For the visualization of the taxonomy data in the main **Figure 3**, the phyloFlash data utilizing the SILVA 138.1 database phylum annotations were manually mapped to GTDB phylum level annotations.

*Processing of microbial functional metabolic data*

Sample-wise unique alignment hit counts (for both KEGG and metabolic marker gene datasets) were first length-normalized to account for variations in gene size and expressed as reads per kilobase (RPK). RPK values were then summarized to KEGG ortholog (KO) level for KEGG data and to marker gene level for the marker gene dataset and compiled into matrices. These RPK values were further transformed to transcripts per million (TPM), a standard approach in sequencing data analysis (Conesa *et al.* 2016) that has been also applied to both MG and MT datasets (Hatch *et al.* 2019, Krinos *et al.* 2024, Villette *et al.* 2025). TPM accounts for differences in both gene length and total sequencing depth, allowing more accurate within- and between-sample comparisons of abundance and expression. In this study, TPM-adjusted values for both MG and MT datasets are jointly referred to as “copies per million.” The total prokaryotic RPK per sample (sum of RPK for all genes in a sample) used in TPM calculation was derived from KEGG alignments by summing the RPK for all the unique KEGG prokaryotic genes detected in that sample. For KEGG genes mapping to multiple KO groups, the RPK value was assigned to each KO group and the sample-wise RPK sum was adjusted by including the duplicated counts to maintain comparability across samples.

Functional metabolic copies per million data were filtered using the same criteria as for the taxonomic datasets: only KO groups or marker genes with a relative abundance/expression equivalent to ≥ five counts in the sample with the smallest library size (based on total KEGG gene alignment hits) in at least six samples were retained. This approach follows common sequencing data filtering practices, including those described in the *edgeR* R package manual (Robinson, McCarthy, and Smyth 2010), to reduce noise from low-abundance/expression features. Ratios of marker genes were calculated on filtered TPM-adjusted marker gene data. Prior to downstream analyses, filtered data were $\log_2$-transformed with an offset of 1. This transformation is commonly used for omics data (Huber *et al.* 2002, Karpievitch, Dabney, and Smith 2012, Xia 2023) to improve normality, stabilize variance and reduce the dependence of variance on feature abundance (i.e., mitigating the typically greater variance observed for highly expressed features compared to lowly expressed ones in RNA-seq data).

In summary, all taxonomic data were CLR-transformed and all functional data (KO groups, marker genes) were TPM-normalized (referred to as “copies per million”) and $\log_2$-transformed before downstream analyses.

*Ordination of the microbial taxonomic and metabolic functional data*

Multiomics Factor Analysis (MOFA) (Argelaguet *et al.* 2018), implemented in the *MOFA2* R package v1.14.0, was used to jointly ordinate MG and MT microbial taxonomic and metabolic functional data to identify overall patterns among samples. Input datasets consisted of filtered CLR-transformed NTU data and $\log_2$-transformed copies per million KEGG KO data from both MG and MT. To balance the number of features across datasets, only the most variable features were retained for datasets with larger feature sets, following the *MOFA2* manual: 50% of features with the highest coefficient of variation (standard deviation divided by mean) were selected for MT NTU data and MG KEGG data, while all features were included for MG NTU data and MT KEGG data. MOFA was trained in "slow" mode with 8 factors, as recommended by the method for these datasets (not exceeding 9). Factors most associated with the exclusion treatment were selected for visualization.

To evaluate classification performance based on the MOFA ordination, the randomForest function in the *randomForest* R package v4.7-1.1 (Liaw and Wiener 2002) was used to train models (1,000 trees) for the exclusion treatment, snow treatment and vegetation cluster, using leave-one-out cross-validation. Class predictions for out-of-sample cases were evaluated using the confusionMatrix function in the *caret* R package v6.0-94 (Kuhn 2008) to obtain confusion matrices and balanced accuracies. Principal component analysis (PCA) was performed with the prcomp function in the *stats* R package included in the R v4.3.2 on filtered CLR-transformed NTU data and $\log_2$-transformed copies per million data.

*Differential abundance and expression analysis*

Linear mixed-effects models (LMMs) were used to identify DA (MG) and DE (MT) taxa, metabolic features (KO groups, marker genes), as well as differences in methane fluxes and pore water measurements in relation to the exclusion treatment (fence) and snow treatment, while accounting for vegetation effects. For each feature, two LMMs were fitted using the *lmerTest* R package v3.1-3 (Kuznetsova, Brockhoff, and Christensen 2017) with maximum likelihood estimation: (1) a base model including exclusion and snow treatment as fixed effects and vegetation cluster as a random effect and (2) a full model including the interaction between exclusion and snow treatment. A likelihood ratio test was used to compare the models; if the full model provided a significantly better fit, it was selected, otherwise the base model was retained.

For each selected model, p-values for exclusion, snow treatment and, where applicable, their interaction, were recorded. To control for false positives, p-values from all features within each analysis of large data sets (taxa, KO groups, for both DA and DE) were adjusted for false discovery rate (FDR) using the Benjamini–Hochberg procedure implemented in the p.adjust function of the *stats* R package v4.3.2 (R Core Team 2025). DA and DE features were determined using a FDR of ≤ 0.1, a commonly used cutoff in exploratory high-throughput omics studies to balance sensitivity and control of false discoveries under large multiple-testing burdens (Liao et al. 2014, Zhernakova et al. 2016, Barlow, Bogatyrev, and Ismagilov 2020, Zhang et al. 2021, Spottiswoode et al. 2026).

The filtered and CLR-transformed NTU data and $\log_2$-transformed copies per million data were used as input for the LMMs. The methane fluxes, pore water measurements and marker gene ratios, except the *nrfA* - (*nirS* + *nirK*) with negative values, were transformed using a natural logarithm for the LMM.

*Association of marker gene ratios, environmental variables and taxa*

Associations between marker gene abundances/expression/ratios and environmental variables were tested with simple linear regression using the lm function in the *stats* R package v4.3.2 (R Core Team 2025). $Log_2$-transformed values were used for marker gene abundances and expression and natural-log–transformed values for gene ratios and environmental measurements. Correlations between taxa abundance or expression and marker gene ratios were assessed with Pearson's correlation, using centered log-ratio (CLR)–transformed NTU data and natural-log–transformed ratio data.

*Gene set enrichment analysis*

Gene Set Enrichment Analysis (GSEA) (Subramanian *et al.* 2005) was used to examine the enrichment of functional KEGG modules within KO groups in the MG and MT datasets. A set of core prokaryotic metabolic modules (**Supplementary Table 24**) was selected as the target for enrichment testing for the exclusion and snow treatments. Module completeness was calculated using *anvi'o* development version 8 (v8-dev) (Eren *et al.* 2021) with the anvi-estimate-metabolism function (Veseli *et al.* 2025) (--include-kos-not-in-kofam --exclude-dashed-reactions --output-modes modules,module_steps,module_paths --add-copy-number). The KEGG module definitions were obtained via anvi-setup-kegg-data (snapshot v2025-02-04). Modules with ≥ 75% stepwise completeness were retained for enrichment testing.

Enrichment was assessed with the fgsea function in the *fgsea* R package v1.28.0 (Korotkevich *et al.* 2021). The input score for each KO group was calculated as (1 – p-value) from the DA or DE analysis LMMs, multiplied by the direction of change (model coefficient). P-values from the base LMM models were used and enrichment was evaluated separately for each treatment comparison (outside vs. inside, snow addition vs. snow control, snow removal vs. snow control).

<u>Figure licence information</u>

The world base map in Figure 1A is adapted from [Wikimedia Commons, File:BlankMap-World.svg] by [Canuckguy and many others], released under CC0 Public Domain Dedication and the Finland map is adapted from [Wikimedia Commons, File: Finland_administrative_divisions.svg] by [Fenn-O-maniC], licensed under Creative Commons Attribution-ShareAlike 4.0 International (CC BY-SA 4.0). Aerial imagery for Figure1B and Figure1C provided by the National Land Survey of Finland under the Creative Commons CC BY 4.0 license. All the adapted figures are licensed under CC BY-SA 4.0.
