## Supplementary Tables for "Soil microbiome structure and function reflect environmental variation rather than reindeer presence in a northern peatland"

**Supplementary Table 1**. Differentially expressed (false discovery rate ≤ 0.1) taxa in the metatranscriptomics taxonomic data at the order level associated with the exclusion treatment (outside vs. inside) according to the used linear mixed effects models.

| | Coefficient | P-value | FDR |
|---|---|---|---|
| *Clostridia UCG-014* | -0.674 | <0.001 | 0.011 |
| *KF-JG30-C25* | 1.464 | <0.001 | 0.011 |
| *Thiotrichales* | 1.552 | <0.001 | 0.064 |
| *Candidatus Staskawiczbacteria* | -0.383 | 0.001 | 0.064 |
| *Lachnospirales* | -0.557 | 0.001 | 0.067 |
| *Rhizobiales* | -0.286 | 0.001 | 0.091 |

**Supplementary Table 2**. Differentially expressed (false discovery rate ≤ 0.1) taxa in the metatranscriptomics taxonomic data at the order level associated with snow increase (snow increase vs. control) according to the used linear mixed effects models.

| | Coefficient | P-value | FDR |
|---|---|---|---|
| *Thermoflexales* | 0.344 | 0.001 | 0.089 |

**Supplementary Table 3.** Differentially expressed (false discovery rate ≤ 0.1) taxa in the metatranscriptomics taxonomic data at the order level associated with snow decrease (snow decrease vs. control) according to the used linear mixed effects models.

| | Coefficient | P-value | FDR |
|---|---|---|---|
| *Woesearchaeales* | -0.667 | <0.001 | 0.013 |
| *Omnitrophales* | -0.324 | <0.001 | 0.064 |
| *Candidatus Kerfeldbacteria* | -0.490 | <0.001 | 0.064 |
| *Thermoflexales* | 0.375 | 0.001 | 0.064 |
| *Frankiales* | 0.676 | 0.001 | 0.064 |
| *053A03-B-DI-P58* | -0.633 | 0.001 | 0.064 |
| *Spirochaetales* | -0.483 | 0.001 | 0.064 |
| *Candidatus Buchananbacteria* | -0.478 | 0.001 | 0.091 |

**Supplementary Table 4.** Differentially expressed (false discovery rate ≤ 0.1) taxa in the metatranscriptomics taxonomic data at the order level with significant interactions between the exclusion treatment and snow decrease according to the used linear mixed effects models.

| | Coefficient exclusion*snow increase | Coefficient exclusion*snow decrease | P-value exclusion*snow increase | P-value exclusion*snow decrease | FDR exclusion*snow increase | FDR exclusion*snow decrease |
|---|---|---|---|---|---|---|
| *1-20* | 0.296 | -0.713 | 0.134 | 0.001 | 0.608 | 0.064 |
| *Archaeoglobales* | 1.261 | 2.563 | 0.072 | 0.001 | 0.525 | 0.064 |
| *PB19* | 0.305 | -0.472 | 0.033 | 0.001 | 0.402 | 0.093 |

**Supplementary Table 5.** Differentially expressed (false discovery rate ≤ 0.1) taxa in the metatranscriptomics taxonomic data at the phylum level associated with the exclusion treatment (outside vs. inside) according to the used linear mixed effects models.

| | **Coefficient** | **P-value** | **FDR** |
|---|---|---|---|
| *Iainarchaeota* | -0.276 | <0.001 | 0.009 |
| *Firmicutes (A-Z)* | -0.369 | 0.001 | 0.035 |
| *SAR324* | -0.333 | 0.001 | 0.039 |

**Supplementary Table 6.** Differentially expressed (false discovery rate ≤ 0.1) taxa in the metatranscriptomics taxonomic data at the phylum level associated with snow decrease (snow decrease vs. control) according to the used linear mixed effects models.

| | **Coefficient** | **P-value** | **FDR** |
|---|---|---|---|
| *Margulisbacteria* | -0.720 | <0.001 | 0.024 |
| *Nanoarchaeota* | -0.497 | <0.001 | 0.026 |
| *Actinomycetota* | 0.532 | <0.001 | 0.026 |
| *FCPU426* | -0.372 | 0.003 | 0.081 |
| *Aquificota* | 0.453 | 0.003 | 0.081 |
| *Firestonebacteria* | -0.758 | 0.004 | 0.098 |

**Supplementary Table 7.** Differentially expressed (false discovery rate ≤ 0.1) taxa in the metatranscriptomics taxonomic data at the phylum level with significant interactions between the exclusion treatment and snow decrease according to the used linear mixed effects models.

| | **Coefficient exclusion*snow increase** | **Coefficient exclusion*snow decrease** | **P-value exclusion*snow increase** | **P-value exclusion*snow decrease** | **FDR exclusion*snow increase** | **FDR exclusion*snow decrease** |
|---|---|---|---|---|---|---|
| *Abditibacteriota* | -0.430 | -0.993 | 0.099 | <0.001 | 0.505 | 0.026 |

**Supplementary Table 8.** Differentially abundant (false discovery rate ≤ 0.1) KEGG orthology (KO) groups in the metagenomics KEGG KO data associated with the exclusion treatment (outside vs. inside) according to the used linear mixed effects models.

| | **Coefficient** | **P-value** | **FDR** |
|---|---|---|---|
| K06181; rluE; 23S rRNA pseudouridine2457 synthase [EC:5.4.99.20] | 0.222 | <0.001 | 0.006 |
| K19003; mgdA; 1,2-diacylglycerol 3-beta-glucosyltransferase [EC:2.4.1.336] | 0.372 | <0.001 | 0.044 |
| K09014; sufB; Fe-S cluster assembly protein SufB | 0.100 | <0.001 | 0.044 |
| K14441; rimO; ribosomal protein S12 methylthiotransferase [EC:2.8.4.4] | 0.087 | <0.001 | 0.044 |
| K02986; RP-S4, NAM9, rpsD; small subunit ribosomal protein S4 | 0.109 | <0.001 | 0.044 |
| K02484; K02484; two-component system, OmpR family, sensor kinase [EC:2.7.13.3] | 0.153 | <0.001 | 0.050 |
| K18479; yihS; sulfoquinovose isomerase [EC:5.3.1.31] | 0.433 | <0.001 | 0.050 |
| K03090; sigB; RNA polymerase sigma-B factor | 0.396 | <0.001 | 0.050 |
| K10544; xylH; D-xylose transport system permease protein | 0.248 | <0.001 | 0.050 |
| K23536; nupC; general nucleoside transport system permease protein | 0.187 | <0.001 | 0.050 |
| K06168; miaB; tRNA-2-methylthio-N6-dimethylallyladenosine synthase [EC:2.8.4.3] | 0.086 | <0.001 | 0.050 |
| K09015; sufD; Fe-S cluster assembly protein SufD | 0.101 | <0.001 | 0.050 |
| K21471; cwlO; peptidoglycan DL-endopeptidase CwlO [EC:3.4.-.-] | 0.286 | <0.001 | 0.058 |
| K16329; psuG; pseudouridylate synthase [EC:4.2.1.70] | 0.190 | <0.001 | 0.063 |
| K08093; hxlA; 3-hexulose-6-phosphate synthase [EC:4.1.2.43] | 0.295 | <0.001 | 0.066 |
| K02871; RP-L13, MRPL13, rplM; large subunit ribosomal protein L13 | 0.085 | <0.001 | 0.089 |
| K00331; nuoB; NADH-quinone oxidoreductase subunit B [EC:7.1.1.2] | 0.080 | <0.001 | 0.099 |
| K20327; xagB; glycosyltransferase XagB | 0.405 | <0.001 | 0.099 |
| K18955; whiB1_2_3_4; WhiB family transcriptional regulator, redox-sensing transcriptional regulator | 0.436 | <0.001 | 0.099 |
| K03867; BGLUT; UDP-glucose:tetrahydrobiopterin glucosyltransferase [EC:2.4.1.-] | 0.627 | <0.001 | 0.099 |
| K09013; sufC; Fe-S cluster assembly ATP-binding protein | 0.124 | <0.001 | 0.099 |
| K07695; devR; two-component system, NarL family, response regulator DevR | 0.310 | <0.001 | 0.099 |
| K00339; nuoJ; NADH-quinone oxidoreductase subunit J [EC:7.1.1.2] | 0.084 | <0.001 | 0.099 |
| K03076; secY; preprotein translocase subunit SecY | 0.092 | <0.001 | 0.099 |
| K01609; trpC; indole-3-glycerol phosphate synthase [EC:4.1.1.48] | 0.087 | <0.001 | 0.099 |
| K00423; E1.10.3.3; L-ascorbate oxidase [EC:1.10.3.3] | -0.477 | <0.001 | 0.099 |

**Supplementary Table 9.** Differentially expressed (false discovery rate ≤ 0.1) KEGG orthology (KO) groups in the metatranscriptomics KEGG KO data associated with the exclusion treatment (outside vs. inside) according to the used linear mixed effects models.

| | **Coefficient** | **P-value** | **FDR** |
|---|---|---|---|
| K00548; metH, MTR; 5-methyltetrahydrofolate--homocysteine methyltransferase [EC:2.1.1.13] | 0.255 | <0.001 | 0.032 |

**Supplementary Table 10.** Differentially abundant (false discovery rate ≤ 0.1) KEGG orthology (KO) groups in the metagenomics KEGG KO data associated with snow decrease (outside vs. inside) according to the used linear mixed effects models.

| | Coefficient | P-value | FDR |
|---|---|---|---|
| K03462; NAMPT; nicotinamide phosphoribosyltransferase [EC:2.4.2.12] | -0.555 | <0.001 | 0.044 |

**Supplementary Table 11.** Differentially expressed (false discovery rate ≤ 0.1) KEGG orthology (KO) groups in the metatranscriptomics KEGG KO data associated with snow decrease (snow decrease vs. control) according to the used linear mixed effects models.

| | Coefficient | P-value | FDR |
|---|---|---|---|
| K13195; CIRBP; cold-inducible RNA-binding protein | -0.706 | <0.001 | 0.032 |

**Supplementary Table 12**. The enriched KEGG modules in metagenomics data associated with the exclusion treatment (outside vs. inside) according to gene set enrichment analysis (GSEA) performed on the direction-adjusted significance scores (1 – p-value) from the used linear mixed effects models.

| Module | P-value | BH adjusted P-value | Log2err | Enrichment score | Normalized enrichment score | Module size |
|---|---|---|---|---|---|---|
| M00529_Denitrification, nitrate => nitrogen | <0.001 | 0.012 | 0.498 | -0.712 | -2.459 | 10 |
| M00567_Methanogenesis, CO2 => methane | 0.001 | 0.025 | 0.455 | 0.483 | 1.856 | 29 |

**Supplementary Table 13**. The enriched KEGG modules in metagenomics data associated with snow increase (snow increase vs. control)) according to gene set enrichment analysis (GSEA) performed on the direction-adjusted significance scores (1 – p-value) from the used linear mixed effects models.

| Module | P-value | BH adjusted P-value | Log2err | Enrichment score | Normalized enrichment score | Module size |
|---|---|---|---|---|---|---|
| M00174_Methane oxidation, methanotroph, methane => formaldehyde | <0.001 | <0.001 | 0.611 | -0.847 | -2.235 | 10 |
| M00804_Complete nitrification, comammox, ammonia => nitrite => nitrate | 0.001 | 0.019 | 0.455 | -0.824 | -1.844 | 6 |
| M00378_F420 biosynthesis, archaea | 0.001 | 0.019 | 0.455 | 0.890 | 1.950 | 5 |
| M00567_Methanogenesis, CO2 => methane | 0.002 | 0.025 | 0.432 | 0.478 | 1.787 | 29 |
| M00935_Methanofuran biosynthesis | 0.004 | 0.037 | 0.407 | 0.786 | 1.816 | 6 |
| M00596_Dissimilatory sulfate reduction, sulfate => H2S | 0.009 | 0.063 | 0.381 | -0.649 | -1.768 | 11 |
| M00528_Nitrification, ammonia => nitrite | 0.011 | 0.069 | 0.381 | -0.834 | -1.631 | 4.000 |

**Supplementary Table 14**. The enriched KEGG modules in metagenomics data associated with snow decrease (snow decrease vs. control)) according to gene set enrichment analysis (GSEA) performed on the direction-adjusted significance scores (1 – p-value) from the used linear mixed effects models.

| Module | P-value | BH adjusted P-value | Log2err | Enrichment score | Normalized enrichment score | Module size |
|---|---|---|---|---|---|---|
| M00378_F420 biosynthesis, archaea | 0.001 | 0.027 | 0.477 | 0.884 | 2.036 | 5 |
| M00567_Methanogenesis, CO2 => methane | 0.001 | 0.027 | 0.455 | 0.476 | 1.934 | 29 |
| M00009_Citrate cycle (TCA cycle, Krebs cycle) | 0.005 | 0.070 | 0.407 | -0.480 | -1.731 | 32 |
| M00002_Glycolysis, core module involving three-carbon compounds | 0.006 | 0.071 | 0.407 | -0.655 | -1.775 | 11.000 |
| M00308_Semi-phosphorylative Entner-Doudoroff pathway, gluconate => glycerate-3P | 0.008 | 0.072 | 0.381 | -0.744 | -1.652 | 6.000 |

**Supplementary Table 15**. The enriched KEGG modules in metatranscriptomics data associated with the exclusion treatment (outside vs. inside) according to gene set enrichment analysis (GSEA) performed on the direction-adjusted significance scores (1 – p-value) from the used linear mixed effects models.

| Module | P-value | BH adjusted P-value | Log2err | Enrichment score | Normalized enrichment score | Module size |
|---|---|---|---|---|---|---|
| M00579_Phosphate acetyltransferase-acetate kinase pathway, acetyl-CoA => acetate | 0.001 | 0.033 | 0.477 | -0.894 | -1.975 | 4 |
| M00003_Gluconeogenesis, oxaloacetate => fructose-6P | 0.002 | 0.033 | 0.455 | -0.502 | -2.025 | 19 |
| M00374_Dicarboxylate-hydroxybutyrate cycle | 0.005 | 0.046 | 0.407 | -0.490 | -1.876 | 17 |
| M00529_Denitrification, nitrate => nitrogen | 0.005 | 0.046 | 0.407 | -0.707 | -2.050 | 8 |
| M00567_Methanogenesis, CO2 => methane | 0.006 | 0.046 | 0.407 | 0.472 | 1.706 | 27 |
| M00307_Pyruvate oxidation, pyruvate => acetyl-CoA | 0.007 | 0.046 | 0.407 | -0.587 | -1.828 | 10.000 |
| M00173_Reductive citrate cycle (Arnon-Buchanan cycle) | 0.018 | 0.098 | 0.352 | -0.347 | -1.614 | 31.000 |

**Supplementary Table 16**. The enriched KEGG modules in metatranscriptomics data associated with snow increase (snow increase vs. control)) according to gene set enrichment analysis (GSEA) performed on the direction-adjusted significance scores (1 – p-value) from the used linear mixed effects models.

| Module | P-value | BH adjusted P-value | Log2err | Enrichment score | Normalized enrichment score | Module size |
|---|---|---|---|---|---|---|
| M00804_Complete nitrification, comammox, ammonia => nitrite => nitrate | <0.001 | 0.001 | 0.557 | -0.902 | -2.052 | 6 |
| M00358_Coenzyme M biosynthesis | <0.001 | 0.007 | 0.498 | 0.974 | 1.757 | 3 |
| M00528_Nitrification, ammonia => nitrite | 0.002 | 0.020 | 0.455 | -0.901 | -1.789 | 4 |
| M00165_Reductive pentose phosphate cycle (Calvin cycle) | 0.008 | 0.076 | 0.407 | -0.564 | -1.805 | 16 |

**Supplementary Table 17**. The enriched KEGG modules in metatranscriptomics data associated with snow decrease (snow decrease vs. control)) according to gene set enrichment analysis (GSEA) performed on the direction-adjusted significance scores (1 – p-value) from the used linear mixed effects models.

| Module | P-value | BH adjusted P-value | Log2err | Enrichment score | Normalized enrichment score | Module size |
|---|---|---|---|---|---|---|
| M00358_Coenzyme M biosynthesis | 0.001 | 0.032 | 0.477 | 0.954 | 1.760 | 3 |

**Supplementary Table 18**. The summary statistics of metabolic marker gene ratios associated with methane oxidation to methanogenesis (*pmoa* / *mcrA*), ammonification to denitrification (*nrfA* - (*nirK* + *nirS*) and $N_2O$ production to $N_2O$ reduction (*nirK* + *nirS*) / *nosZ* in metagenomics and metatranscriptomics data.

| | Metagenomics | | | Metatranscriptomics | | |
|---|---|---|---|---|---|---|
| | ***mcrA / pmoA*** | ***nrfA - (nirK + nirS)*** | ***(nirK + nirS) / nosZ*** | ***mcrA / pmoA*** | ***nrfA - (nirK + nirS)*** | ***(nirK + nirS) / nosZ*** |
| **Min.** | 0.535 | -38.173 | 1.978 | 0.165 | -153.681 | 0.946 |
| **1st Qu.** | 1.773 | -28.407 | 2.216 | 0.817 | -72.478 | 1.962 |
| **Median** | 4.267 | -16.557 | 2.386 | 2.113 | -50.183 | 2.554 |
| **Mean** | 9.028 | -15.989 | 2.490 | 26.158 | -55.014 | 2.920 |
| **3rd Qu.** | 11.274 | -5.858 | 2.658 | 11.845 | -32.646 | 3.253 |
| **Max.** | 55.173 | 43.175 | 3.511 | 684.795 | 2.282 | 8.024 |

**Supplementary Table 19**. The distribution of vegetation clusters across the exclusion treatment and the snow manipulation treatments.

| | *Trichophorum cespitosum* | *Carex chordorrhiza* | *Carex rostrata* |
|---|---|---|---|
| **Inside exclosure** | 14 | 4 | 0 |
| **Outside exclosure** | 6 | 2 | 10 |
| **Snow ambient (AMB)** | 8 | 2 | 2 |
| **Snow addition (+S)** | 5 | 3 | 4 |
| **Snow removal (-S)** | 7 | 1 | 4 |

**Supplementary Table 20**. The summary statistics for key variables for the 113 medium- and high-quality metagenome assembled genomes (MAGs) in the Puukkosuo metagenomics data.

| | Minimum | First quartile | Median | Mean | Third quartile | Maximum |
|---|---|---|---|---|---|---|
| Total length | 518458 | 1920180 | 2809684 | 2781683.0 | 3328616 | 6978053 |
| Number of contigs | 108 | 417 | 534 | 560.5 | 668 | 1441 |
| N50 of the contigs | 3200 | 3961 | 4667 | 6069.6 | 5851 | 59884 |
| GC content | 38.9 | 55.1 | 60.3 | 59.4 | 64.6 | 70.5 |
| Completion (%) | 50.7 | 57.7 | 67.6 | 69.5 | 78.9 | 98.6 |
| Redundandy (%) | 0 | 1.4 | 4.2 | 4.6 | 7.0 | 9.9 |

**Supplementary Table 21.** Individual metrics and taxonomic assignments for the 113 medium- and high-quality metagenome assembled genomes (MAGs) in the Puukkosuo metagenomics data. Metagenomic reads were co-assembled separately plots outside and inside the exclosure with MEGAHIT, processed in anvi’o, and binned with MetaBAT2, followed by manual refinement to MIMAG standards (≥ 50 % completeness, < 10 % redundancy). MAGs were dereplicated using FastANI, and taxonomy was assigned with GTDB-Tk.

| MAG name; Phylum; Order; Genus | Total length | Number of contigs | N50 of the contigs | GC content | Completion (%) | Redundancy (%) | Domain | Phylum | Class | Order | Family | Genus | Species |
|---|---|---|---|---|---|---|---|---|---|---|---|---|---|
| MAG1; Nitrospirota; Thermodesulfovibrionales; GW-Nitrospira-1 | 1715042 | 335 | 5445 | 47.2 | 77.5 | 5.6 | Bacteria | Nitrospirota | Thermodesulfovibrionia | Thermodesulfovibrionales | UBA6898 | GW-Nitrospira-1 | GW-Nitrospira-1 sp002839535 |
| MAG2; Electryoneota; RPQS01; | 1369625 | 350 | 3976 | 57.5 | 52.1 | 0.0 | Bacteria | Bacillota_A | Clostridia | Lachnospirales | Lachnospiraceae | RGIG7926 | RGIG7926 sp016282595 |
| MAG3; Nitrospirota; Thermodesulfovibrionales; UBA6898 | 2930882 | 280 | 17269 | 49.5 | 91.5 | 7.0 | Bacteria | Nitrospirota | Thermodesulfovibrionia | Thermodesulfovibrionales | UBA6898 | UBA6898 | UBA6898 sp016217675 |
| MAG4; Chloroflexota; Anaerolineales; DYJB01 | 2664205 | 595 | 4593 | 66.1 | 64.8 | 4.2 | Bacteria | Chloroflexota | Anaerolineae | Anaerolineales | E44-bin32 | DYJB01 | DYJB01 sp020723365 |
| MAG5; Verrucomicrobiota; Chthoniobacterales; Terrimicrobium | 2522620 | 604 | 4255 | 58.4 | 71.8 | 4.2 | Bacteria | | | | | | |
| MAG6; Pseudomonadota; Burkholderiales; | 2475112 | 534 | 4865 | 59.7 | 76.1 | 0.0 | Bacteria | Pseudomonadota | Gammaproteobacteria | Burkholderiales | Usitatibacteraceae | JAEUMY01 | JAEUMY01 sp016791265 |
| MAG7; Pseudomonadota; Pseudomonadales; SZUA-521 | 2010477 | 481 | 4063 | 61.6 | 56.3 | 1.4 | Bacteria | | | | | | |
| MAG8; Actinomycetota; IMCC26256; JALHSW01 | 1266938 | 333 | 3724 | 69.3 | 57.7 | 8.5 | Bacteria | Actinomycetota | Acidimicrobiia | IMCC26256 | PALSA-555 | JALHSW01 | JALHSW01 sp022865365 |
| MAG9; Acidobacteriota; Fen-336; | 1879369 | 514 | 3584 | 67.7 | 66.2 | 2.8 | Bacteria | | | | | | |
| MAG10; Methylomirabilota; Rokubacteriales; AR37 | 3839230 | 790 | 5085 | 67.5 | 91.5 | 5.6 | Bacteria | Methylomirabilota | Methylomirabilia | Rokubacteriales | CSP1-6 | AR37 | AR37 sp016179165 |
| MAG11; Actinomycetota; UBA4738; AC-51 | 1457159 | 375 | 3858 | 68.2 | 64.8 | 2.8 | Bacteria | | | | | | |
| MAG12; Pseudomonadota; Burkholderiales; | 1867187 | 469 | 3931 | 63.1 | 62.0 | 8.5 | Bacteria | Pseudomonadota | Gammaproteobacteria | Burkholderiales | | | |
| MAG13; Bacteroidota; Bacteroidales; LD21 | 1896446 | 473 | 4094 | 41.4 | 50.7 | 0.0 | Bacteria | Bacteroidota | Bacteroidia | Bacteroidales | VadinHA17 | LD21 | LD21 sp003520925 |
| MAG14; Pseudomonadota; Burkholderiales; Sideroxyarcus | 1738360 | 433 | 4012 | 55.6 | 64.8 | 4.2 | Bacteria | Pseudomonadota | Gammaproteobacteria | Burkholderiales | Gallionellaceae | Sideroxyarcus | |
| MAG15; Eisenbacteria; RBG-16-71-46; WS-11 | 711033 | 206 | 3320 | 68.0 | 53.5 | 0.0 | Bacteria | Eisenbacteria | RBG-16-71-46 | RBG-16-71-46 | RBG-16-71-46 | WS-11 | WS-11 sp005893365 |
| MAG16; Pseudomonadota; Dongiales; | 2809684 | 646 | 4356 | 63.5 | 56.3 | 9.9 | Bacteria | | | | | | |
| MAG17; Pseudomonadota; Burkholderiales; | 3370801 | 607 | 5851 | 60.3 | 94.4 | 4.2 | Bacteria | Pseudomonadota | Gammaproteobacteria | Burkholderiales | Usitatibacteraceae | JAEUMY01 | JAEUMY01 sp016791265 |
| MAG18; Methylomirabilota; Rokubacteriales; | 2730710 | 718 | 3676 | 67.7 | 54.9 | 1.4 | Bacteria | | | | | | |
| MAG19; Nitrospirota; Nitrospirales; Palsa-1315 | 3474959 | 323 | 17969 | 56.7 | 95.8 | 1.4 | Bacteria | Nitrospirota | Nitrospiria | Nitrospirales | Nitrospiraceae | Palsa-1315 | Palsa-1315 sp003135435 |
| MAG20; Chloroflexota; Anaerolineales; VGNF01 | 3127218 | 633 | 5229 | 55.1 | 77.5 | 2.8 | Bacteria | Chloroflexota | Anaerolineae | Anaerolineales | EnvOPS12 | OLB14 | OLB14 sp016789305 |
| MAG21; Nitrospirota; UBA9217; JAJYKN01 | 2227524 | 471 | 4796 | 53.4 | 67.6 | 2.8 | Bacteria | Nitrospirota | UBA9217 | UBA9217 | UBA9217 | JAJYKN01 | |
| MAG22; Nitrospirota; Nitrospirales; Palsa-1315 | 3094438 | 447 | 8403 | 57.1 | 80.3 | 0.0 | Bacteria | Nitrospirota | Nitrospiria | Nitrospirales | Nitrospiraceae | Palsa-1315 | |
| MAG23; Pseudomonadota; Burkholderiales; SG8-41 | 3032004 | 642 | 4771 | 64.4 | 93.0 | 4.2 | Bacteria | Pseudomonadota | Gammaproteobacteria | Burkholderiales | SG8-41 | SG8-41 | SG8-41 sp001771935 |
| MAG24; Pseudomonadota; Rhizobiales; Methyloceanibacter | 1481764 | 377 | 3905 | 63.5 | 52.1 | 2.8 | Bacteria | Pseudomonadota | Alphaproteobacteria | Rhizobiales | Methyloligellaceae | Methyloceanibacter | |
| MAG25; Pseudomonadota; Rhizobiales; Methylocystis | 3840714 | 593 | 7862 | 61.5 | 93.0 | 0.0 | Bacteria | Pseudomonadota | Alphaproteobacteria | Rhizobiales | Beijerinckiaceae | Methylocystis | Methylocystis sp021731785 |
| MAG26; Nitrospirota; UBA9217; JAJYKN01 | 2144969 | 534 | 3981 | 57.4 | 62.0 | 9.9 | Bacteria | Nitrospirota | UBA9217 | UBA9217 | UBA9217 | JAJYKN01 | JAJYKN01 sp021788515 |
| MAG27; Nitrospirota; Nitrospirales; Palsa-1315 | 2843453 | 375 | 10165 | 56.3 | 87.3 | 4.2 | Bacteria | Nitrospirota | Nitrospiria | Nitrospirales | Nitrospiraceae | Palsa-1315 | |
| MAG28; Desulfobacterota; Desulfomonilales; | 3873733 | 951 | 4080 | 53.1 | 57.7 | 8.5 | Bacteria | | | | | | |
| MAG29; Chloroflexota; Anaerolineales; Defluviilinea | 1482676 | 387 | 3799 | 53.2 | 60.6 | 0.0 | Bacteria | Chloroflexota | Anaerolineae | Anaerolineales | EnvOPS12 | UBA12294 | UBA12294 sp013388855 |
| MAG30; Pseudomonadota; Burkholderiales; JALORV01 | 3282903 | 672 | 5095 | 63.4 | 70.4 | 4.2 | Bacteria | Pseudomonadota | Gammaproteobacteria | Burkholderiales | Burkholderiaceae_A | JACCZX01 | JACCZX01 sp013696665 |
| MAG31; Methylomirabilota; Rokubacteriales; AR37 | 4429943 | 1044 | 4372 | 67.3 | 54.9 | 7.0 | Bacteria | Methylomirabilota | Methylomirabilia | Rokubacteriales | CSP1-6 | AR37 | AR37 sp016179165 |
| MAG32; Desulfobacterota; Geobacterales; CAIPTY01 | 3311607 | 696 | 4956 | 56.5 | 70.4 | 5.6 | Bacteria | Desulfobacterota_F | Desulfuromonadia | Geobacterales | Pseudopelobacteraceae | CAIPTY01 | CAIPTY01 sp016720395 |
| MAG33; Chloroflexota; Anaerolineales; Defluviilinea | 1920180 | 444 | 4326 | 56.7 | 60.6 | 4.2 | Bacteria | | | | | | |
| MAG34; Bacteroidota; Bacteroidales; LD21 | 2540130 | 673 | 3615 | 39.7 | 53.5 | 8.5 | Bacteria | Bacteroidota | Bacteroidia | Bacteroidales | VadinHA17 | LD21 | |
| MAG35; Myxococcota_A; UBA9160; PR03 | 1559847 | 410 | 3704 | 67.8 | 59.2 | 0.0 | Bacteria | Myxococcota_A | UBA9160 | UBA9160 | PR03 | PR03 | PR03 sp018262315 |
| MAG36; Actinomycetota; Acidimicrobiales; JAENVT01 | 3931835 | 780 | 5382 | 64.5 | 77.5 | 4.2 | Bacteria | Actinomycetota | Acidimicrobiia | Acidimicrobiales | Ilumatobacteraceae | JAENVT01 | JAENVT01 sp016650435 |
| MAG37; Myxococcota; Myxococcales; Anaeromyxobacter | 2059703 | 454 | 4667 | 70.5 | 64.8 | 1.4 | Bacteria | Myxococcota | Myxococcia | Myxococcales | Anaeromyxobacteraceae | | |
| MAG38; Actinomycetota; IMCC26256; | 1518780 | 442 | 3333 | 68.5 | 50.7 | 9.9 | Bacteria | Actinomycetota | Acidimicrobiia | IMCC26256 | PALSA-555 | | |
| MAG39; Pseudomonadota; Burkholderiales; SG8-41 | 3016676 | 622 | 5108 | 64.4 | 90.1 | 5.6 | Bacteria | Pseudomonadota | Gammaproteobacteria | Burkholderiales | | | |
| MAG40; Methylomirabilota; Methylomirabilales; 2-02-FULL-66-22 | 5618893 | 1030 | 6020 | 64.1 | 87.3 | 2.8 | Bacteria | | | | | | |
| MAG41; Desulfobacterota_B; UBA9968; DP-20 | 2842652 | 668 | 4339 | 57.3 | 74.6 | 5.6 | Bacteria | Desulfobacterota_B | Binatia | UBA9968 | UBA9968 | DP-20 | DP-20 sp945889495 |
| MAG42; Pseudomonadota; Rhizobiales; Rhodoplanes | 2931872 | 610 | 5049 | 64.6 | 74.6 | 5.6 | Bacteria | Pseudomonadota | Alphaproteobacteria | Rhizobiales | Xanthobacteraceae | Rhodoplanes | Rhodoplanes sp016793685 |
| MAG43; Pseudomonadota; Steroidobacterales; CADEED01 | 2014806 | 351 | 6385 | 64.3 | 78.9 | 1.4 | Bacteria | Pseudomonadota | Gammaproteobacteria | | | | |
| MAG44; Desulfobacterota_B; DP-6; | 1717869 | 497 | 3297 | 68.2 | 50.7 | 1.4 | Bacteria | | | | | | |
| MAG45; Desulfobacterota_B; UBA9968; DP-1 | 4216331 | 687 | 6878 | 54.2 | 90.1 | 5.6 | Bacteria | Desulfobacterota_B | Binatia | UBA9968 | UBA9968 | DP-1 | DP-1 sp020027655 |
| MAG46; Actinomycetota; UBA5794; JALYNX01 | 1499894 | 381 | 3910 | 62.7 | 67.6 | 4.2 | Bacteria | | | | | | |
| MAG47; Chloroflexota; Anaerolineales; VGNF01 | 2947147 | 603 | 5156 | 56.5 | 78.9 | 1.4 | Bacteria | Chloroflexota | Anaerolineae | Anaerolineales | EnvOPS12 | OLB14 | |
| MAG48; Verrucomicrobiota; Chthoniobacterales; Terrimicrobium | 2468182 | 627 | 3871 | 58.4 | 62.0 | 5.6 | Bacteria | | | | | | |
| MAG49; Actinomycetota; Acidimicrobiales; JAENVT01 | 2973888 | 656 | 4673 | 65.6 | 84.5 | 5.6 | Bacteria | Actinomycetota | Acidimicrobiia | Acidimicrobiales | Ilumatobacteraceae | JAENVT01 | JAENVT01 sp016650435 |
| MAG50; Methylomirabilota; Rokubacteriales; AR37 | 2516603 | 467 | 6060 | 67.6 | 67.6 | 8.5 | Bacteria | Methylomirabilota | Methylomirabilia | Rokubacteriales | CSP1-6 | AR37 | AR37 sp016179165 |
| MAG51; Pseudomonadota; Burkholderiales; 2-12-FULL-64-23 | 4826209 | 786 | 7191 | 63.1 | 97.2 | 0.0 | Bacteria | Pseudomonadota | Gammaproteobacteria | Burkholderiales | SG8-39 | 2-12-FULL-64-23 | |
| MAG52; Chloroflexota; UBA4142; | 575945 | 173 | 3200 | 53.0 | 50.7 | 2.8 | Bacteria | | | | | | |
| MAG53; Chloroflexota; Anaerolineales; Defluviilinea | 3264859 | 450 | 8684 | 53.9 | 76.1 | 4.2 | Bacteria | Chloroflexota | Anaerolineae | Anaerolineales | EnvOPS12 | UBA12294 | UBA12294 sp016932715 |
| MAG54; Methylomirabilota; Rokubacteriales; CAMLFJ01 | 1526534 | 427 | 3494 | 68.0 | 50.7 | 4.2 | Bacteria | Methylomirabilota | Methylomirabilia | Rokubacteriales | CSP1-6 | AR37 | AR37 sp003220345 |
| MAG55; Pseudomonadota; Burkholderiales; 2-12-FULL-64-23 | 3924542 | 645 | 7136 | 63.0 | 87.3 | 7.0 | Bacteria | Pseudomonadota | Gammaproteobacteria | Burkholderiales | SG8-39 | 2-12-FULL-64-23 | |
| MAG56; Chloroflexota; Anaerolineales; Villigracilis | 3064206 | 604 | 5588 | 48.9 | 69.0 | 1.4 | Bacteria | Chloroflexota | Anaerolineae | Anaerolineales | EnvOPS12 | OLB14 | OLB14 sp014379595 |
| MAG57; Nitrospirota; UBA9217; JAAXXJ01 | 2893875 | 440 | 7812 | 58.3 | 76.1 | 4.2 | Bacteria | Nitrospirota | UBA9217 | UBA9217 | UBA9217 | JAAXXJ01 | JAAXXJ01 sp017885365 |
| MAG58; Nitrospirota; Nitrospirales; Palsa-1315 | 3328616 | 223 | 25662 | 56.1 | 98.6 | 0.0 | Bacteria | Nitrospirota | Nitrospiria | Nitrospirales | Nitrospiraceae | Palsa-1315 | |
| MAG59; Bacteroidota; Bacteroidales; LD21 | 3233407 | 485 | 7991 | 38.9 | 77.5 | 1.4 | Bacteria | Bacteroidota | Bacteroidia | Bacteroidales | VadinHA17 | LD21 | |
| MAG60; Desulfobacterota_E; Deferrimicrobiales; Deferrimicrobium | 1632087 | 351 | 4998 | 66.7 | 69.0 | 9.9 | Bacteria | Desulfobacterota_E | Deferrimicrobia | Deferrimicrobiales | Deferrimicrobiaceae | Deferrimicrobium | |
| MAG61; Chloroflexota; Anaerolineales; Defluviilinea | 1179682 | 339 | 3351 | 53.3 | 50.7 | 7.0 | Bacteria | Chloroflexota | Anaerolineae | Anaerolineales | EnvOPS12 | UBA12294 | UBA12294 sp013388855 |
| MAG62; Pseudomonadota; Rhizobiales; Methyloceanibacter | 2982805 | 834 | 3402 | 62.6 | 53.5 | 8.5 | Bacteria | Pseudomonadota | Alphaproteobacteria | Rhizobiales | Methyloligellaceae | Methyloceanibacter | |
| MAG63; Actinomycetota; UBA4738; AC-51 | 1452837 | 338 | 4288 | 68.6 | 62.0 | 1.4 | Bacteria | | | | | | |
| MAG64; Chloroflexota; Anaerolineales; Defluviilinea | 5092441 | 839 | 6830 | 53.4 | 76.1 | 8.5 | Bacteria | Chloroflexota | Anaerolineae | Anaerolineales | EnvOPS12 | | |
| MAG65; Chloroflexota; Anaerolineales; VGNF01 | 3479450 | 465 | 9228 | 51.0 | 81.7 | 2.8 | Bacteria | Chloroflexota | Anaerolineae | Anaerolineales | EnvOPS12 | OLB14 | |
| MAG66; Acidobacteriota; Pyrinomonadales; UBA11740 | 3364690 | 789 | 4331 | 56.6 | 50.7 | 8.5 | Bacteria | Acidobacteriota | Blastocatellia | Pyrinomonadales | Pyrinomonadaceae | UBA11740 | UBA11740 sp003168335 |
| MAG67; Nitrospirota; UBA9217; JAJYKN01 | 1984827 | 463 | 4271 | 53.4 | 60.6 | 4.2 | Bacteria | Nitrospirota | UBA9217 | UBA9217 | UBA9217 | JAJYKN01 | JAJYKN01 sp021788515 |
| MAG68; Pseudomonadota; Burkholderiales; JAEUMW01 | 1612268 | 436 | 3605 | 64.6 | 62.0 | 0.0 | Bacteria | Pseudomonadota | Gammaproteobacteria | Burkholderiales | JAEUMW01 | JAEUMW01 | JAEUMW01 sp016791285 |
| MAG69; Chloroflexota; Anaerolineales; UBA700 | 2644304 | 680 | 3863 | 54.1 | 67.6 | 1.4 | Bacteria | Chloroflexota | Anaerolineae | Anaerolineales | Anaerolineaceae | | |
| MAG70; Chloroflexota; Anaerolineales; VGNF01 | 3506739 | 582 | 7070 | 50.5 | 57.7 | 7.0 | Bacteria | Chloroflexota | Anaerolineae | Anaerolineales | EnvOPS12 | | |
| MAG71; Pseudomonadota; Rhizobiales; Hyphomicrobium_A | 3106484 | 653 | 4866 | 62.9 | 62.0 | 8.5 | Bacteria | Pseudomonadota | Alphaproteobacteria | Rhizobiales | Hyphomicrobiaceae | Hyphomicrobium_A | Hyphomicrobium_A album |
| MAG72; Bacteroidota; Bacteroidales; LD21 | 2899426 | 556 | 5490 | 41.0 | 69.0 | 9.9 | Bacteria | Bacteroidota | Bacteroidia | Bacteroidales | VadinHA17 | LD21 | LD21 sp003520925 |
| MAG73; Bacteroidota; Bacteroidales; LD21 | 3252435 | 399 | 9887 | 39.2 | 70.4 | 5.6 | Bacteria | Bacteroidota | Bacteroidia | Bacteroidales | VadinHA17 | LD21 | |
| MAG74; Nitrospirota; UBA9217; JALNZF01 | 1327174 | 325 | 4286 | 54.6 | 54.9 | 2.8 | Bacteria | Nitrospirota | UBA9217 | UBA9217 | UBA9217 | JALNZF01 | JALNZF01 sp023229435 |
| MAG75; Pseudomonadota; Burkholderiales; CAISUK01 | 2263784 | 519 | 4401 | 62.1 | 70.4 | 4.2 | Bacteria | Pseudomonadota | Gammaproteobacteria | Burkholderiales | Rhodocyclaceae | CAISUK01 | CAISUK01 sp903888645 |
| MAG76; Nitrospirota; Thermodesulfovibrionales; JAJYIF01 | 3253218 | 108 | 59884 | 49.9 | 94.4 | 1.4 | Bacteria | Nitrospirota | Thermodesulfovibrionia | Thermodesulfovibrionales | UBA9159 | JAMCQB01 | JAMCQB01 sp023382175 |
| MAG77; Chloroflexota; Anaerolineales; Villigracilis | 2469219 | 575 | 4252 | 49.1 | 53.5 | 5.6 | Bacteria | Chloroflexota | Anaerolineae | Anaerolineales | EnvOPS12 | OLB14 | |
| MAG78; Acidobacteriota; Fen-336; | 4600401 | 809 | 6582 | 68.2 | 87.3 | 8.5 | Bacteria | | | | | | |
| MAG79; Actinomycetota; Gaiellales; GMQP-bins7 | 1338404 | 400 | 3244 | 67.8 | 52.1 | 9.9 | Bacteria | Actinomycetota | Thermoleophilia | Gaiellales | Gaiellaceae | GMQP-bins7 | GMQP-bins7 sp021323315 |
| MAG80; Desulfobacterota_B; UBA9968; JACPFE01 | 3358669 | 717 | 5018 | 56.7 | 60.6 | 5.6 | Bacteria | Desulfobacterota_B | Binatia | UBA9968 | UBA9968 | DP-1 | DP-1 sp005879605 |
| MAG81; Pseudomonadota; Burkholderiales; | 4098068 | 634 | 7656 | 63.5 | 87.3 | 1.4 | Bacteria | Pseudomonadota | Gammaproteobacteria | | | | |
| MAG82; Desulfobacterota_B; HRBIN30; JAKLJW01 | 4445937 | 1101 | 4021 | 66.6 | 76.1 | 8.5 | Bacteria | Desulfobacterota_B | Binatia | HRBIN30 | JAGDMS01 | JAKLJW01 | JAKLJW01 sp023150935 |
| MAG83; Actinomycetota; Gaiellales; GMQP-bins7 | 1970417 | 417 | 4962 | 68.0 | 74.6 | 4.2 | Bacteria | Actinomycetota | Thermoleophilia | Gaiellales | Gaiellaceae | GMQP-bins7 | GMQP-bins7 sp021323315 |
| MAG84; Pseudomonadota; Rariloculales; FEN-1219 | 2320846 | 552 | 4315 | 65.2 | 64.8 | 1.4 | Bacteria | Pseudomonadota | Gammaproteobacteria | UBA6522 | UBA6522 | FEN-1219 | FEN-1219 sp003155415 |
| MAG85; Pseudomonadota; Rhizobiales; Rhodoplanes | 3662794 | 763 | 5074 | 63.6 | 59.2 | 2.8 | Bacteria | Pseudomonadota | Alphaproteobacteria | Rhizobiales | Xanthobacteraceae | Rhodoplanes | Rhodoplanes sp016793685 |
| MAG86; Pseudomonadota; Dongiales; | 4235489 | 548 | 9404 | 64.0 | 87.3 | 5.6 | Bacteria | | | | | | |
| MAG87; Acidobacteriota; Pyrinomonadales; JAOAPY01 | 6978053 | 672 | 13743 | 56.4 | 98.6 | 8.5 | Bacteria | Acidobacteriota | Blastocatellia | Pyrinomonadales | Pyrinomonadaceae | PSRF01 | |
| MAG88; Methylomirabilota; Rokubacteriales; | 4526313 | 1158 | 3868 | 67.1 | 54.9 | 9.9 | Bacteria | | | | | | |
| MAG89; Acidobacteriota; Fen-336; | 4620181 | 1054 | 4443 | 68.8 | 83.1 | 5.6 | Bacteria | | | | | | |
| MAG90; Acidobacteriota; Pyrinomonadales; PSRF01 | 6547371 | 733 | 11808 | 55.1 | 93.0 | 4.2 | Bacteria | Acidobacteriota | Blastocatellia | Pyrinomonadales | Pyrinomonadaceae | PSRF01 | PSRF01 sp023717565 |
| MAG91; Bacteroidota; Bacteroidales; LD21 | 2914866 | 642 | 4706 | 41.9 | 80.3 | 5.6 | Bacteria | Bacteroidota | Bacteroidia | Bacteroidales | VadinHA17 | LD21 | LD21 sp003520925 |
| MAG92; Tectomicrobia; Entotheonellales; | 5099274 | 1441 | 3432 | 58.9 | 50.7 | 7.0 | Bacteria | | | | | | |
| MAG93; Nitrospirota; Thermodesulfovibrionales; GW-Nitrospira-1 | 1796401 | 362 | 5352 | 46.8 | 69.0 | 7.0 | Bacteria | Nitrospirota | Thermodesulfovibrionia | Thermodesulfovibrionales | UBA6898 | GW-Nitrospira-1 | GW-Nitrospira-1 sp002839535 |
| MAG94; Pseudomonadota; Competibacterales; CAIMUS01 | 1989426 | 513 | 3915 | 57.6 | 57.7 | 4.2 | Bacteria | Pseudomonadota | Gammaproteobacteria | Competibacterales | Competibacteraceae | CAIMUS01 | CAIMUS01 sp903844665 |
| MAG95; Tectomicrobia; J063; | 3300993 | 866 | 3752 | 58.7 | 59.2 | 2.8 | Bacteria | Bacillota_A | Clostridia | Clostridiales | Clostridiaceae | Clostridium | Clostridium sp012519155 |
| MAG96; Chloroflexota; Anaerolineales; Defluviilinea | 1699598 | 411 | 4220 | 57.2 | 64.8 | 0.0 | Bacteria | | | | | | |
| MAG97; Pseudomonadota; Rhizobiales; Hyphomicrobium_A | 3306998 | 689 | 5042 | 63.2 | 73.2 | 7.0 | Bacteria | Pseudomonadota | Alphaproteobacteria | Rhizobiales | Hyphomicrobiaceae | Hyphomicrobium_A | Hyphomicrobium_A sp009026145 |
| MAG98; Chloroflexota; Anaerolineales; VGNF01 | 2125303 | 513 | 4179 | 56.5 | 53.5 | 1.4 | Bacteria | Chloroflexota | Anaerolineae | Anaerolineales | EnvOPS12 | OLB14 | |
| MAG99; Pseudomonadota; Rhizobiales; Rhodoplanes | 3031342 | 685 | 4567 | 64.2 | 70.4 | 2.8 | Bacteria | Pseudomonadota | Alphaproteobacteria | Rhizobiales | Xanthobacteraceae | Rhodoplanes | Rhodoplanes sp016793685 |
| MAG100; Pseudomonadota; Burkholderiales; Fen-999 | 518458 | 154 | 3299 | 62.9 | 50.7 | 8.5 | Bacteria | Pseudomonadota | Gammaproteobacteria | Burkholderiales | Rhodocyclaceae | Fen-999 | Fen-999 sp003157035 |
| MAG101; Pseudomonadota; Burkholderiales; JALORV01 | 3517878 | 553 | 7554 | 63.4 | 87.3 | 9.9 | Bacteria | Pseudomonadota | Gammaproteobacteria | Burkholderiales | Burkholderiaceae_A | JACCZX01 | JACCZX01 sp013696665 |
| MAG102; Methylomirabilota; Rokubacteriales; AR37 | 1679482 | 314 | 5534 | 67.8 | 50.7 | 7.0 | Bacteria | Methylomirabilota | Methylomirabilia | Rokubacteriales | CSP1-6 | AR37 | AR37 sp016179165 |
| MAG103; Bacteroidota; UBA10030; UBA6906 | 1548479 | 387 | 3961 | 58.3 | 66.2 | 5.6 | Bacteria | Bacteroidota | UBA10030 | UBA10030 | UBA6906 | | |
| MAG104; Chloroflexota; Anaerolineales; DYJB01 | 2215591 | 528 | 4254 | 66.0 | 70.4 | 8.5 | Bacteria | Chloroflexota | Anaerolineae | Anaerolineales | E44-bin32 | DYJB01 | DYJB01 sp020723365 |
| MAG105; Pseudomonadota; Rhizobiales; Methyloceanibacter | 2241862 | 445 | 5505 | 60.9 | 80.3 | 7.0 | Bacteria | Pseudomonadota | Alphaproteobacteria | Rhizobiales | Methyloligellaceae | Methyloceanibacter | |
| MAG106; Chloroflexota; Anaerolineales; VGNF01 | 3249445 | 401 | 10147 | 55.5 | 93.0 | 5.6 | Bacteria | Chloroflexota | Anaerolineae | Anaerolineales | EnvOPS12 | OLB14 | |
| MAG107; Bacteroidota; UBA10030; UBA9647 | 3499185 | 749 | 4906 | 53.1 | 69.0 | 2.8 | Bacteria | Bacteroidota | UBA10030 | UBA10030 | UBA10030 | UBA9647 | UBA9647 sp003170475 |
| MAG108; Myxococcota; Haliangiales; | 2225521 | 575 | 3850 | 68.8 | 59.2 | 5.6 | Bacteria | | | | | | |
| MAG109; Desulfobacterota_B; HRBIN30; | 2091254 | 581 | 3508 | 66.8 | 52.1 | 1.4 | Bacteria | Desulfobacterota_B | Binatia | HRBIN30 | CADEER01 | JAHBUY01 | JAHBUY01 sp019637795 |
| MAG110; Pseudomonadota; Rhizobiales; Aestuariivirga | 2792052 | 516 | 6180 | 56.0 | 71.8 | 1.4 | Bacteria | Pseudomonadota | Alphaproteobacteria | Rhizobiales | Aestuariivirgaceae | Aestuariivirga | |
| MAG111; Nitrospirota; UBA9217; JAAXXJ01 | 2056699 | 501 | 4167 | 58.3 | 52.1 | 4.2 | Bacteria | Nitrospirota | UBA9217 | UBA9217 | UBA9217 | JAAXXJ01 | JAAXXJ01 sp017885365 |
| MAG112; Chloroflexota; Anaerolineales; Defluviilinea | 3041390 | 608 | 5370 | 52.9 | 77.5 | 5.6 | Bacteria | Chloroflexota | Anaerolineae | Anaerolineales | EnvOPS12 | UBA12294 | UBA12294 sp013388855 |
| MAG113; Methylomirabilota; Methylomirabilales; 2-02-FULL-66-22 | 2345599 | 573 | 4158 | 63.6 | 67.6 | 0.0 | hloroflexot | Anaerolineae | Anaerolineales | Villigracilaceae | Defluviilinea | | |

**Supplementary Table 22**. Additional manually identified and filtered eukaryotic proteins from the Compiled Greening Lab metabolic marker gene database.

| **Manually identified and supplemented eukaryotic proteins** |
|---|
| AclB-XP_003613199.1 - Medicago truncatula |
| SdhA_FrdA-Arabidopsis thaliana (Group 5c) |
| SdhA_FrdA-Ascaris suum (Group 5c) |
| SdhA_FrdA-Aspergillus niger (Group 5c) |
| SdhA_FrdA-Blattella germanic (Group 1c) |
| SdhA_FrdA-Caenorhabditis elegans (Group 5c) |
| SdhA_FrdA-Candida albicans (Group 5c) |
| SdhA_FrdA-Drosophila melanogaster (Group 5c) |
| SdhA_FrdA-Homo sapiens (Group 5c) |
| SdhA_FrdA-Neurospora crassa (Group 5c) |
| SdhA_FrdA-Penicillium chrysogenum (Group 5c) |
| SdhA_FrdA-Porphyra umbilicalis (Group 1a) |
| SdhA_FrdA-Saccharomyces cerevisiae (Group 5c) |
| SdhA_FrdA-Trichophyton rubrum (Group 5c) |
| SdhA_FrdA-Ustilago maydis (Group 5c) |
| NirK-XP_003067932.1 - Coccidioides posadasii |
| FeFe-XP_001318941.1 - Trichomonas vaginalis - [FeFe] Group A1 |
| Sqr-XP_029646137.1 - Octopus vulgaris - Sqr II |

**Supplementary Table 23.** Read mapping statistics for the metagenomics and metatrancriptomics data for the different metabolic functional and taxonomics alignments used in the study. The total number of reads for all samples are given in first column for metagenomics and metatranscriptomics data, respectively. The second column gives the proportion of trimmed reads passing the quality control. For metagenomics, all remaining columns give the proportion of quality controlled mate-pair combined read alignments to the trimmed reads. For metatranscriptomics, the trimmed reads are further filtered for ribosomal RNA (rRNA) given as proportions to the trimmed reads. For metatranscriptomics, all the KEGG and metabolic marker gene hits are given as proportions to the rRNA-filtered reads while the SILVA SSU rRNA counts are given as proportions to the trimmed reads.

**Metagenomics**

| Sample | Total read pairs | Total read pairs trimmed | Total KEGG hits | Total KEGG gene with KO | Total metabolic marker gene hits | Total SILVA SSU rRNA counts |
|---|---|---|---|---|---|---|
| P1 | 25167271 | 98.08 % | 49.24 % | 30.85 % | 0.71 % | 0.07 % |
| P2 | 13005673 | 92.27 % | 42.62 % | 27.09 % | 0.62 % | 0.05 % |
| P3 | 9579647 | 92.37 % | 41.44 % | 26.17 % | 0.54 % | 0.06 % |
| P4 | 15748479 | 91.83 % | 42.32 % | 26.56 % | 0.56 % | 0.05 % |
| P5 | 20364130 | 93.69 % | 43.20 % | 26.91 % | 0.58 % | 0.06 % |
| P6 | 17432323 | 97.11 % | 47.03 % | 29.52 % | 0.68 % | 0.07 % |
| P7 | 11665137 | 91.93 % | 43.61 % | 27.65 % | 0.63 % | 0.05 % |
| P8 | 17001416 | 92.99 % | 41.55 % | 26.39 % | 0.62 % | 0.05 % |
| P9 | 13533695 | 91.73 % | 42.93 % | 27.21 % | 0.63 % | 0.05 % |
| P10 | 12834923 | 92.68 % | 40.87 % | 25.75 % | 0.50 % | 0.06 % |
| P11 | 18398807 | 94.98 % | 44.80 % | 28.26 % | 0.64 % | 0.06 % |
| P12 | 15463174 | 92.01 % | 42.03 % | 26.74 % | 0.61 % | 0.06 % |
| P13 | 15687452 | 95.83 % | 45.97 % | 28.87 % | 0.62 % | 0.06 % |
| P14 | 16371329 | 92.54 % | 42.54 % | 26.70 % | 0.54 % | 0.05 % |
| P15 | 12709520 | 92.27 % | 43.28 % | 26.80 % | 0.56 % | 0.04 % |
| P16 | 12035466 | 91.24 % | 41.07 % | 25.87 % | 0.56 % | 0.05 % |
| P17 | 22564118 | 97.41 % | 48.38 % | 30.13 % | 0.69 % | 0.06 % |
| P18 | 15740660 | 94.19 % | 44.14 % | 27.60 % | 0.59 % | 0.05 % |
| P19 | 9518922 | 90.65 % | 42.48 % | 26.60 % | 0.56 % | 0.05 % |
| P20 | 14856433 | 94.77 % | 43.82 % | 27.61 % | 0.66 % | 0.07 % |
| P21 | 20442272 | 97.03 % | 47.33 % | 29.29 % | 0.65 % | 0.06 % |
| P22 | 16316648 | 99.30 % | 49.71 % | 31.45 % | 0.76 % | 0.08 % |
| P23 | 8774224 | 91.04 % | 43.70 % | 27.01 % | 0.52 % | 0.04 % |
| P24 | 17761720 | 92.24 % | 39.87 % | 25.64 % | 0.60 % | 0.06 % |
| P25 | 25162690 | 98.60 % | 50.13 % | 30.52 % | 0.67 % | 0.05 % |
| P26 | 28886145 | 98.96 % | 52.03 % | 31.92 % | 0.73 % | 0.06 % |
| P27 | 14478034 | 94.46 % | 46.86 % | 28.84 % | 0.63 % | 0.05 % |
| P28 | 12544372 | 90.38 % | 40.41 % | 25.42 % | 0.55 % | 0.05 % |
| P29 | 14180656 | 94.14 % | 44.02 % | 27.44 % | 0.62 % | 0.05 % |
| P30 | 11665715 | 91.10 % | 42.73 % | 27.04 % | 0.60 % | 0.05 % |
| P31 | 18834258 | 97.90 % | 49.92 % | 29.97 % | 0.65 % | 0.06 % |
| P32 | 10610062 | 94.84 % | 47.45 % | 29.50 % | 0.67 % | 0.05 % |
| P33 | 19701008 | 94.18 % | 42.33 % | 26.29 % | 0.58 % | 0.05 % |
| P34 | 8264857 | 91.67 % | 42.31 % | 26.17 % | 0.54 % | 0.04 % |
| P35 | 15030324 | 90.44 % | 41.07 % | 25.54 % | 0.53 % | 0.04 % |
| P36 | 15462190 | 91.81 % | 40.69 % | 25.31 % | 0.50 % | 0.05 % |

**Metatranscriptomics**

| Sample | Total read pairs | Total read pairs trimmed | Total read pairs rRNA filtered | Total KEGG hits | Total KEGG gene with KO | Total metabolic marker gene hits | Total SILVA SSU rRNA counts |
|---|---|---|---|---|---|---|---|
| P1 | 33751654 | 99.85 % | 4.24 % | 18.36 % | 12.24 % | 0.46 % | 35.84 % |
| P2 | 34910802 | 99.84 % | 3.87 % | 17.28 % | 11.23 % | 0.40 % | 34.65 % |
| P3 | 24329493 | 99.87 % | 4.00 % | 21.66 % | 14.21 % | 0.41 % | 34.87 % |
| P4 | 43438804 | 99.88 % | 4.17 % | 21.08 % | 13.49 % | 0.46 % | 35.16 % |
| P5 | 82436550 | 99.77 % | 4.75 % | 21.88 % | 14.13 % | 0.41 % | 35.32 % |
| P6 | 50743771 | 99.87 % | 3.98 % | 18.84 % | 12.37 % | 0.48 % | 35.89 % |
| P7 | 47603328 | 99.84 % | 5.36 % | 18.26 % | 11.57 % | 0.36 % | 34.07 % |
| P8 | 33887176 | 99.89 % | 5.13 % | 24.64 % | 16.41 % | 0.84 % | 35.08 % |
| P9 | 33545784 | 99.89 % | 4.57 % | 21.47 % | 13.93 % | 0.41 % | 34.23 % |
| P10 | 30477031 | 99.84 % | 4.31 % | 20.00 % | 12.60 % | 0.34 % | 35.12 % |
| P11 | 45862483 | 99.88 % | 5.06 % | 17.92 % | 11.39 % | 0.37 % | 34.70 % |
| P12 | 33640735 | 99.88 % | 5.04 % | 22.38 % | 14.47 % | 0.42 % | 34.63 % |
| P13 | 51056735 | 99.74 % | 4.18 % | 18.95 % | 12.46 % | 0.43 % | 36.17 % |
| P14 | 29241743 | 99.86 % | 4.79 % | 19.38 % | 12.08 % | 0.31 % | 35.70 % |
| P15 | 33944127 | 99.86 % | 5.10 % | 24.48 % | 15.74 % | 0.35 % | 34.74 % |
| P16 | 34143578 | 99.87 % | 3.88 % | 19.98 % | 12.89 % | 0.33 % | 35.49 % |
| P17 | 51385751 | 99.87 % | 3.87 % | 18.61 % | 12.18 % | 0.45 % | 36.29 % |
| P18 | 39422598 | 99.88 % | 4.63 % | 20.96 % | 13.57 % | 0.36 % | 35.23 % |
| P19 | 34140671 | 99.84 % | 4.04 % | 19.73 % | 12.59 % | 0.33 % | 34.65 % |
| P20 | 39974397 | 99.88 % | 4.23 % | 16.77 % | 10.91 % | 0.48 % | 35.44 % |
| P21 | 35926836 | 99.32 % | 4.01 % | 18.03 % | 11.89 % | 0.39 % | 36.06 % |
| P22 | 26800494 | 99.88 % | 4.56 % | 18.01 % | 11.95 % | 0.56 % | 34.67 % |
| P23 | 72336681 | 99.78 % | 5.02 % | 19.79 % | 12.27 % | 0.25 % | 33.59 % |
| P24 | 35206478 | 99.87 % | 4.67 % | 17.81 % | 11.93 % | 0.57 % | 33.86 % |
| P25 | 41719853 | 99.81 % | 4.38 % | 17.01 % | 10.86 % | 0.37 % | 34.36 % |
| P26 | 31456366 | 99.88 % | 4.00 % | 18.51 % | 11.72 % | 0.47 % | 35.71 % |
| P27 | 32189269 | 99.84 % | 4.22 % | 20.07 % | 12.96 % | 0.38 % | 35.37 % |
| P28 | 29075206 | 99.81 % | 5.22 % | 22.39 % | 14.08 % | 0.34 % | 35.07 % |
| P29 | 34332202 | 99.89 % | 4.81 % | 21.56 % | 14.03 % | 0.48 % | 34.08 % |
| P30 | 26487311 | 99.87 % | 4.42 % | 23.56 % | 15.22 % | 0.38 % | 35.55 % |
| P31 | 27396219 | 99.87 % | 4.35 % | 20.99 % | 12.87 % | 0.38 % | 36.01 % |
| P32 | 34915485 | 99.88 % | 4.31 % | 19.59 % | 12.82 % | 0.46 % | 36.39 % |
| P33 | 25802311 | 99.88 % | 3.67 % | 18.18 % | 11.68 % | 0.40 % | 36.47 % |
| P34 | 27390388 | 99.84 % | 4.59 % | 23.77 % | 15.12 % | 0.34 % | 34.83 % |
| P35 | 29249396 | 99.85 % | 4.52 % | 20.86 % | 13.01 % | 0.31 % | 34.94 % |
| P36 | 30457805 | 99.83 % | 4.93 % | 20.09 % | 12.76 % | 0.32 % | 33.81 % |

**Supplementary Table 24**. The selected set of interesting core metabolic Kyoto Encyclopedia of Genes and Genomes (KEGG) modules investigated in this study.

| **Selected core KEGG modules** |
|---|
| M00001 Glycolysis (Embden-Meyerhof pathway), glucose => pyruvate |
| M00002 Glycolysis, core module involving three-carbon compounds |
| M00003 Gluconeogenesis, oxaloacetate => fructose-6P |
| M00307 Pyruvate oxidation, pyruvate => acetyl-CoA |
| M00009 Citrate cycle (TCA cycle, Krebs cycle) |
| M00010 Citrate cycle, first carbon oxidation, oxaloacetate => 2-oxoglutarate |
| M00011 Citrate cycle, second carbon oxidation, 2-oxoglutarate => oxaloacetate |
| M00004 Pentose phosphate pathway (Pentose phosphate cycle) |
| M00006 Pentose phosphate pathway, oxidative phase, glucose 6P => ribulose 5P |
| M00007 Pentose phosphate pathway, non-oxidative phase, fructose 6P => ribose 5P |
| M00580 Pentose phosphate pathway, archaea, fructose 6P => ribose 5P |
| M00005 PRPP biosynthesis, ribose 5P => PRPP |
| M00008 Entner-Doudoroff pathway, glucose-6P => glyceraldehyde-3P + pyruvate |
| M00308 Semi-phosphorylative Entner-Doudoroff pathway, gluconate => glycerate-3P |
| M00633 Semi-phosphorylative Entner-Doudoroff pathway, gluconate/galactonate => glycerate-3P |
| M00309 Non-phosphorylative Entner-Doudoroff pathway, gluconate/galactonate => glycerate |
| M00165 Reductive pentose phosphate cycle (Calvin cycle) |
| M00168 CAM (Crassulacean acid metabolism), dark |
| M00169 CAM (Crassulacean acid metabolism), light |
| M00172 C4-dicarboxylic acid cycle, NADP - malic enzyme type |
| M00171 C4-dicarboxylic acid cycle, NAD - malic enzyme type |
| M00170 C4-dicarboxylic acid cycle, phosphoenolpyruvate carboxykinase type |
| M00173 Reductive citrate cycle (Arnon-Buchanan cycle) |
| M00376 3-Hydroxypropionate bi-cycle |
| M00375 Hydroxypropionate-hydroxybutylate cycle |
| M00374 Dicarboxylate-hydroxybutyrate cycle |
| M00377 Reductive acetyl-CoA pathway (Wood-Ljungdahl pathway) |
| M00579 Phosphate acetyltransferase-acetate kinase pathway, acetyl-CoA => acetate |
| M00620 Incomplete reductive citrate cycle, acetyl-CoA => oxoglutarate |
| M00567 Methanogenesis, CO2 => methane |
| M00357 Methanogenesis, acetate => methane |
| M00356 Methanogenesis, methanol => methane |
| M00563 Methanogenesis, methylamine/dimethylamine/trimethylamine => methane |
| M00358 Coenzyme M biosynthesis |
| M00608 2-Oxocarboxylic acid chain extension, 2-oxoglutarate => 2-oxoadipate => 2-oxopimelate => 2-oxosuberate |
| M00174 Methane oxidation, methanotroph, methane => formaldehyde |
| M00346 Formaldehyde assimilation, serine pathway |
| M00345 Formaldehyde assimilation, ribulose monophosphate pathway |
| M00344 Formaldehyde assimilation, xylulose monophosphate pathway |
| M00378 F420 biosynthesis, archaea |
| M00935 Methanofuran biosynthesis |
| M00422 Acetyl-CoA pathway, CO2 => acetyl-CoA |
| M00175 Nitrogen fixation, nitrogen => ammonia |
| M00531 Assimilatory nitrate reduction, nitrate => ammonia |
| M00530 Dissimilatory nitrate reduction, nitrate => ammonia |
| M00529 Denitrification, nitrate => nitrogen |
| M00528 Nitrification, ammonia => nitrite |
| M00804 Complete nitrification, comammox, ammonia => nitrite => nitrate |
| M00973 Anammox, nitrite + ammonia => nitrogen |
| M00176 Assimilatory sulfate reduction, sulfate => H2S |
| M00596 Dissimilatory sulfate reduction, sulfate => H2S |
| M00595 Thiosulfate oxidation by SOX complex, thiosulfate => sulfate |
