## Supplementary Figures for "Soil microbiome structure and function reflect environmental variation rather than reindeer presence in a northern peatland"

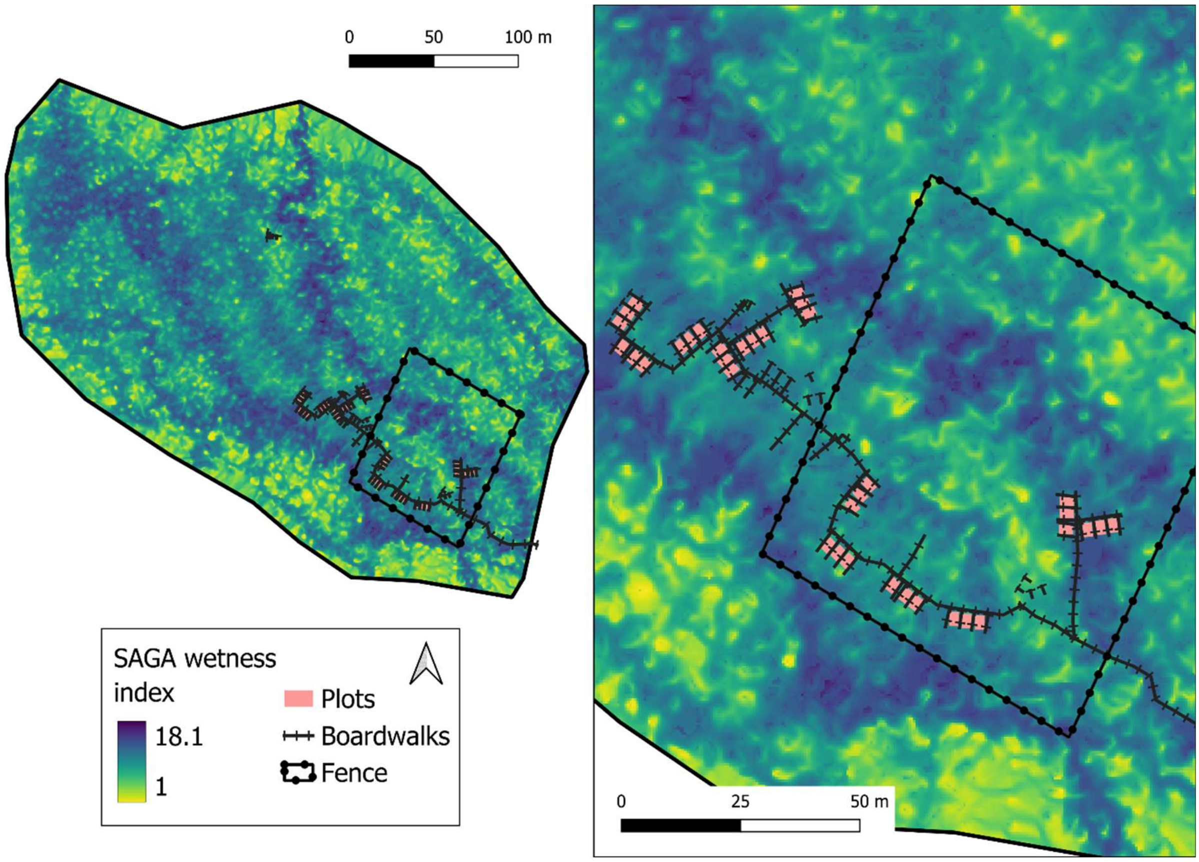


**Supplementary Figure 1.** SAGA Wetness index for Puukkosuo rich fen and, in a close-up, for the experimental plots showing positions relative to the exclusion fence and boardwalk. Values of SAGA wetness index can be used as an index of surface water flow paths with greater values corresponding to wetter conditions.

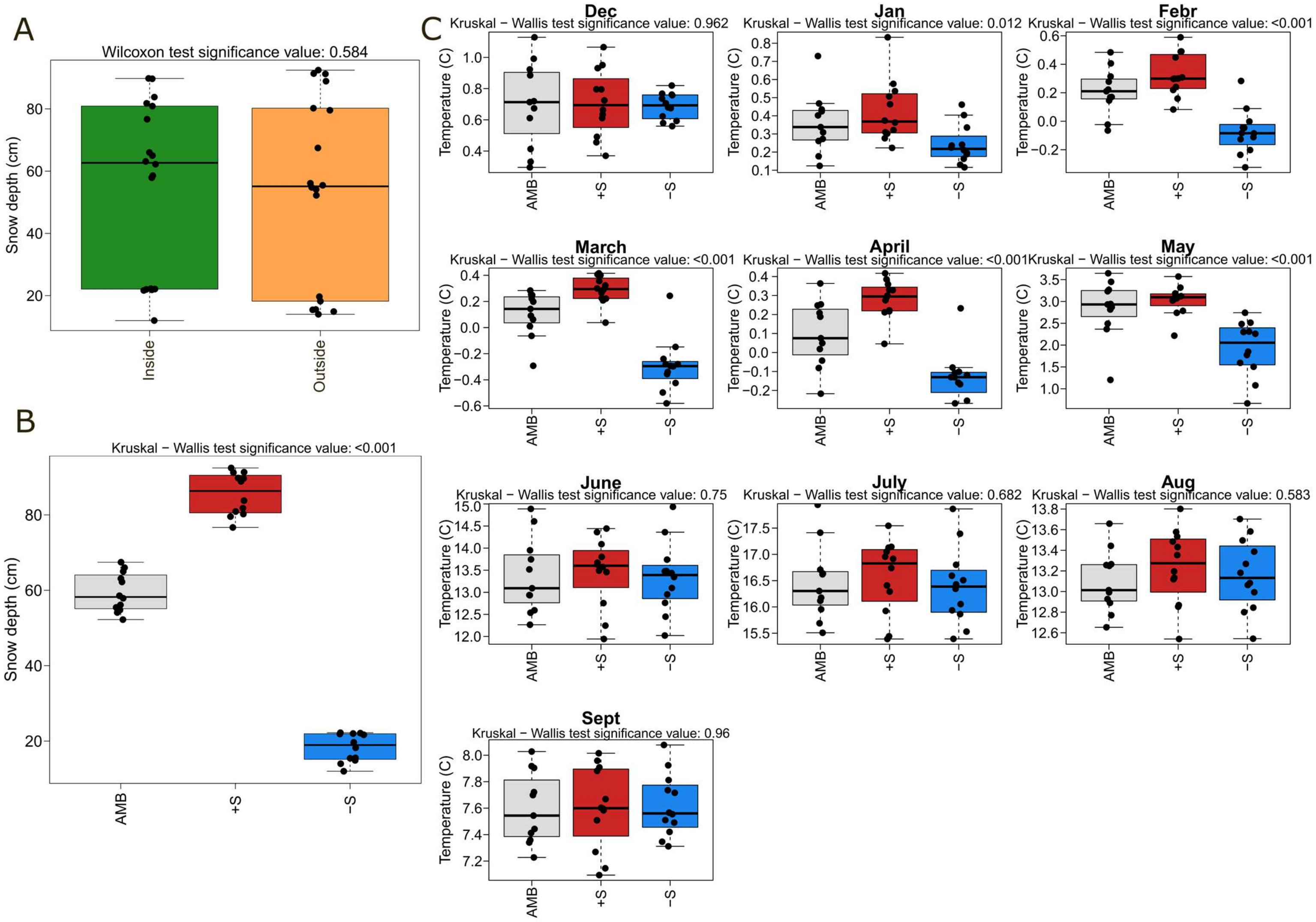


**Supplementary Figure 2.** Snow depths and soil temperatures in Puukkosuo during the winter 2020-2021 and the growing season 2021. A) Average snow depths by the exclusion treatment during the maximum snow depth time in March 2021. The Wilcoxon signed rank test was used to test whether snow depths in plots outside and inside the exclosure differed significantly. B) Average snow depths by the snow manipulation treatments during the maximum snow depth time in March 2021. Kruskal-Wallis's test was used to test whether snow depths differed significantly between the snow treatments. At each plot, snow depth was measured at three spots to calculate average depth per plot. C) Average soil temperatures by the snow treatments from December 2020 to September 2021. Kruskal-Wallis's test was used to test whether soil temperatures differed significantly between the snow treatments. Significance values in all plots refer to p-values from the used statistical tests. Soil temperature was measured at 5 cm depth using Campbell Scientific T107 temperature probes and the temperature data were recorded at 10-minute intervals using Campbell Scientific CR1000X measurement and control dataloggers. These data were used to calculate monthly mean soil temperatures for each. Months from December to May were the months when the snow treatments were carried out and months from June to September were without snow.

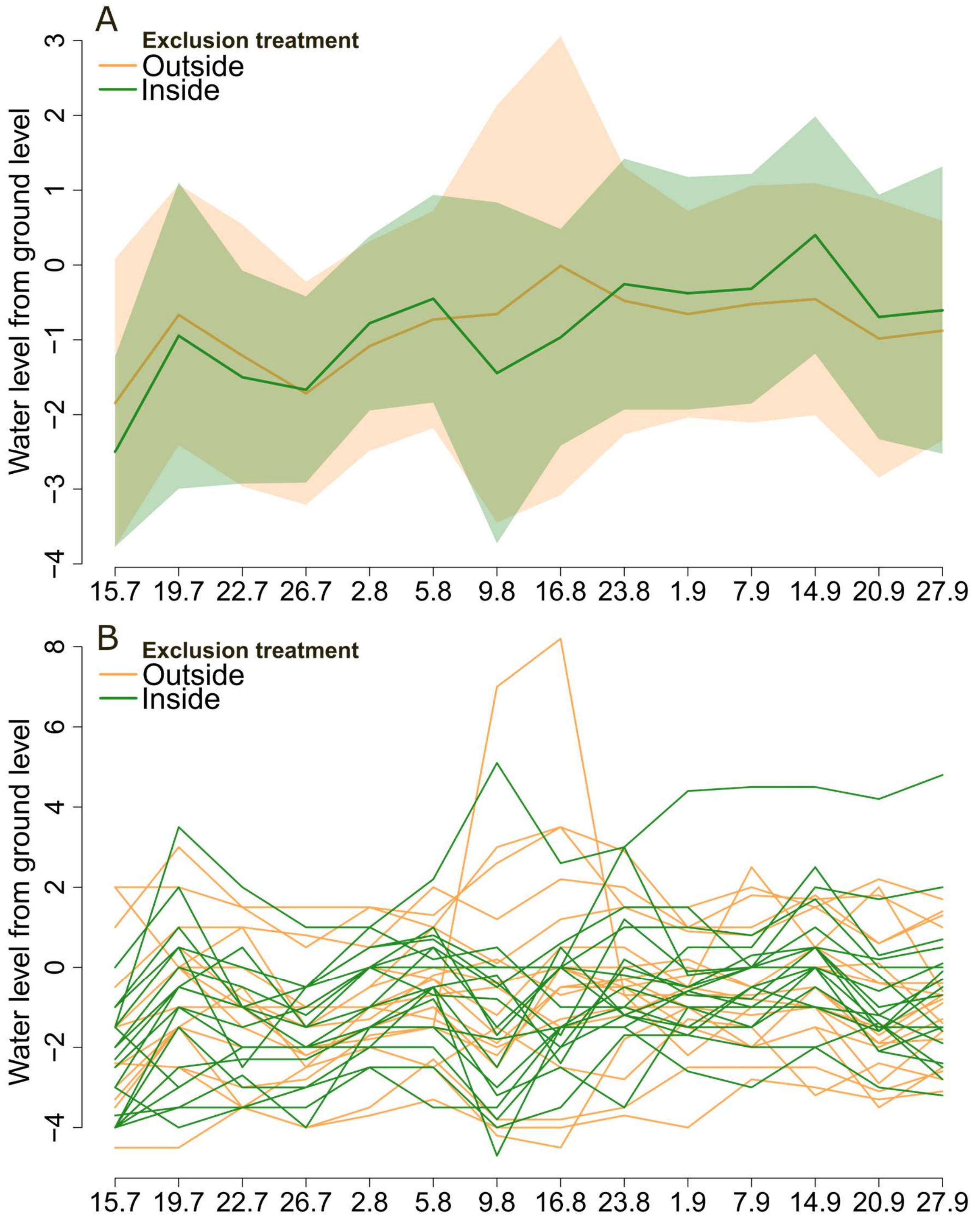


**Supplementary Figure 3**. Water table levels in Puukkosuo during Summer 2021 (from mid-July to September) by the exclusion treatment. A) Mean water table levels from the ground level outside and inside the exclosure. The colored lines display mean water table levels over all the plots in the treatment while the shaded area represents standard deviation. B) Water table levels from the ground level outside and inside the exclosure shown indivually for all the sample plots.

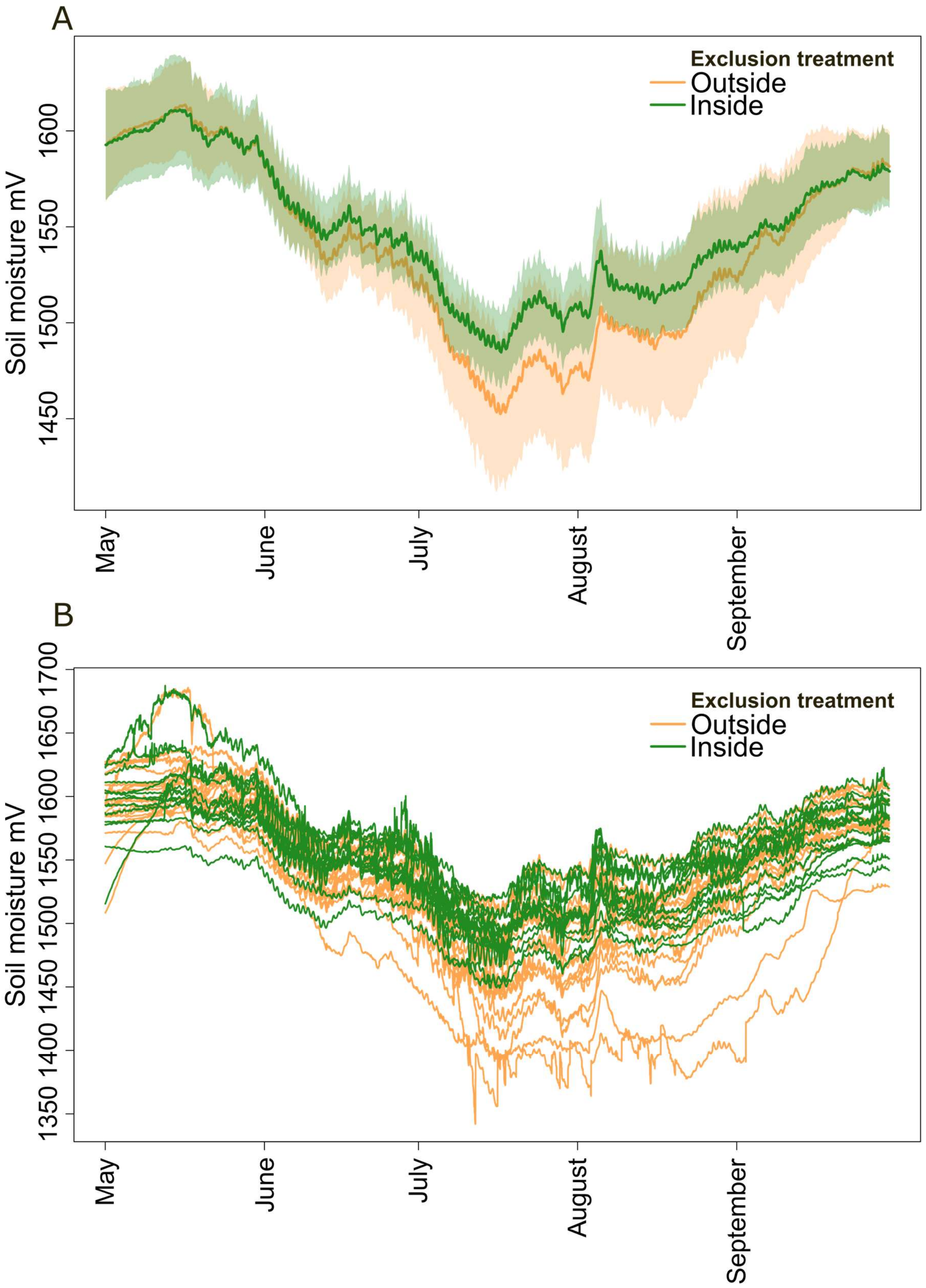


**Supplementary Figure 4.** Soil moisture logger data in Puukkosuo during the snow free period of 2021 by the exclusion treatment. A) Mean soil moisture logger data outside and inside the exclosure. The colored lines display mean values over all the plots in the treatment while the shaded area represents standard deviation. B) Soil moisture logger data outside and inside the exclosure shown individually for all the sample plots. The soil moisture was measured at a depth of 5 cm using Delta-T SM150T soil moisture sensors and sensor data were recorded and stored using Campbell Scientific CR1000X measurement and control dataloggers at 10-minute intervals between 1 May to 30 September 2021. The mV values are not calibrated for the specific soil type but are comparable between plots and between areas outside and inside the exclosure. Higher values indicate higher moisture. Measurements for four plots with sensor/logger malfunctions inside the exclosure are excluded from the visualization.

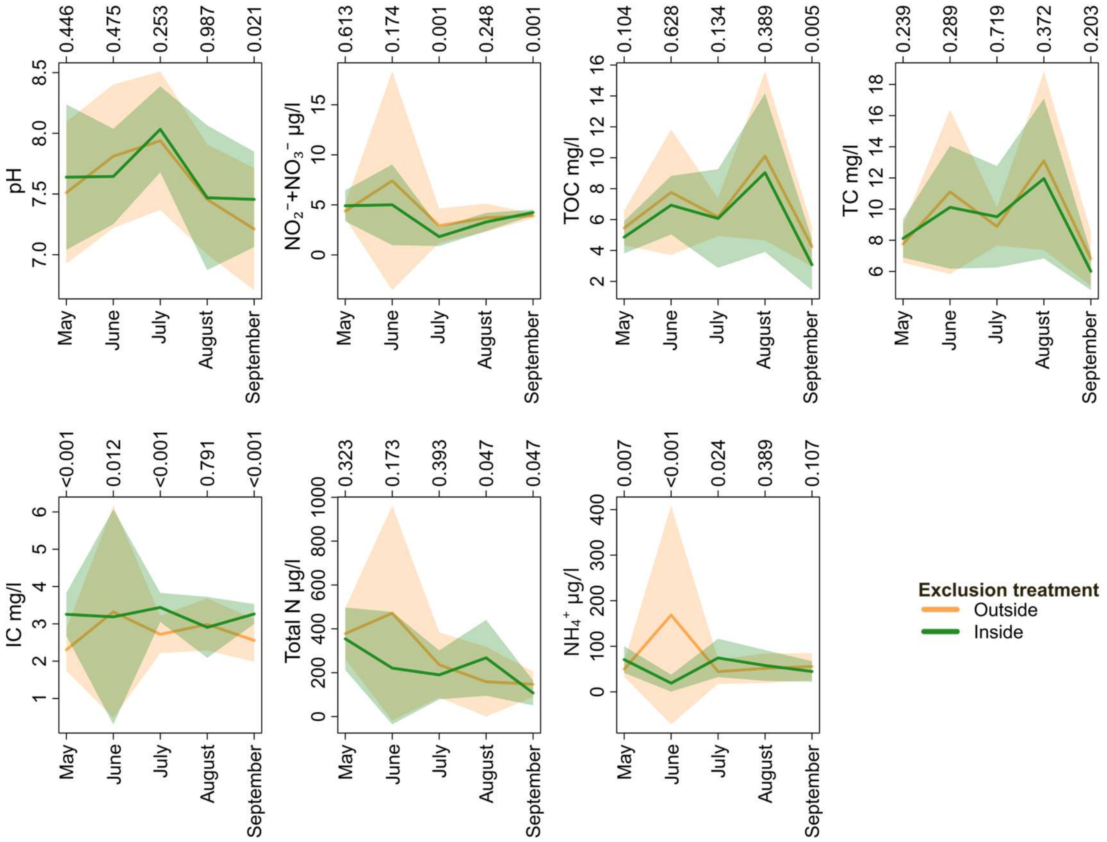


**Supplementary Figure 5.** Mean values for the measured pore water variables during the snow free period 2021 by the exclusion treatment. The colored lines display mean values over all the plots in the treatment while the shaded area represents standard deviation. The Wilcoxon signed rank test was used to test whether the pore water variable values differ significantly between outside and inside the exclosure for each month and significance values (p-values) are displayed on the top axis. TC refers to total dissolved carbon, TOC to dissolved organic carbon and IC to dissolved inorganic carbon. Peat pore-water was collected five times from May to September 2021 using Rhizosphere Rhizon samplers at 10 cm depth into evacuated opaque syringes, filtered with Sarstedt 0.45 µm sterile nylon and frozen at −18 °C. Thawed samples were analyzed for pH using Metrohm 913 pH/DO Meter, for TOC and IC with Shimadzu TOC-L CPN and for Total N, $NH_4^+$ and $NO_2^-$ + $NO_3^-$ with Seal Analytical AA500.

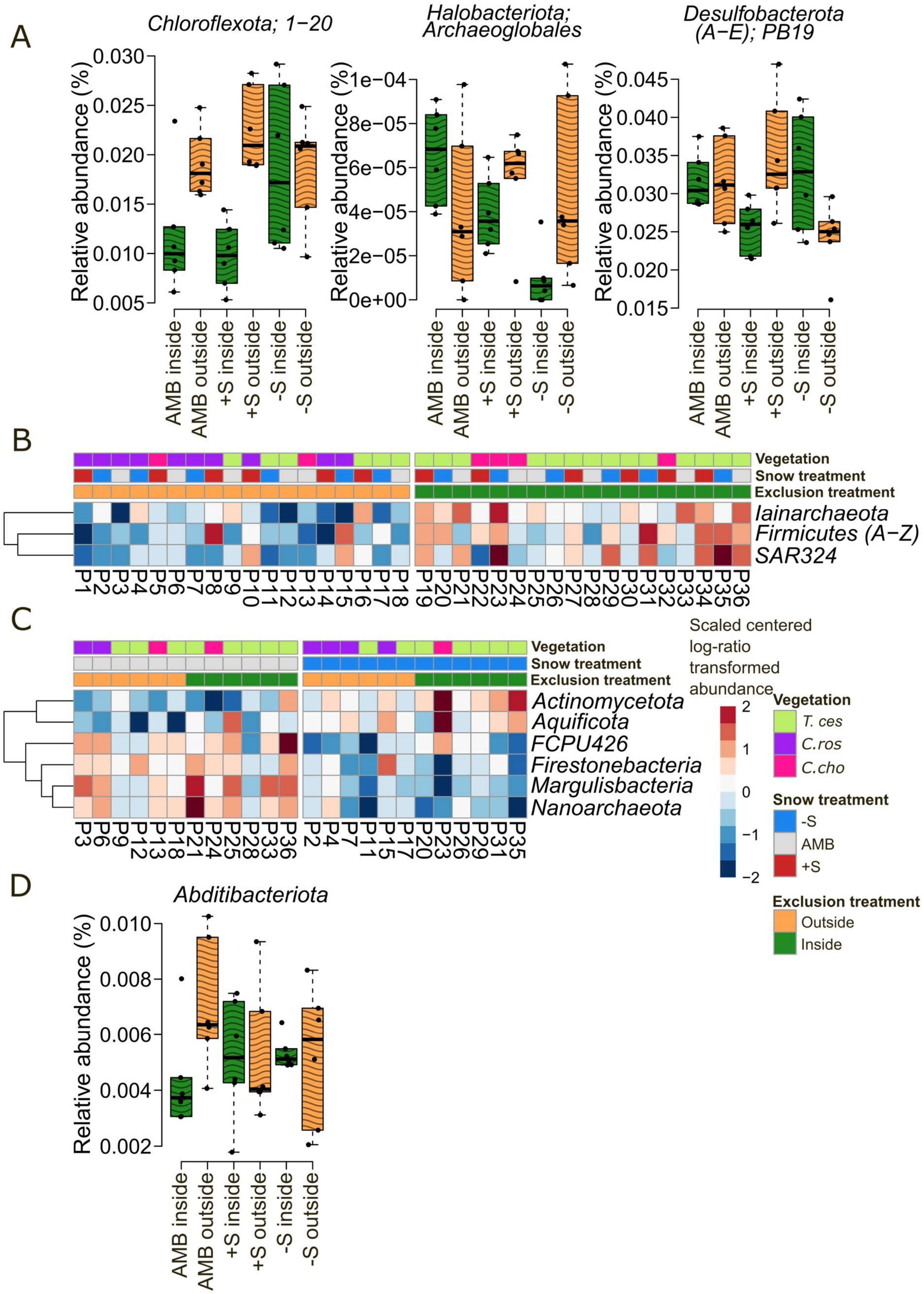


**Supplementary Figure 6.** Differentially expressed taxa in metatranscriptomic data. A) Orders with significant exclusion treatment and snow treatment interactions. B) Phyla differentially expressed between plots outside and inside the exclosure. C) Phyla differentially expressed between snow removal and snow control plots. D) Phyla with significant exclusion treatment and snow treatment interactions. Linear mixed-effects models (exclusion treatment and snow treatment as fixed effects, vegetation cluster as a random effect) were used to identify differentially expressed taxa (false discovery rate ≤ 0.1) from center-log-ratio (CLR) transformed data. For the heatmaps in B and C, the CLR abundances are scaled and clustered using hierarchical clustering with Euclidean distance.

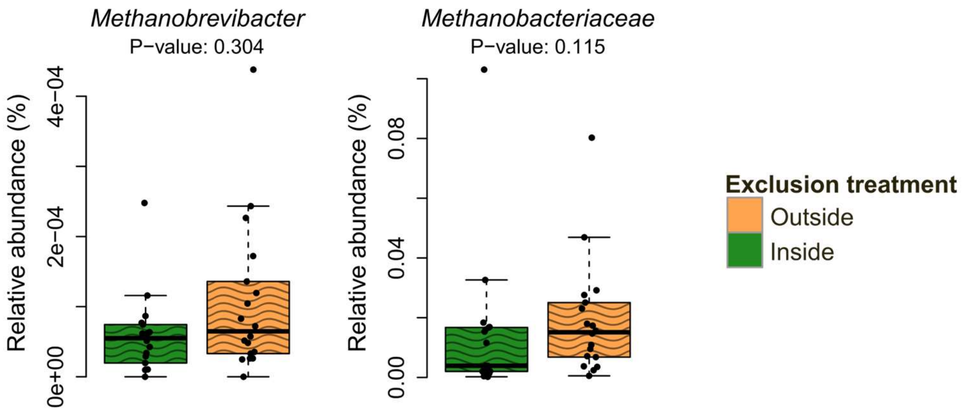


**Supplementary Figure 7.** The relative expression of reindeer Rumen associated genus *Methanobrevibacter* and the parent family *Methanobacteriaceae* outside and inside the exclosure in the metatranscriptomics data.

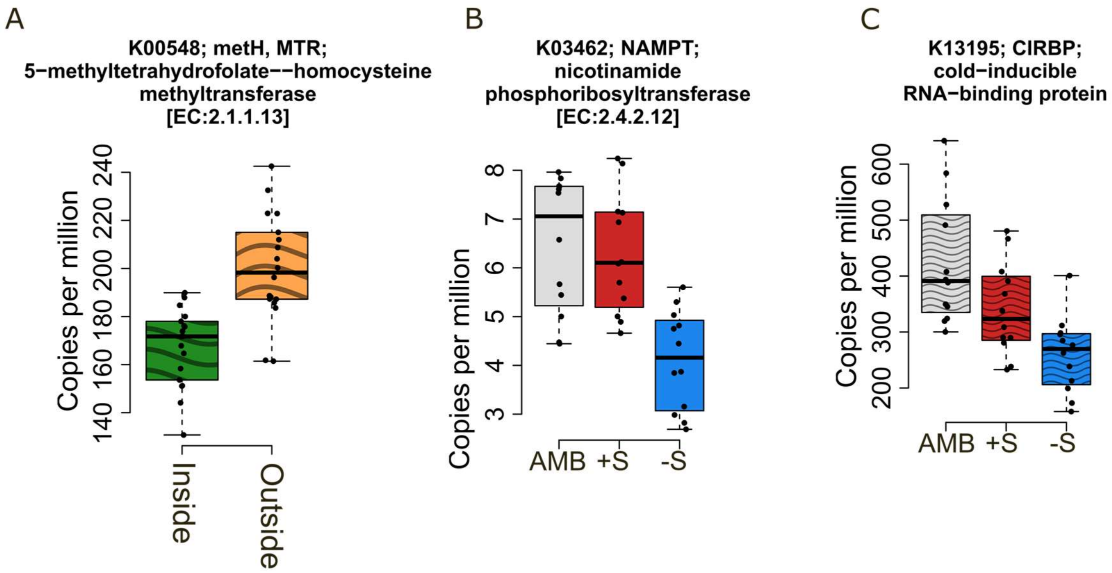


**Supplementary Figure 8.** A) Differentially expressed Kyoto Encyclopedia of Genes and Genomes (KEGG) orthologs (KOs) associated with the exclusion treatment in the metatranscriptomics data. B) Differentially abundant KOs associated with the snow removal treatment in the metagenomics data and C) Differentially expressed KOs associated with the snow removal treatment in the metatranscriptomics data.

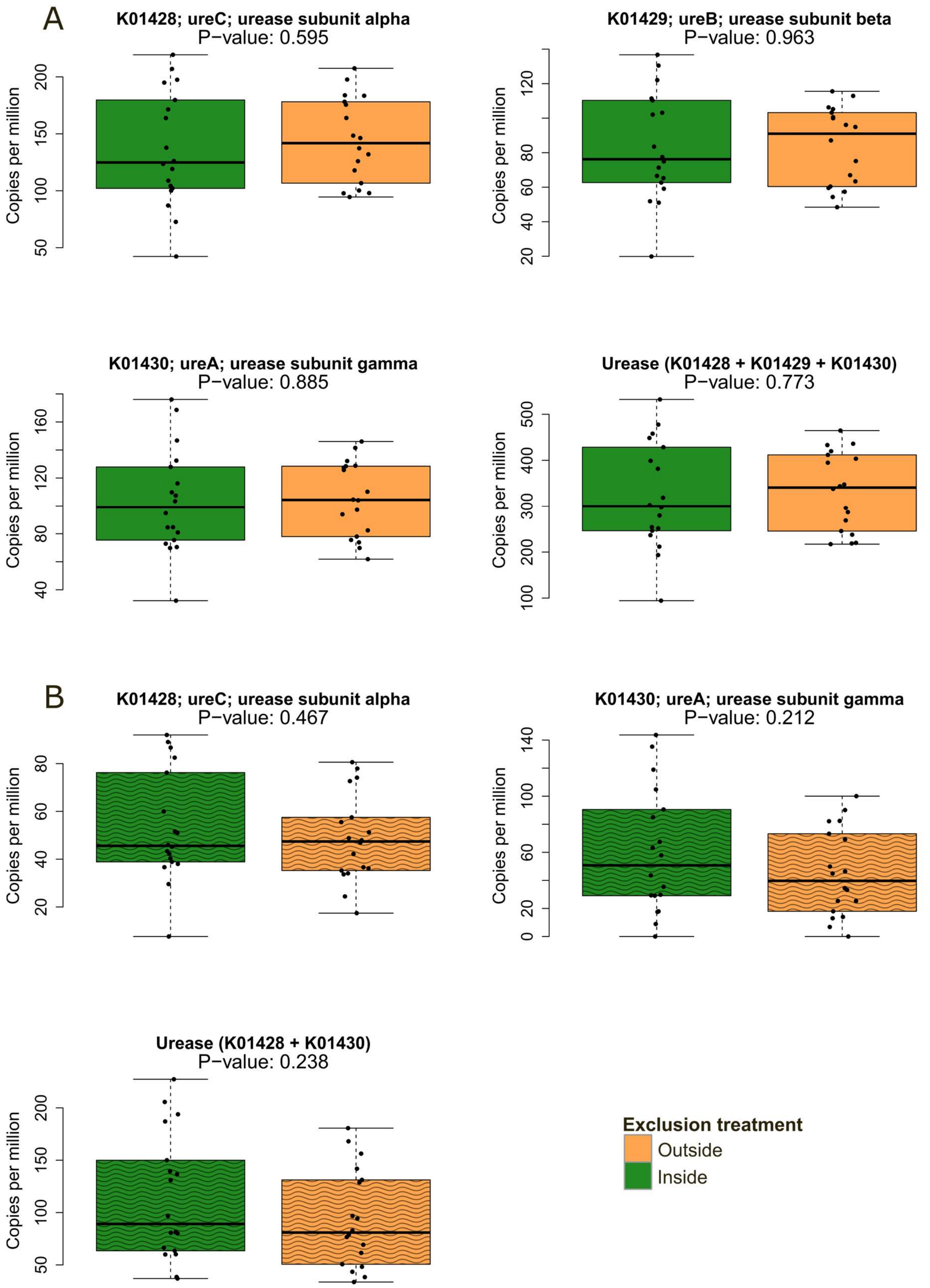


**Supplementary Figure 9.** The relative abundance and expression of the detected Kyoto Encyclopedia of Genes and Genomes (KEGG) orthologs (KOs) for the urease-enzyme subunits and the aggregated urease-enzyme by the exclusion treatment in the A) metagenomics data and B) metatranscriptomics data.

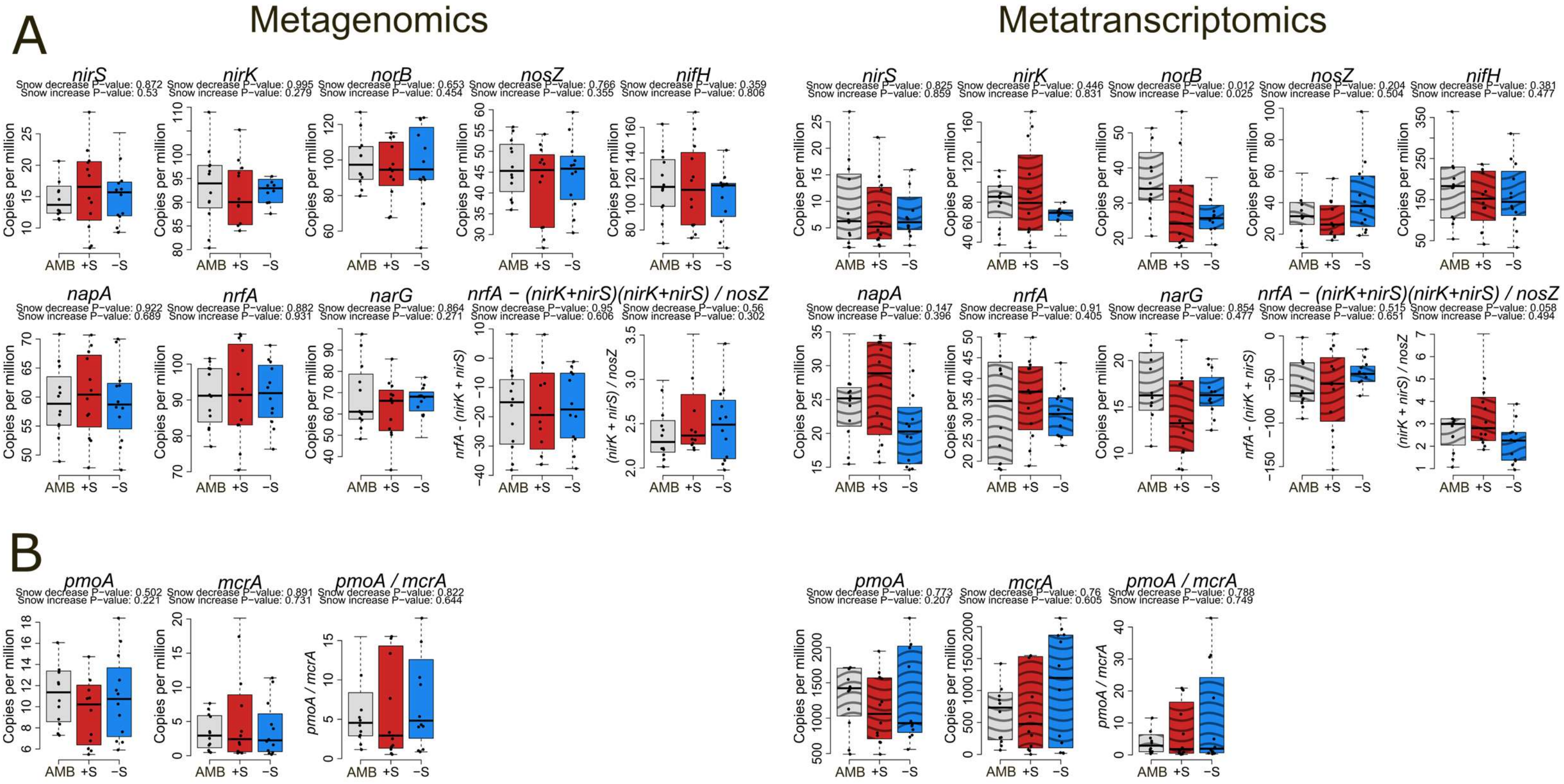


**Supplementary Figure 10.** Relative abundance and expression of nitrogen- and methane-cycling marker genes and ratios under the snow treatments in the metagenomics and metatranscriptomics data. A) Nitrogen-cycling genes and ratios. B) Methane-cycling genes and ratios. Metagenomic (MG) and metatranscriptomic (MT) reads were aligned to the eukaryote-filtered Greening Lab metabolic marker database and summarized to marker-gene level. All marker gene counts were converted to copies-per-million (as for transcripts-per-million). Linear mixed-effects models (LMMs) (exclusion treatment and snow treatment as fixed effects, vegetation cluster as a random effect) were used to determine significance of the MG abundance and MT expression association to treatments. For the LMMs the marker gene abundance and expression were $log_2$-transformed. Boxplots are plotted without potential outlier observations for better visualization.

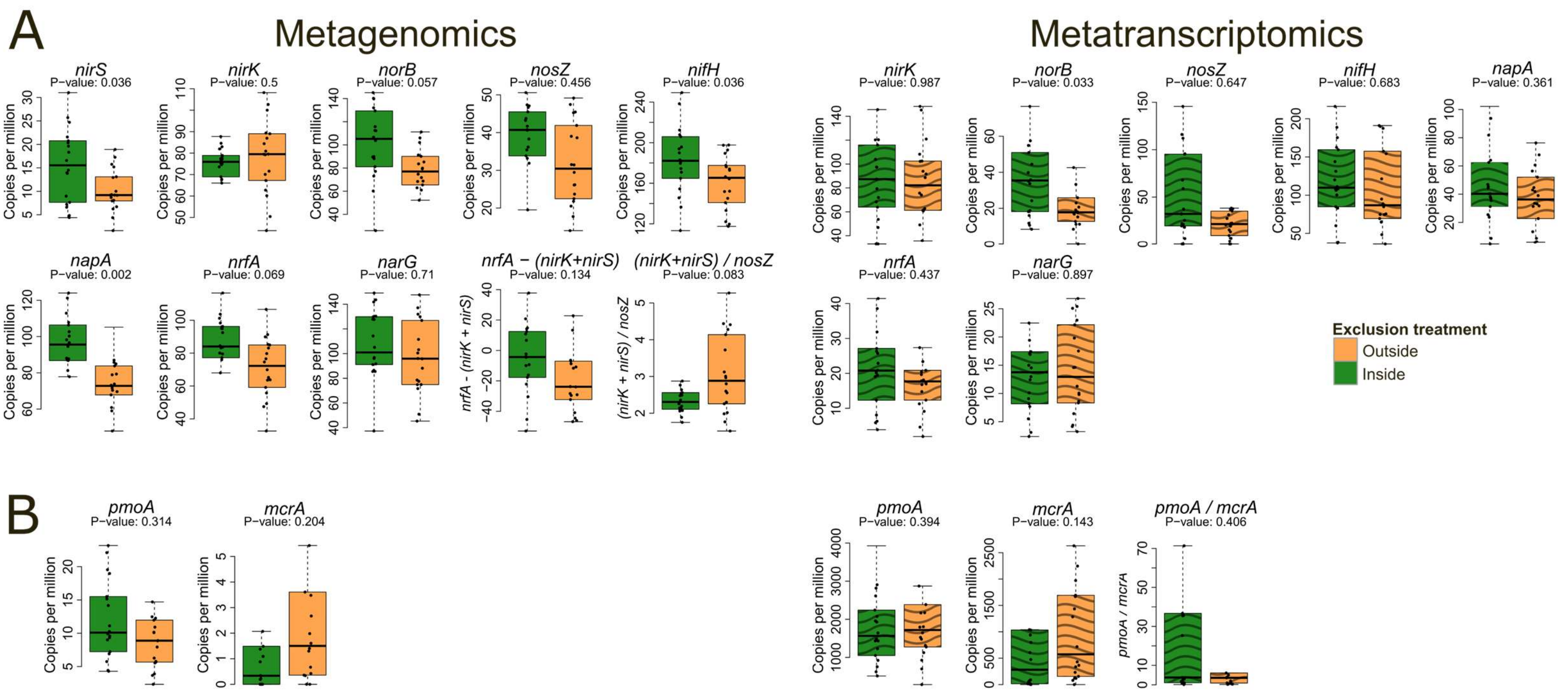


**Supplementary Figure 11.** Relative abundance and expression of nitrogen- and methane-cycling marker genes and ratios in relation to exclusion treatment in the contig-based analyses in metagenomics and metatranscriptomics data. A) Nitrogen-cycling genes and ratios B) Methane-cycling genes and ratios. Contigs were assembled with MEGAHIT and genes predicted with Prodigal in anvi'o. The contig genes were aligned to the eukaryote-filtered Greening Lab metabolic marker database and the metagenomic (MG) and metatranscriptomic (MT) reads were mapped back to the contig genes and summarized to marker-gene level. All marker gene counts were converted to copies-per-million (as for transcripts-per-million). Linear mixed-effects models (LMMs) (exclusion treatment and snow treatment as fixed effects, vegetation cluster as a random effect) were used to determine significance of the MG abundance and MT expression association to treatments. For the LMMs the marker gene abundance and expression were $log_2$-transformed. Boxplots are plotted without potential outlier observations for better visualization.

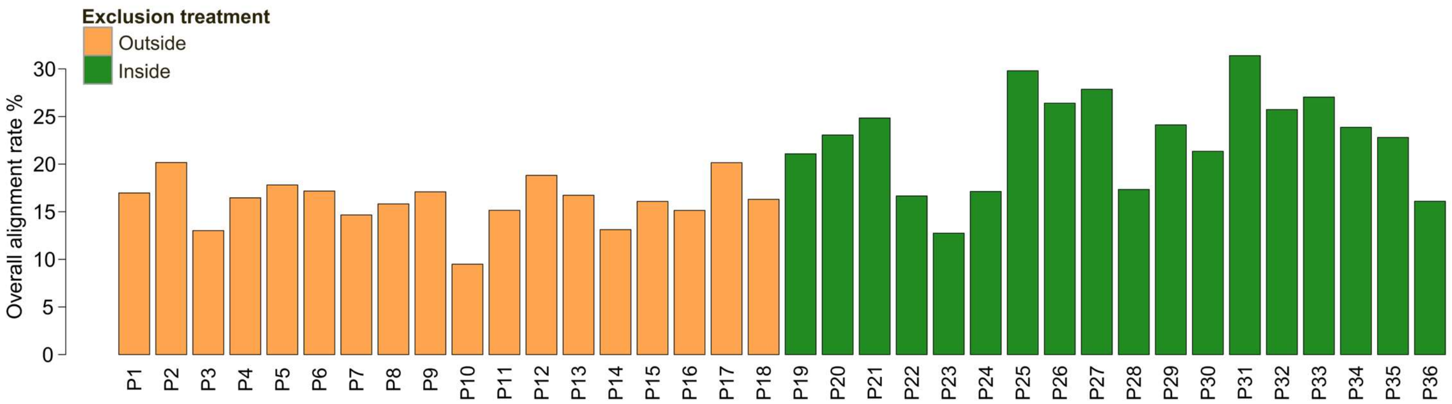


**Supplementary Figure 12.** Overall alignment rates of metagenomic reads to the MEGAHIT co-assemblies. Each sample was mapped with Bowtie2 to its corresponding co-assembly from outside or inside the exclosure.

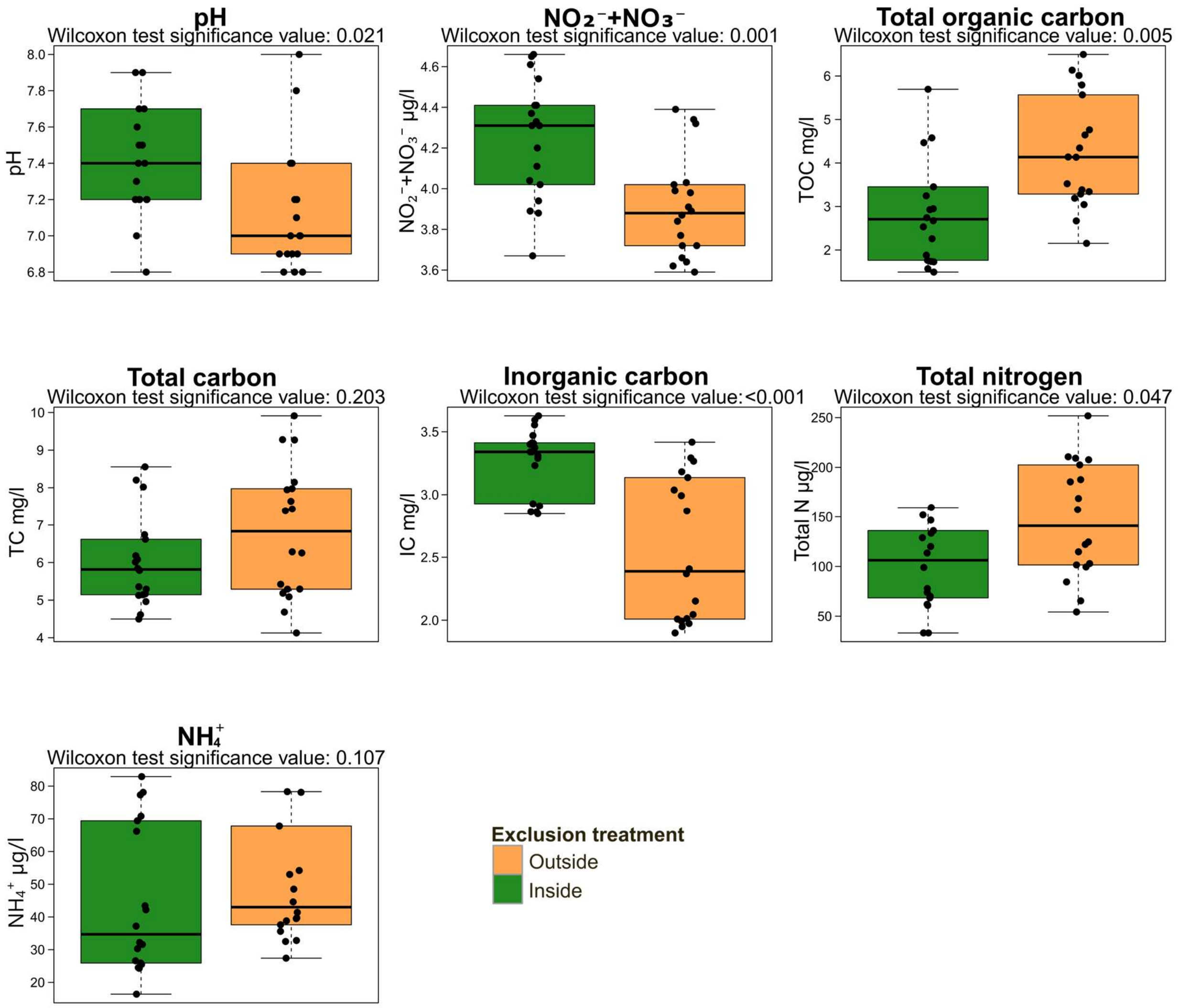


**Supplementary Figure 13.** Puukkosuo pore water variables measured in September 2021 by the exclusion treatment. The Wilcoxon signed rank test was used to test whether the pore water variable values differed significantly between areas outside and inside the exclosure. Significance values in all plots refer to p-values from the used statistical test.

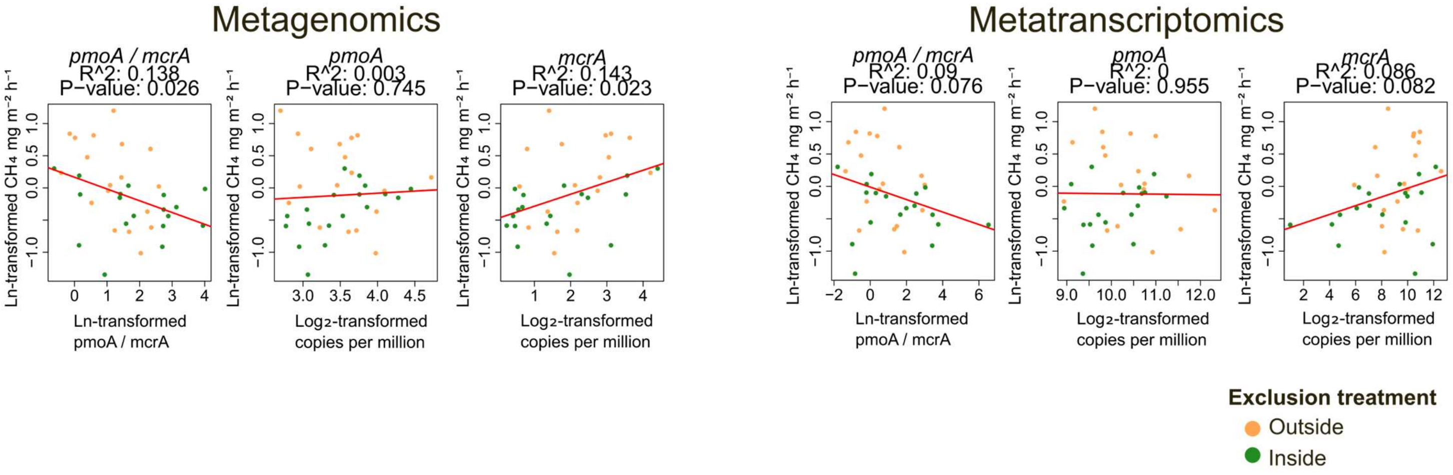


**Supplementary Figure 14.** Associations of methane fluxes (measured 20.9.2021) with methane-cycling genes and the methane oxidation / methanogenesis (*pmoA* / *mcrA*) ratio in the metagenomics and metatranscriptomics data. Fluxes and gene ratios were natural-log transformed; gene abundances and expression were $\log_2$-transformed. Metagenomic and metatranscriptomic reads were aligned to the eukaryote-filtered Greening Lab metabolic marker database, summarized to marker-gene level and the marker gene counts were converted to copies-per-million (as for transcripts-per-million).

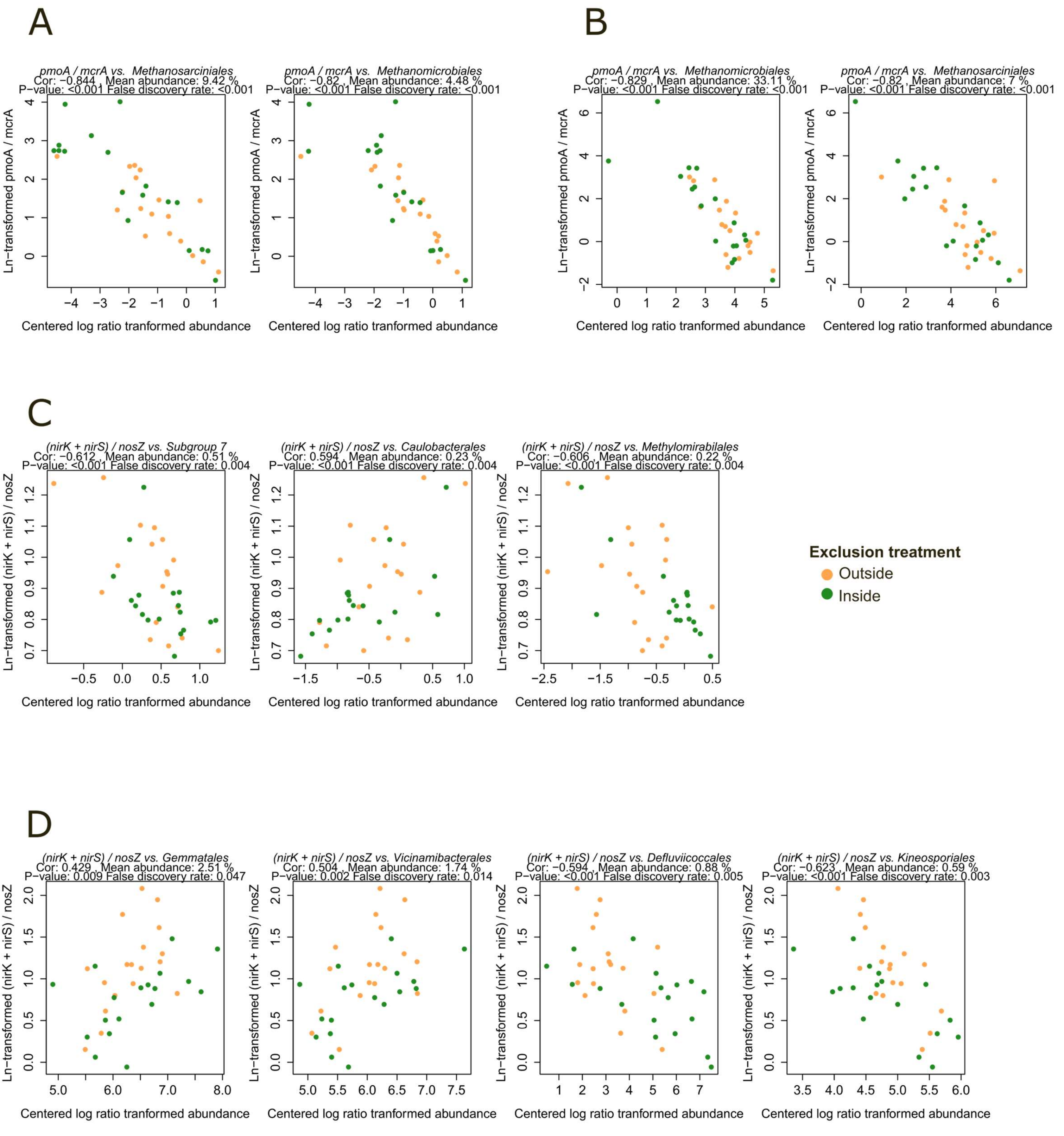


**Supplementary Figure 15.** Examples of correlations between order-level taxa and metabolic marker gene ratios in metagenomics and metatranscriptomics data. A) Taxa vs. methane oxidation / methanogenesis (*pmoA* / *mcrA*) in metagenomes. B) Taxa vs. *pmoA* / *mcrA* in metatranscriptomes. C) Taxa vs. (*nirS* + *nirK*) / *nosZ* in metagenomes. D) Taxa vs. (*nirS* + *nirK*) / *nosZ* in metatranscriptomes. Pearson correlations were computed on CLR-transformed taxon data and natural-log transformed gene ratios. Only significant correlations (Benjamini–Hochberg false discovery rate ≤ 0.1) for the selected most abundant/expressed taxa are shown.

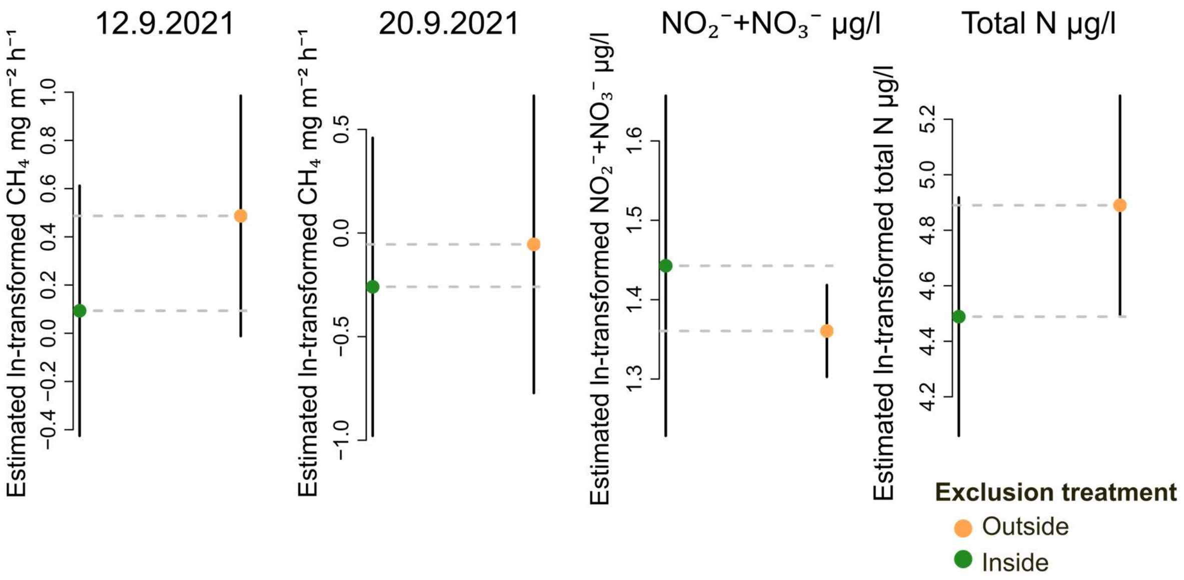


**Supplementary Figure 16.** Estimated marginal means (natural-log scale) from the linear mixed-effects models (LMMs) for methane fluxes and pore-water nitrate + nitrite ($NO_3^-$ + $NO_2^-$) and total nitrogen for the levels of the exclusion treatment. Models include reindeer exclusion treatment and snow treatment as fixed effects and vegetation cluster as a random effect.

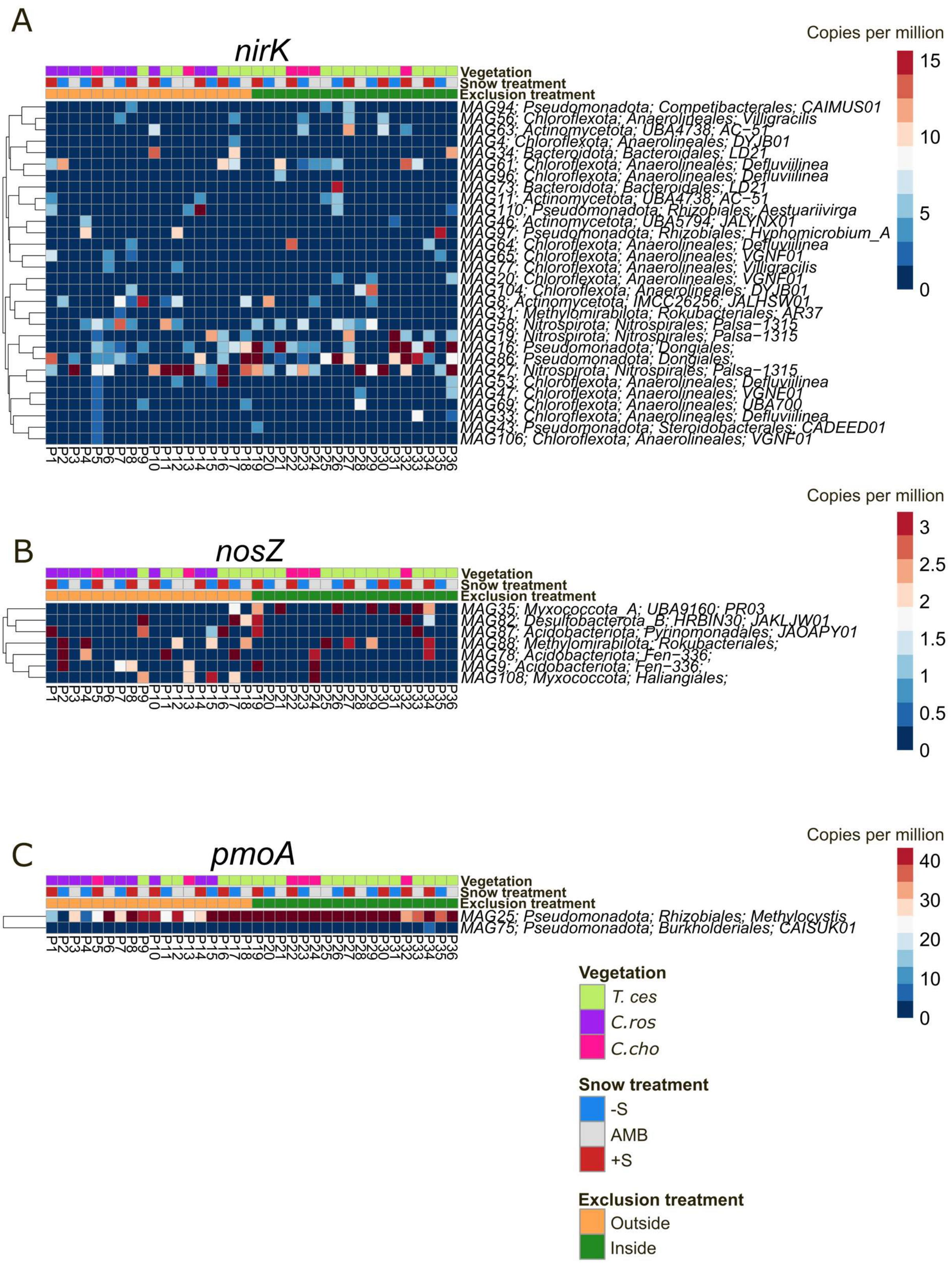


**Supplementary Figure 17.** Metagenome assembled genome (MAG)-level expression patterns for denitrification and methane oxidation related genes in the metatranscriptomics data. Expression patterns for A) *nirK*, B) *nosZ* and C) *pmoA*. Metagenomic reads were coassembled separately for the sample plots outside and inside the exclosure with MEGAHIT, processed in anvi'o, and binned with MetaBAT2, followed by manual refinement to MIMAG standards (≥ 50 % completeness, < 10 % redundancy). MAGs were dereplicated using FastANI, and taxonomy was assigned with GTDB-Tk. MAG MT genes were aligned to the eukaryote-filtered Greening Lab metabolic marker database and summarized to marker-gene level. Transcript counts were generated with featureCounts (Liao, Smyth, and Shi 2014) (Subread v2.0.6). Marker gene counts were converted to copies-per-million (as for transcripts-per-million). For all heatmaps, MAGs are clustered by hierarchical clustering with correlation distance.

Paired-end KEGG Ortholog (KO) assignment workflow

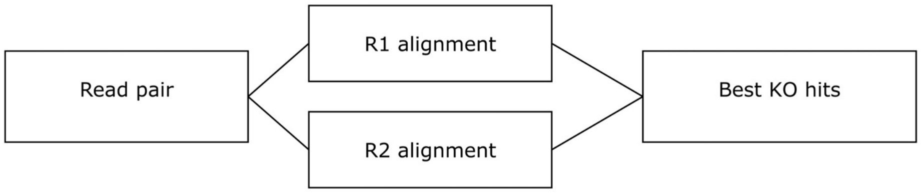


| Case | R1 best alignment | R2 best alignment | Assigned KO | Retained | Rationale |
|---|---|---|---|---|---|
| 1 | KO1234 | KO1234 | KO1234 | Yes | Both reads support the same KO |
| 2 | KO1234 | none | KO1234 | Yes | One read supports KO assignment |
| 3 | none | KO1234 | KO1234 | Yes | One read supports KO assignment |
| 4 | KO1234 (bitscore 60) | KO5678 (bitscore 44) | KO1234 | Yes | Reads support different KOs; KO assigned based on clearly higher alignment score for R1 |
| 5 | KO1234 (bitscore 44) | KO5678 (bitscore 60) | KO5678 | Yes | Reads support different KOs; KO assigned based on clearly higher alignment score for R2 |
| 6 | KO1234, KO5678 (bitscore 44) | KO5678 (bitscore 52) | KO5678 | Yes | Reads support partially same KOs; no clear bitscore difference, KO assignment based on only the shared KOs |
| 7 | KO1234 (bitscore 44) | KO5678 (bitscore 52) | ambiguous | No | Reads support different KOs; no clear bitscore difference, read pair discarded as ambiguous |

**Supplementary Figure 18**. Schematic overview of the paired-end read assignment workflow for KEGG-based functional annotation. Forward (R1) and reverse (R2) reads were aligned independently to the KEGG prokaryote database using DIAMOND. Alignments from both mates were then combined using a hierarchical filtering approach to assign KEGG Orthologs (KOs). In this approach, KO assignments were retained when 1) both reads supported the same KO, 2) when only one read produced a valid hit with a KO annotation, or 3) when reads aligned to different targets, based on the higher alignment bitscore if the bitscore difference was large enough (>|15|). Read pairs with conflicting or ambiguous annotations were conservatively discarded.
